# Genetic correlation-guided mega-analysis of DO mice provides mechanistic insight and candidate genes for age-related pathologies

**DOI:** 10.1101/2025.05.13.653808

**Authors:** Martin N. Mullis, Austin E.Y.T. Lefebvre, Kathyayini Sivasubramanian, Angela Luo, Florian Schmid, Matt Sooknah, Kevin M. Wright, Anil Raj, José Zavala-Solorio, Chunlian Zhang, Johannes Riegler, Astrid Gillich, J. Graham Ruby

## Abstract

Diversity Outbred (DO) mice are a powerful model system for mapping complex traits due to their high genetic diversity and mapping resolution. However, while there are extensive tools available for standard genetic analysis in DO mice, fewer techniques have been implemented to facilitate integrated, cross-study analysis. Here, we implement Haseman-Elston regression to estimate genetic correlations among 7,233 phenotypes measured across eleven independent DO mouse studies. We used this network of genetic correlations to cluster phenotypes according to shared genetics, which enhanced the power to detect quantitative trait loci (QTL). This approach empowered the detection of 884 QTL for 383 meta-phenotypes, explaining an average of 40.36% of the total genetic variance per mega-analysis. We leveraged this network for insights into specific areas of biology, including lifespan, frailty, immune composition, histological and functional lung phenotypes, and histological phenotypes of the aorta. We found the genetics of lifespan to share limited correlation with the genetics of frailty but stronger correlation with the genetics of immune cell composition. Additionally, mega-analyses driven by genetic correlations identified candidate genes (e.g. *Cdkn2b*) associated with degraded extracellular matrix in the aorta. Finally, an ensemble of genetic analyses implicated pulmonary neuroendocrine cell signaling and/or differentiation as a key driver of multiple lung pathophenotypes.

## INTRODUCTION

Diversity Outbred (DO) mice are an outbred mouse population consisting of individuals derived from the Collaborative Cross (CC) and are a powerful model system for mapping complex traits (Churchill et al. 2012). The inbred founding strains are genetically diverse and high levels of heterozygosity can lead to sub-megabase (Mb) mapping resolution (Gatti et al. 2014; Churchill et al. 2012; Svenson et al. 2012). Cohorts of genetically unique individuals also provide an intervention testing platform in which strain background effects can be avoided (Nadon et al. 2017).

Versus human cohorts, DO mice provide the typical advantages of in vivo biology, including enhanced opportunities for experimental manipulation. However, salient to the application of genetic analyses, mouse experiments typically use smaller numbers of individuals than human cohort studies, limiting their power. Further, while data are typically shared amongst the DO research community (Vincent et al. 2022), a similar ecosystem of analytical tools that facilitate cross-study analyses is lacking. The DO platform would benefit from implementation of methodologies to enable comparative analysis between independent studies. Ideally, these approaches could be used to inform genetic meta-analysis of phenotypes within and across studies, empowering the detection of quantitative trait loci (QTL) influencing phenotypes of interest.

Genetic correlation (*r_g_*) is one such methodology, which measures the proportion of genetic variance that is shared between two traits due to common underlying genetic effectors (Van Rheenen et al. 2019). This relationship can be useful for discovering shared biological mechanisms or pathways, particularly when the focal traits are genetically complex. Importantly, genetic correlations can be estimated for pairs of traits measured in different cohorts, facilitating cross-study genetic analyses, and can provide insight into gene-by-gene (*GxG*) or gene-by-environment (*GxE*) interactions through the measurement of a single phenotype in different populations, environments, or treatment groups. Multiple methods for *r_g_* estimation are described, but only a residual maximum likelihood (REML)-based method has been implemented for DO mice (Furlotte and Eskin 2015; Zhang et al. 2022). This implementation additionally requires the compared traits to be measured in a common set of animals, limiting it to within-study applications (e.g. Zhang et al. 2022). Another method, linkage disequilibrium score regression (LDSC), can be performed using meta-statistics and is commonly applied to human GWAS; however, this method provides biased heritability and genetic correlation estimates within admixed populations with limited recombination (Bulik-Sullivan et al. 2015; Luo et al. 2021).

Haseman-Elston regression is a method originally developed for the estimation of trait heritability (*h^2^*) and involves regression of phenotypic similarity against kinship across pairs of related individuals (Haseman and Elston 1972). Heritability and genetic correlation are linked concepts (see Methods), implying that Haseman-Elston can be extended to compute *r_g_* for any pair of phenotypes. Its reliance on only between-animal comparisons, independence from a population’s haplotype structure, and straight-forward calculation all evince appropriateness for the estimation of *r_g_* between traits measured in independent cohorts of DO mice.

We implemented Haseman-Elston for DO mice and used it to estimate *h^2^* and *r_g_* across 7,233 phenotypes spanning 11 independent studies. These include traits such as lifespan, frailty, immune cell composition, and newly-reported histological phenotypes in the lung and aorta. Intra-trait *r_g_* estimates between independent longitudinal measurements or intervention groups revealed that the genetic architectures of many complex traits in DO mice are more consistent between age groups than across dietary interventions. Phenotypes were clustered according to *r_g_* patterns across the dataset, then aggregated for QTL discovery via mega-analysis. In total, 884 QTL were detected across 383 meta-phenotypes, with statistically-significant QTL explaining an average of 40.36% of the meta-traits’ heritable phenotypic variances. Patterns of genetic correlations among clusters of traits and lifespan revealed a significant overlap in genetic architecture governing immune cell composition and longevity, while measurements captured by frailty assays were less genetically correlated with lifespan. Finally, *r_g_*-guided meta-analyses of aorta- and lung-derived phenotypes identified candidate genes associated with degradation of the extracellular matrix in the aorta and inflammation and/or tumorigenesis in the lungs.

## RESULTS

### Implementation of Haseman-Elston regression to estimate the heritabilities and genetic correlations of DO mouse traits

We sought to utilize genetic correlations (*r_g_*) to guide mega-analysis of phenotypes from DO mouse populations by identifying groups of traits with shared genetic components. To this end, we implemented an updated form of Haseman-Elston regression in which the cross-product of z-score normalized pairwise trait values (a measure of phenotypic similarity) are regressed on pairwise relatedness to estimate the narrow-sense heritability (*h*^2^) for traits (**Fig S1A-C, Table S1, Data S1-2**). This implementation was tested on simulated phenotypes and produced accurate *h^2^* estimates across a range of *h^2^* values and sample sizes, with precision increasing as a function of the sample size (**Fig. S1D, Methods**). It was further evaluated using real data by comparing *h^2^* estimates it produced to those generated by the REML method. Haseman-Elston and REML produced highly correlated *h*^2^ estimates across 36 traits from (Zhang et al. 2022) that reflected diverse physiology (**Fig. 1A**; ρ = 0.98; p = 1.82*10^-24^).

**Figure 1.**
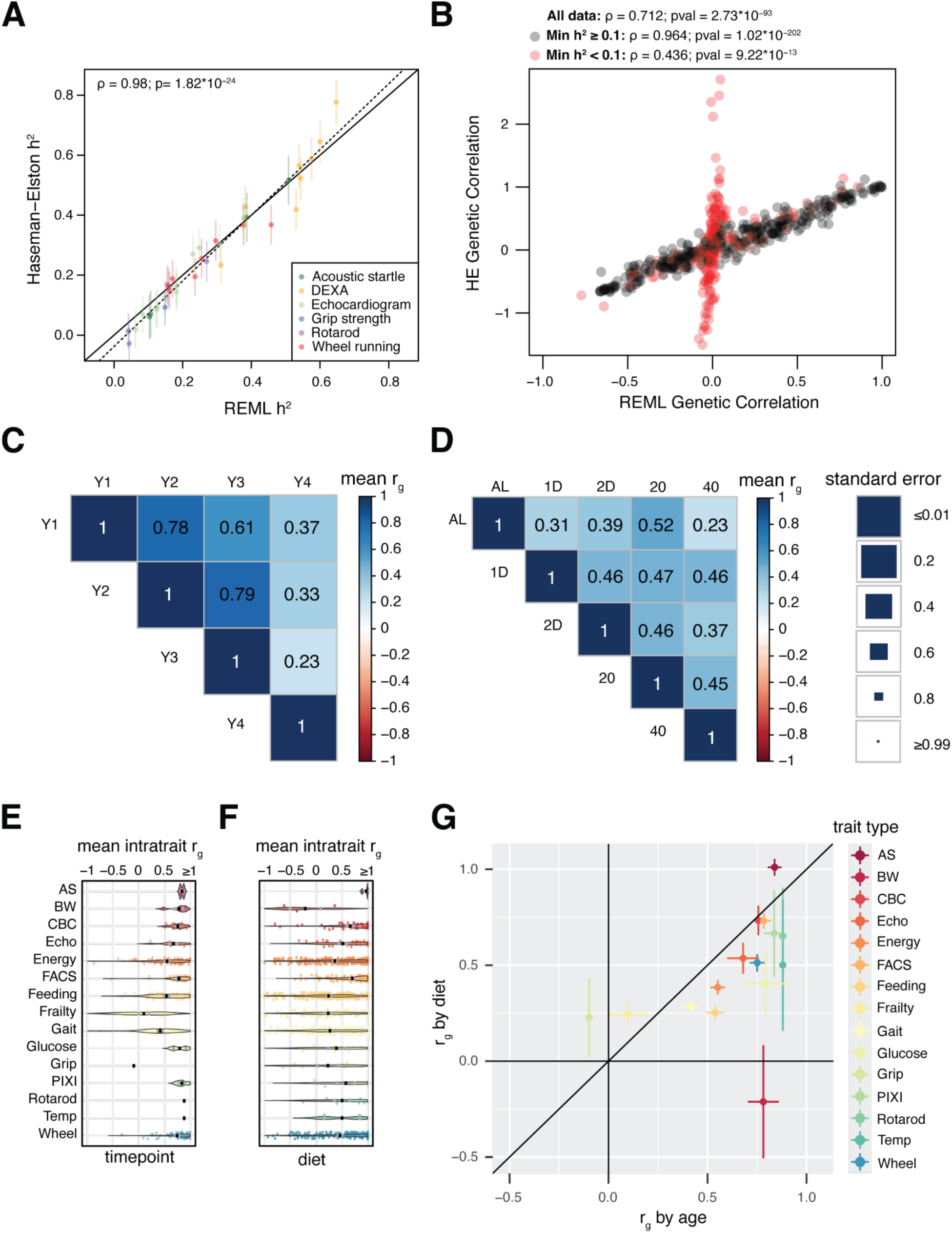
Performance of Haseman-Elston regression implementation and estimation of r_g_ among traits measured across diet and time point. **A**, Comparison of h^2^ estimates for 36 DO mouse traits derived from our implementation of Haseman-Elston regression to those from an orthogonal method, Residual Maximum Likelihood (REML). The line of identity is denoted by a solid line; the best-fit line is denoted by a dashed line. The Pearson correlation coefficient and accompanying p-value are reported in the upper-left of the plot area. **B**, Comparison of r_g_ estimates among all pairs of 36 DO mouse traits derived from Haseman-Elston regression to those from REML. **C**, Mean intratrait *r_g,t_* across time points for individual traits measured in the DRiDO study. Black bars denote the average mean intratrait r_g_ for traits of a particular type, and gray bars represent the mean intratrait r_g_ +/- the standard error of the mean. ‘AS’ - acoustic startle. ‘BW’ - body weight. ‘CBC’ - complete blood count. ‘FACS’ -immune cell composition via fluorescence activated cell sorting. ‘PIXI’ - body composition (lean, fat, bone density). **D**, Mean intratrait *r_g,d_* dietary groups for individual traits measured in the DRiDO study. **E**, Mean intratrait r_g_ across diet and time point for each category of trait, with accompanying standard errors. **F**, Mean genetic correlations among traits measured at different time points in the Dietary Restriction in DO Mice (DRiDO) study. Size of the squares are inversely proportional to the standard error of each mean genetic correlation (*left*). G, Mean genetic correlations among traits measured in different dietary groups in the DRiDO study.

Initial calculation of the standard error (SE) for Haseman-Elston estimates of *h*^2^, using techniques also applied elsewhere (Haseman and Elston 1972), failed to accurately reflect the variances obtained from subset analyses or repeated simulations, becoming less accurate as true *h*^2^ increased. A potential theoretical basis for this discrepancy is that the independence of phenotypic measurements decreases as a function of both the trait *h^2^* and the relatedness of individuals, impacting the number of effective individuals in the analysis; to address this, we derived an empirical ad hoc method for SE calculation, discussed in detail in **Methods**.

We extended Haseman-Elston regression to pairs of traits (i.e. genetic correlation *r_g_*) via a structural equation model **(Fig. S1C**), allowing the estimation of *r_g_* provided there are positive *h^2^* estimates for both traits. Haseman-Elston regression was used to estimate all 630 pairwise genetic correlations among the 36 phenotypes from (Zhang et al. 2022), and those values were compared to *r_g_* estimates produced by REML. Estimates produced by these two methods were positively correlated (**Fig. 1B**; ρ = 0.71; p = 2.73*10^-93^). However, Haseman-Elston produced exaggerated *r_g_* estimates for pairs of traits with low *r_g_* according to REML. Differences between the two methods were inversely proportional to the *h*^2^ of the traits used in the *r_g_* estimates (**Fig. S2A-B**), with REML estimates trending towards zero in low-heritability cases. We hypothesized that the extreme values produced by Haseman-Elston regression were driven by low statistical power, either due to small sample sizes (n) or low *h*^2^ of the traits being compared. Because sample size was relatively large across the 36 traits examined (min=646, max=900, mean=854), we focused on the impact of one or more traits having low *h*^2^. Indeed, the standard error of HE regression estimates increased as either the minimum or mean *h*^2^ of the pair of traits decreased (**Fig. S2C-D**). Haseman-Elston and REML estimates were more strongly correlated when both traits under consideration had *h^2^* ≥ 10% (**Fig. 1B**, black points; (ρ = 0.96; p = 1.02*10^-202^), while estimates were more weakly correlated for trait pairs involving one traits with *h^2^* < 10% (ρ = 0.44; p = 9.22*10^-13^). We therefore imposed a heritability threshold of h^2^≥ 10% for all downstream analysis, unless specified.

### The genetic bases of many complex traits were more sensitive to diet than age

A recent study of Dietary Restriction in Diversity Outbred mice (DRiDO) (Di Francesco et al. 2024) included 429 diverse traits with *h*^2^ ≥10% (**Table S1**). All of these traits were measured in mice on one of five diets: an *ad libitum* diet (‘AL’), a one day intermittent fasting diet (‘1D’), a two day intermittent fasting diet (‘2D’), 20% reduced calorie diet (‘20CR’), or a 40% reduced calorie diet (‘40CR’). Of these traits, 415 had been measured longitudinally (either annually or bi-annually, with at least two measurements across the study) and provided an opportunity to assess the degree to which age or diet modifies the genetic architecture of heritable traits in DO mice.

For each of the 415 traits with annual measurements, *r_g_* was estimated combinatorially between all time points (**Fig. 1C**), and also combinatorially between all diet groups for each yearly measurement (**Fig. 1D**). When averaged across all traits, mean intra-trait *r_g_* estimates (*r̃_g_*) were the highest at younger timepoints, with a marked decrease late in an animal’s lifespan (year 4). With the exception of year 4 data, intra-trait *r̃_g_* estimates were consistently lower among diet groups than time points.

To explore whether these patterns were consistent across diverse sets of traits, the six longitudinal intra-trait *r_g_* values and ten dietary intra-trait *r_g_* values were separately averaged within each individual trait, resulting in mean timepoint intra-trait values (*r̃_g,t_*; **Fig. 1E**) and mean dietary intra-trait values (*r̃_g,d_*; **Fig. 1F**). The average *r̃_g,t_* across the traits was 0.591, revealing substantially shared genetics for traits across ages. The genetics of rotarod (*r̃_g,t_* = 0.880), body temperature (‘Temp’; *r̃_g,t_* = 0.880), and acoustic startle (‘AS’; *r̃_g,t_* =0.839) were the most stable with age; and the genetics of gait (*r̃_g,t_* = 0.417), frailty (*r̃_g,t_* = 0.098), and grip strength (‘Grip’; mean *r̃_g,t_* = −0.098) were the least stable. The average *r̃_g,d_* across traits was 0.390 (**Fig. 1F**), i.e. the genetics of phenotypes measured in that study interacted more with dietary interventions (*GxE*) than with age (*GxAge)*. The genetics of acoustic startle (*r̃_g,d_* = 1.010), hematology (‘CBC’; *r̃_g,d_* = 0.733), and immune cell composition (‘FACS’; *r̃_g,d_* = 0.731) were the most stable across diet and the genetics of frailty (*r̃_g,d_* = 0.244), grip strength (*r̃_g,d_* = 0.228), and body weight (‘BW’; *r̃_g,d_* = −0.212) were the least stable. Across all of the traits measured in the DRiDO study, the only categories in which the genetics were more responsive to age than diet were acoustic startle, grip strength, and frailty (**Fig. 1G**).

### Constructing an atlas of genetic correlations across DO mouse studies

Haseman-Elston regression was next used to broadly characterize the degree to which traits spanning a variety of independent studies and physiological domains share genetic architecture in DO mice. To this end, publicly available data were amassed from 11 studies (sources listed in **Tables S1-2, Data S1**), some of which accompany published manuscripts (Z. Chen et al. 2022; Bagley et al. 2022; Vincent et al. 2022; Gatti et al. 2017; Di Francesco et al. 2024; Mullis et al. 2025; Starcher et al. 2021; Al-Barghouthi et al. 2021; Keller et al. 2018). To enable estimation of pairwise kinship across studies using different genotype arrays for sequencing, genotypes from mice were interpolated across a common set of 69,005 pseudomarkers (**Methods, Data S3**) prior to analysis. Phenotypes from each study were also corrected for age, sex, diet and/or drug treatment (when applicable), and DO generation wave prior to z-score normalization. In total, our analysis drew from 7,233 mice and 15,339 phenotypes, including 12,075 diet-specific phenotypes and 9 aggregated phenotypes consisting of phenotypically clustered frailty measurements (**Methods**). The heritability of each trait was estimated and traits with *h*^2^ < 10% excluded. We also excluded phenotypes for which *h*^2^ exceeded 1 by greater than the standard error associated with the *h*^2^ estimate (**Methods**). After filtering, 7,233 traits remained, including 5,335 diet-specific phenotypes. The mean and median *h*^2^ of the remaining set of high-confidence traits were 45.93% and 34.8%, respectively (**Fig. 2A; Table S3**). The mean and median |*r_g_*| were 0.708 and 0.489, respectively **Fig. 2A**). Of the 26,154,528 trait pairs examined, 276,0833 pairs (10.56%) of traits exhibited individually significant non-zero genetic correlations (p < 0.05), with 13,125 pairs (0.05%) remaining significant after multiple hypothesis correction (p-value ≤1.912*10^-9^) (**Table S4-5**).

**Figure 2.**
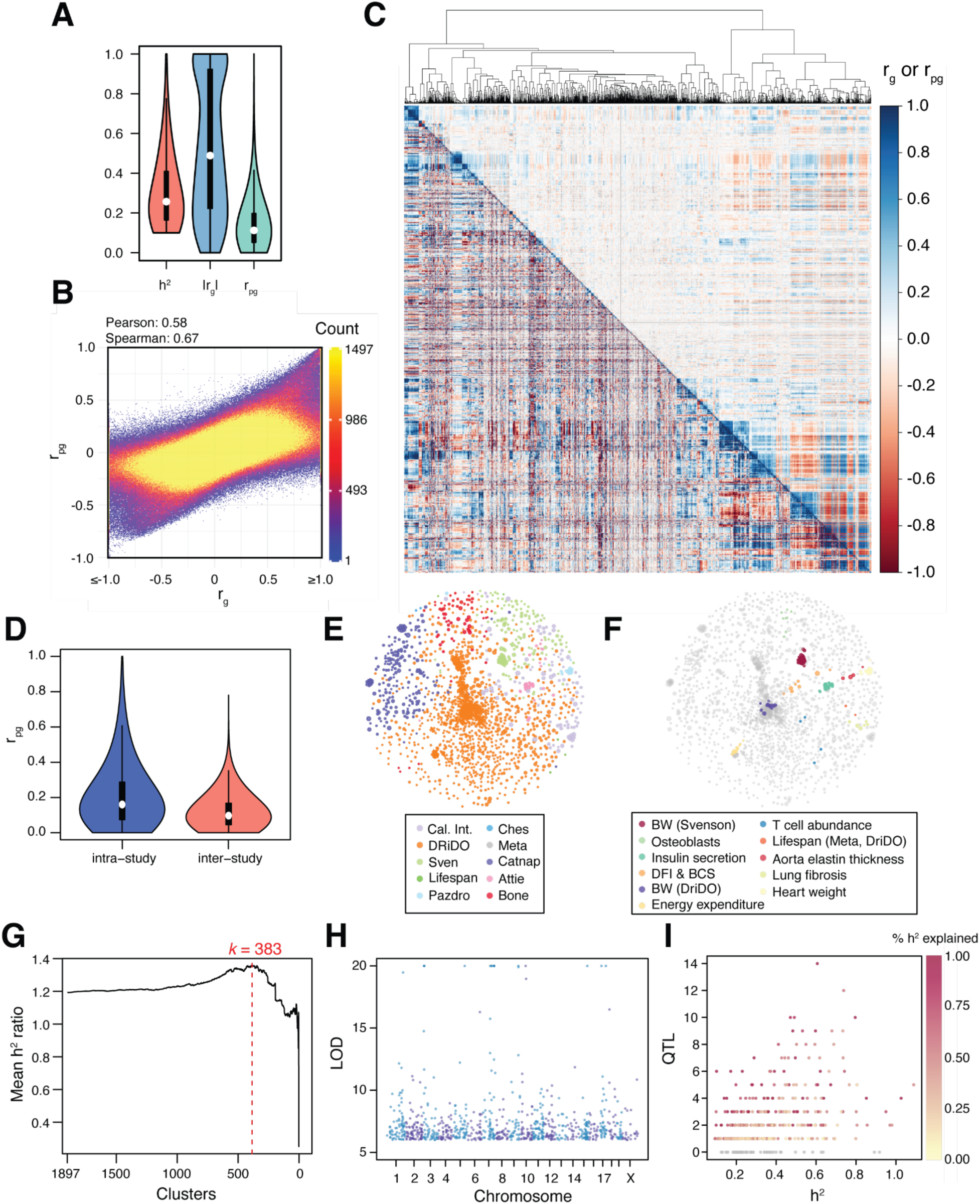
An atlas of r_g_ and r_pg_ estimates among complex phenotypes in DO mice. **A**, The distribution of h^2^, r_g_, and r_pg_, estimates for DO mouse phenotypes used in downstream analysis. The median, 25^th^ percentile, and 75^th^ percentile of the distribution are denoted via box and whisker plots. **B,** The relationship between the pairwise genetic correlation (r_g_) and Pearson correlation of genetic correlations (r_pg_) for each pair of traits in the dataset. Pearson and Spearman correlation coefficients are reported. **C,** A hierarchically clustered heatmap of r_g_ (*lower triangle*) and r_pg_ (*upper triangle*) estimates among 1,898 DO phenotypes. Phenotypes measured in individual dietary groups from the DRiDO study are excluded. The accompanying dendrogram shows euclidean distances among traits based on their r_pg_ estimates. **D**, The distribution of *r_pg_* values among traits measured in the same study (intra-study, blue) or in different studies (inter-study, red). The median, 25^th^ percentile, and 75^th^ percentile of the distribution are denoted via box and whisker plots. **E-F,** Force-directed network diagrams of the 1,898 DO mouse phenotypes used in downstream analysis, with nodes corresponding to individual traits. Networks are colored by study (**E**) and trait type (**F**). *Cal. Int.* - Calico internal phenotypes. *Sven -* Svenson high fat diet study. *Pazdro* - Heart study. *Ches* - Chesler striatum study. *Meta -* meta analysis of lifespan. *Attie* - Pancreas and insulin secretion study. *BW* - bodyweight. *DFI & BCS -* Digital frailty index and body condition score. **G,** Plot of mean h^2^ ratios among clusters of traits at different levels of hierarchical clustering (k). **H**, The location and LOD scores of the 884 QTL detected in meta-analyses of 383 clusters of genetically correlated traits (meta-traits). **I,** The number of QTL detected at LOD ≥ 6 in each of the 383 meta-traits plotted against the h^2^ of each meta-trait. Points are colored by the percent of heritable trait variation explained by the detected QTL.

1,898 of the 7,233 phenotypes were measured in studies that did not include dietary interventions or were corrected for diet prior to analysis. These traits were hierarchically clustered based on their patterns of *r_g_* estimates across the entire dataset of 7,233 traits by taking the Pearson correlation coefficient of *r_g_* vectors (‘Pearson genetic correlation’ or ‘*r_pg_’*, **Table S6**) for each trait pair (**Methods**). *r_pg_* values were significantly correlated with *r_g_* estimates across pairs of traits (ρ = 0.58, p < 2.2*10^-16^; Spearman r = 0.67, p = p < 2.2*10^-16^) but were notably less biased towards extreme values (>1 or <-1) (**Fig. 2A-B**). Traits were then hierarchically clustered by the absolute value of r_pg_, since the strength of genetic correlations, rather than the sign, is indicative of shared genetic architecture (**Fig. 2C**). Phenotypes with higher r_pg_ tended to come from the same study (mean intra-study *r_pg_* = 0.2063, 95% CI = 0.2060 – 0.2065; mean inter-study *r_pg_* = 0.1166048, 95% CI = 0.1165 – 0.1167), an unsurprising result given that the studies integrated into our dataset tended to focus on singular organ systems or domains of physiology (**Fig. 2D-E, Fig. S3**). Traits reflecting similar aspects of physiology or biological processes tended to have higher *r_pg_* with one another, although similar traits measured across studies were not necessarily genetically correlated (“body weight” from the DRiDO study and “body weight” from a high-fat diet study, for example) (**Fig. 2F**).

### Mega-analysis of genetically correlated trait clusters identifies hundreds of QTL

We sought to perform meta-analyses on clusters of genetically correlated traits to identify loci influencing particular aspects of physiology. Maximizing power in such analyses requires defining trait clusters that include data from several genetically correlated phenotypes without under-clustering and aggregating data from genetically unrelated phenotypes. To do this, trait clusters were defined such that the *h^2^* of aggregated meta-phenotypes (*h^2^_meta_*) was maximized relative to unclustered traits using a “*h^2^*ratio” for each trait cluster (**Methods**, **Fig. S4**). We computed the mean heritability ratio of clusters from *k* = 1,897 (minimal clustering) to *k* = 2 (**Fig. 2G**). The heritability ratio consistently increased as traits were clustered until reaching a global maximum at *k* = 383 (4.96 traits per cluster), suggesting that continuous clustering of traits until this point continuously increased h^2^ of the resulting meta-traits relative to previous levels of clustering. After clustering, *r_g_* and *r_pg_* were estimated combinatorially among the 383 phenome-wide meta-traits (C_1_ - C_383_; **Fig. S5**) to facilitate downstream analysis (**Tables S7-S12**).

Genome-wide QTL scans were performed on meta-traits C_1_ - C_383_ (**Fig. 2H, Data S4**). At permutation-based, trait-specific, genome-wide significance thresholds, 164 total QTL were discovered (**Table S13**); while 884 QTL were identified at a conservative, FDR-based global threshold of LOD > 6 (**Table S14**; see **Methods**). Loci discovered using permutation-based thresholds explained an average of 3.5% of phenotypic variation in their respective meta-analyses, while inclusion of loci discovered at LOD > 6 threshold increased this percentage to 19.46% (**Fig. 2I**). The mean *h*^2^ of the 383 meta-traits was 36.5%, and the mean percentage of *h*^2^ explained by loci detected at genome-wide significance thresholds was 8.13%. Inclusion of loci at LOD > 6 increased this percentage to 40.36%.

### The genetics of lifespan are modestly correlated across independent DO studies

The atlas of *r_g_* data included lifespan measurements from three independently conducted studies of lifespan in DO mice (“Harrison”, “Shock”, and “DRiDO”), as well as a meta-trait combining those three studies (“meta_lifespan3”) (Mullis et al. 2025; Di Francesco et al. 2024). Collectively, these studies include lifespan measurements for 2,308 mice (232 males and 2076 females) across six diets (ad libitum, or “AL”; 20%, 30%, and 40% caloric restriction, or ‘20CR’, ‘30CR’, and ‘40CR’; one and two day intermittent fasting, or ‘1D’ and ‘2D’) and one drug treatment (rapamycin, or ‘rapa’) (**Table S4**). Haseman-Elston-based estimates of *h^2^* were 11.1% (SE = 8.3%), 12.6**%** (SE = 3.6%), and 23.7% (SE = 6.1%) for the Shock, Harrison, and DRiDO studies, respectively. These were similar to previously-reported REML-based *h^2^* estimates of 15.0% (SE = 15%), 18.5% (SE = 6.5%), and 24.6% (SE = 7.7%) from the three studies, respectively (Mullis et al. 2025).

The mean of individual *h^2^* estimates for lifespan from the DRiDO, Harrison, and Shock studies was 15.8% (SE = 4.0%), while the *h^2^* of meta-lifespan, combining data from those three studies, was 11.7% (SE = 2.3%) (**Fig. 3A**), suggesting some genetic factors contributing to lifespan to have had study-specific effects. Concordantly, the inter-study *r_g_* estimates for lifespan were not statistically distinct from zero, and inter-study lifespan *r_pg_* values were only significantly non-zero between the DRiDO and Shock studies (*r_pg_* = 0.11; p = 1.34*10^-22^). The Harrison study had significant *r_pg_* estimates with the other lifespan traits when correcting for the limited number of lifespan-to-lifespan correlations hypothesized (three hypotheses; Harrison-DRiDO: *r_pg_* = 0.06, p = 9.69 * 10^-7^; Harrison-Shock: *r_pg_* = −0.03, p = 4.5*10^-3^). As might be expected given their relatively low *r_pg_* values, study-specific lifespan traits were not clustered into the same phenome-wide meta-traits. The *r_pg_* estimates computed among the meta-traits were used to determine whether lifespan traits were genetically correlated with the same meta-traits (**Fig. S5 – 6**). The vector of *r_pg_* values from the DRiDO study was significantly correlated with the *r_pg_* values from both the Shock and Harrison studies, while the *r_pg_* values from the Shock and Harrison studies were not significantly correlated (**Fig. 3B**). The most highly correlated meta-traits were largely distinct between the three lifespan studies (**Fig. S6**).

**Figure 3.**
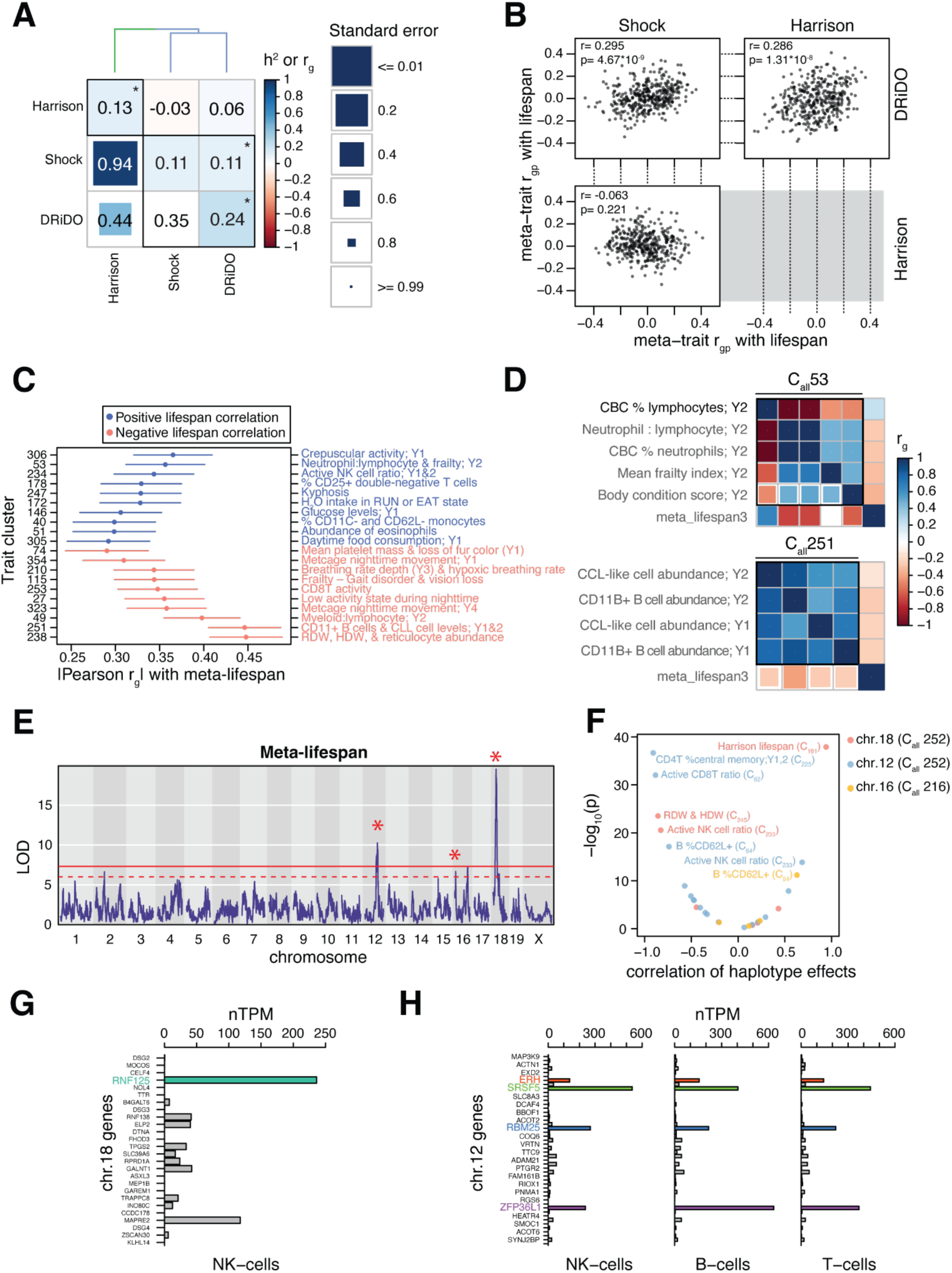
Genetic correlations involving measurements of lifespan in DO mice. **A**, Heatmap of genetic correlations and heritability estimates among three independent measurements of lifespan in DO mice. The diagonal of the heatmap reports the h^2^ of each lifespan trait. The lower triangle of the heatmap shows r_g_ among the lifespan traits and the upper triangle shows the r_pg_ among lifespan traits. Asterisks denote statistically significant h^2^, r_g_, or r_pg_ estimates. The dendrogram shows the results of hierarchical clustering of the lifespan traits based on euclidean distance derived from the r_pg_ values among each pair of traits. Clusters of traits that result in a maximal mean h^2^ ratio when combined are represented by colors in the dendrogram as well as black boxes in the heatmap. **B,** Scatterplots of r_pg_ estimates between lifespan traits and the 380 other non-lifespan meta-traits. **C,** The strongest positive and negative r_pg_ estimates between meta-traits and combined lifespan data. Error bars correspond to the standard error of each r_pg_ estimate. Meta-trait number (cluster number) is shown on the left hand side of the plot and a brief summary of the phenotypes included in each meta-trait is shown on the right. **D,** Examples of trait clusters that, when aggregated into meta-traits, have high r_pg_ estimates with combined lifespan data. C_all_ 53 (*top*) comprises midlife (year 2) traits reflecting relative abundances of neutrophils and lymphocytes in the blood and includes frailty index and body condition scores. C_all_ 251 (*bottom*) comprises CD11+ B and chronic lymphoid leukemia-like (CLL) cell abundance traits from early and midlife (years 1 and 2). **E,** Manhattan plot of an additive whole-genome scan of a mega-analysis of lifespan consisting of data from the DRiDO, Harrison, and Shock lifespan studies. The red line indicates a genome-wide significance threshold based on 1,000 permutations of the data. The dashed red line indicates a FDR-based significance threshold of LOD >= 6 (corresponding to an FDR of ∼9%). QTL mapped at genome-wide significance in the mega-analysis or one of the individual studies comprising the analysis are denoted by asterisks. **F,** Volcano plot depicting the correlation coefficient and -log_10_(p) of haplotype effects at lifespan loci with overlapping QTL from meta-analyses of other trait clusters. **G,** Expression of genes within the 2LOD support interval of the lifespan QTL on chr. 18 in human Natural Killer (NK) cells. **H,** Expression of genes within the 2LOD support interval of the lifespan QTL on chr. 12 in human NK, B, and T cells.

Together, these results were consistent with a limited – but nonzero – overlap in the genetic basis of independently-collected lifespan measurements. Inclusion of varied interventional arms across these studies, including dietary modifications, may have contributed to their varied genetics. Genetic correlation estimates *r_pg_* for lifespan assessed across various dietary intervention and pharmacological treatment groups were consistently positive (mean |*r_pg_*| = 0.09, SE = 0.012), with 17 of 28 pairwise comparisons (60.1%) demonstrating statistical significance, indicating a modest shared genetic architecture influencing lifespan under different intervention conditions (**Fig. S7**).

### Common genetic drivers influence lifespan and immune cell traits

Despite genetic differences in lifespan among studies, a previous meta-analysis of lifespan in DO mice detected six longevity-associated QTL, including a locus on chromosome 18 that reproducibly detected in two independent studies and additional loci mapped at genome-wide significance in individual studies on chromosomes 12 and 16 (**Fig. 3C**, **Fig. S8**, Mullis et al 2025). Motivated by these findings, we considered genetic correlations among the meta-traits and aggregated lifespan data. The meta-traits sharing the highest |*r_pg_*| with meta-lifespan included frailty measurements (Ruby et al. 2023), various metabolic cage readouts (Z. Chen et al. 2022), and many immune cell phenotypes (**Fig. 3D,E**) (Di Francesco et al. 2024). Mega-analysis of C_1_-C_383_ identified 19, 6, and 9 QTL with 2LOD support intervals that overlapped with the chromosome 12, 16.1 and 18 loci, respectively.

To determine if the lifespan QTL and those detected in other meta-traits were driven by the same DO founder alleles, we first performed variant association mapping at each lifespan QTL to determine a set of markers most likely to explain the effect of each locus, defined as the 1% of markers with the most significant effect on lifespan within the 2LOD support interval of each QTL (**Data S5**). For each marker set, we calculated the Pearson correlation coefficient of the best linear unbiased predictors (BLUPs) of each DO founder allele on lifespan and on any overlapping QTL from C_1_ - C_383_. For all loci, the most significant correlations between allelic effects on lifespan and other traits (p ≤ 10^-10^) occurred between lifespan and meta-traits related to ratios of activated to naive B, T, and NK cells (**Fig. 3F**). Notably, both lifespan-associated loci were detected in cluster C_all_233 (ratio of active to inactive NK cells in late life), and the same alleles appeared to drive effects on both sets of phenotypes. On chromosome 18, the CAST allele is associated with both decreased lifespan and an increased ratio of active NK cells; on chromosome 12, the CAST allele is associated with increased lifespan and decreased ratios of active immune cells, including CD4+ T cells, CD8+ T cells, CD62L+ B cells, and NK cells (**Fig. 3F**).

The 2LOD support intervals of the lifespan QTL on chromosome 12 and 18 encompass 48 and 26 genes, respectively (**Data S5**). We utilized single-cell RNA-seq data from the Human Protein Atlas (Uhlén et al. 2015; Karlsson et al. 2021) to identify candidate genes within these intervals that are expressed in the relevant cell types for each lifespan QTL. At chromosome 18, ubiquitin ligase *Rnf125* is the most highly expressed gene in human natural killer cells (NK-cells) and has been shown to be a marker of NK-cells in both humans and mice (**Fig. 3G**) (Bezman et al. 2012; Crinier et al. 2018). At the chromosome 12 locus, a set of four genes appear more highly expressed across NK-cells, B-cells, and T-cells: *Erh*, *Srsf5*, *Rbm25*, and *Zfp36l1* (**Fig. 3H**). The chromosome 16 locus was resolved to a single protein-coding gene, the RNA binding protein *Rbfox1*, which is expressed at low levels in B-cells.

### Mid-life frailty indexes shared limited genetic correlation with lifespan

We sought to quantify the extent to which frailty indexes reflect the underlying genetic basis of lifespan by genotyping animals from a prior study on frailty performed using DO mice (Ruby et al. 2023). Heritability and genetic correlations were calculated for both a video-based digital frailty index (DFI) and a manual frailty index (MFI) (Kane et al. 2017; Whitehead et al. 2014), as well as for the individual components of each frailty index. We also implemented a version of the traditional open field assay (Belzung 1999) amenable to the varied coat colors of DO mice (see Methods), and collected a large corpus of DO data using this assay in order to include the phenotypes measured in our genetic analysis. Due to precedence for inclusion of open field in frailty assessment (Whitehead et al. 2014), these traits were analyzed in conjunction with MFI and DFI. Across animals from which both open field and FI data were collected (N = 125; see **Methods**), each FI score and chronological age had negative phenotypic correlations with the performance-based open field phenotypes (total distance travelled, gait speed, angular rotation, distance per rotation) and positive correlations with exploration of the middle of the box (**Fig. S9A**).

Haseman-Elston-based *h^2^* estimation yielded 0.854 (SE = 0.238) for MFI and 0.600 (SE = 0.237) for DFI. In agreement with the phenotypic correlations for these phenotypes measured in DO mice by (Ruby et al, 2023; R = 0.22; p value = 1.8 × 10^−7^), these two versions of FI shared significantly positive genetic correlations (r_g_ = 0.484, p value = 0.041; r_pg_ = 0.492, p value < 9.88*10^-324^). The most highly-powered lifespan phenotype (meta-lifespan) was not genetically correlated with MFI (r_g_ = −0.001, p value = 0.998; r_pg_ = −0.023, p value = 0.048); but shared a significant negative Pearson genetic correlation with DFI (*r_g_* = 0.173, p value = 0.582; r_pg_ = −0.184, p value = 5.20*10^-56^) (**Fig. 4A**).

**Figure 4.**
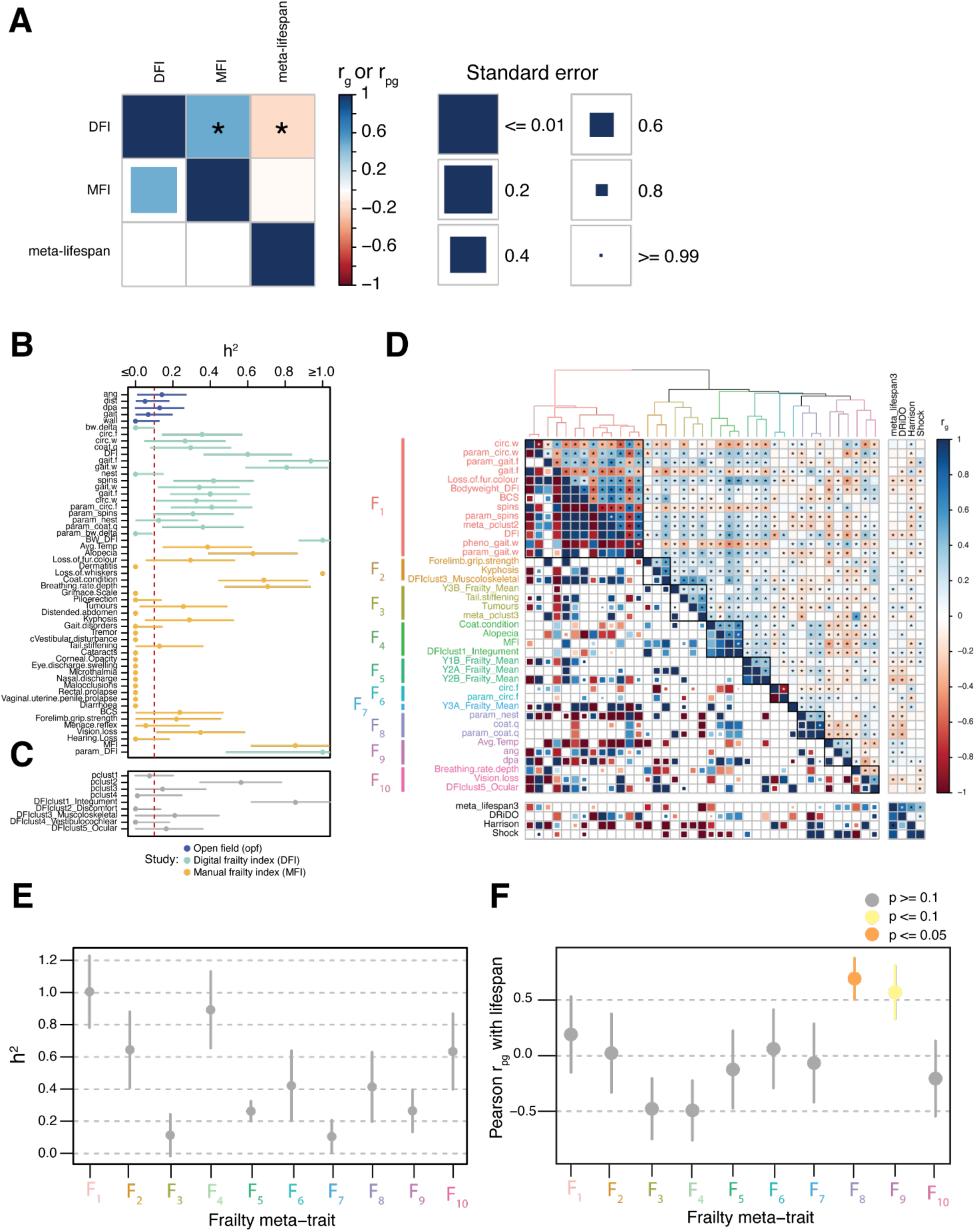
Genetic correlations among frailty and lifespan phenotypes. **A**, Pairwise (*r_g_*, bottom triangle) and Pearson (*r_pg_,* upper triangle), genetic correlations among Manual Frailty Index (MFI), Digital Frailty Index (DFI), and lifespan. Statistically significant genetic correlations are denoted with an asterisk. Box color represents the magnitude of the genetic correlation and box size is inversely correlated with the size of the standard error. **B-C,** Heritability and standard error of individual (**B**) and aggregate (**C**) frailty phenotypes. The dashed red line at h^2^ = 0.1 is the threshold used to determine if phenotypes were included in downstream analysis. **D,** Hierarchically clustered heatmap showing r_g_ (*lower triangle*) and r_pg_ (*upper triangle*) among frailty phenotypes. The accompanying dendrogram shows euclidean distances among traits based on their r_pg_ estimates. Clusters of frailty traits (F_1_ - F_10_) that result in a maximal mean h^2^ ratio when combined are represented by colors in the dendrogram as well as black boxes in the heatmap. r_g_ and r_pg_ among frailty and unclustered lifespan phenotypes are shown in the adjacent panels of the heatmap. **E,** Heritability and standard error of the meta-traits comprising clusters of genetically correlated frailty phenotypes. **F,** r_pg_ and standard error among frailty meta-traits and a meta-analysis of lifespan spanning three studies.

While the insignificance of most *r_g_* and *r_pg_* values between the frailty indexes and lifespan suggested largely non-overlapping genetics, the possibility remained that individual FI components might possess greater but obfuscated correlations, warranting individual analyses. Across the 55 scored components of MFI, DFI, and open field, 37 (67%) had significantly non-zero *h^2^*, with 32 (58.2%) exceeding the 10% *h^2^* threshold we imposed for genetic correlation analysis (**Fig. 4BA, Table S3**). One more trait (“loss of whiskers”) was excluded from subsequent analysis because its estimated *h*^2^ exceeded 100% by more than its standard error. The remaining 31 frailty components (mean, median h2 = 32.3%, 25.6%) had been included in the already-described phenome-wide genetic correlation analysis. We additionally constructed nine aggregate phenotypes (**Fig. 4C**) based on either *a)* a phenotypic correlation analysis (**Fig. S9B**), or *b)* groupings defined by the authors of the MFI (Whitehead et al. 2014). Of the resulting nine aggregate traits, five had *h^2^* ≥ 10%.

Genetic correlations were then estimated among the 31 individual frailty phenotypes, the five heritable aggregate traits, and the rest of the mouse phenome (including lifespan). Individually, none of the individual or aggregate frailty traits had significant *r_g_* with meta-lifespan; however, 18 of the 36 frailty traits (50.0%) had significant *r_pg_* with meta-lifespan when accounting for phenome-wide multiple testing (p value ≤ 1.91*10^-9^) (**Fig. 4D**). Among these, the mean |*r_g_*| with meta-lifespan was 0.364 and the mean |*r_pg_*| with meta-lifespan was 0.092, implying a low, but non-zero, overlap in the genetics of frailty traits and lifespan. Individual and aggregate frailty phenotypes (including MFI and DFI) were hierarchically clustered into 10 meta-traits (F_1_ - F_10_) based on their phenome-wide *r*_pg_ values such that the *h^2^* of the resulting meta-traits was maximized (**Fig. S9C**, **Fig. 4E, Table S15**). We estimated each frailty meta-trait’s genetic correlation with meta-lifespan and used these estimates to compute *r*_pg_ between frailty meta-traits and lifespan (using only the correlation across lifespan and these 10 meta-traits). Of the 10 frailty meta-traits, only two appeared to share the most genetic overlap with lifespan: F_8_ (nesting and coat quality; *r_pg_* = 0.69, p = 0.018) and F_9_ (temperature and two open field behavioral traits; *r_pg_* = 0.57, p = 0.067) (**Fig. 4F**).

### Meta-analysis of histology-derived aorta phenotypes reveals genetic drivers of elastin breakage

A major advantage of DO mice is the ability to perform genetic analyses on phenotypes that would be difficult or impossible to collect from humans, e.g. histological analysis of non-expendable tissues. We conducted a novel histological study of the aortic extracellular matrix using 116 DO mice. Image analyses were performed on aortic cross-sections stained with H&E, TC, or VVG, yielding 26 phenotypes, 21 of which had *h^2^* ≥ 10% (see **Methods**; **Fig. S10**). Phenotypes were hierarchically clustered into seven meta-traits (A_1_ - A_7_, **Table S16**) using their phenome-wide *r_pg_* values, such that the *h^2^* ratio was maximized across meta-traits (**Fig. 5A; Fig. S11**). Broadly, these meta-traits comprised *1)* elastin laminae thickness & % aortic area consisting of elastin, *2)* the number of elastin layers in the aorta, *3)* elastin breakage and tortuosity, *4)* tissue area, *5)* nuclei per tissue area, *6)* external collagen, and *7)* nuclear area and fraction of collagen inside the aorta. A meta analysis of each cluster was performed (**Data S6**), identifying three QTL at permutation-based significance and an additional 14 loci at a FDR-based significance threshold of LOD >= 6 (Mullis et al. 2025), for a total of 17 QTL (2.43 QTL/meta-trait) (**Fig 5B-D; Fig. S12**).

**Figure 5.**
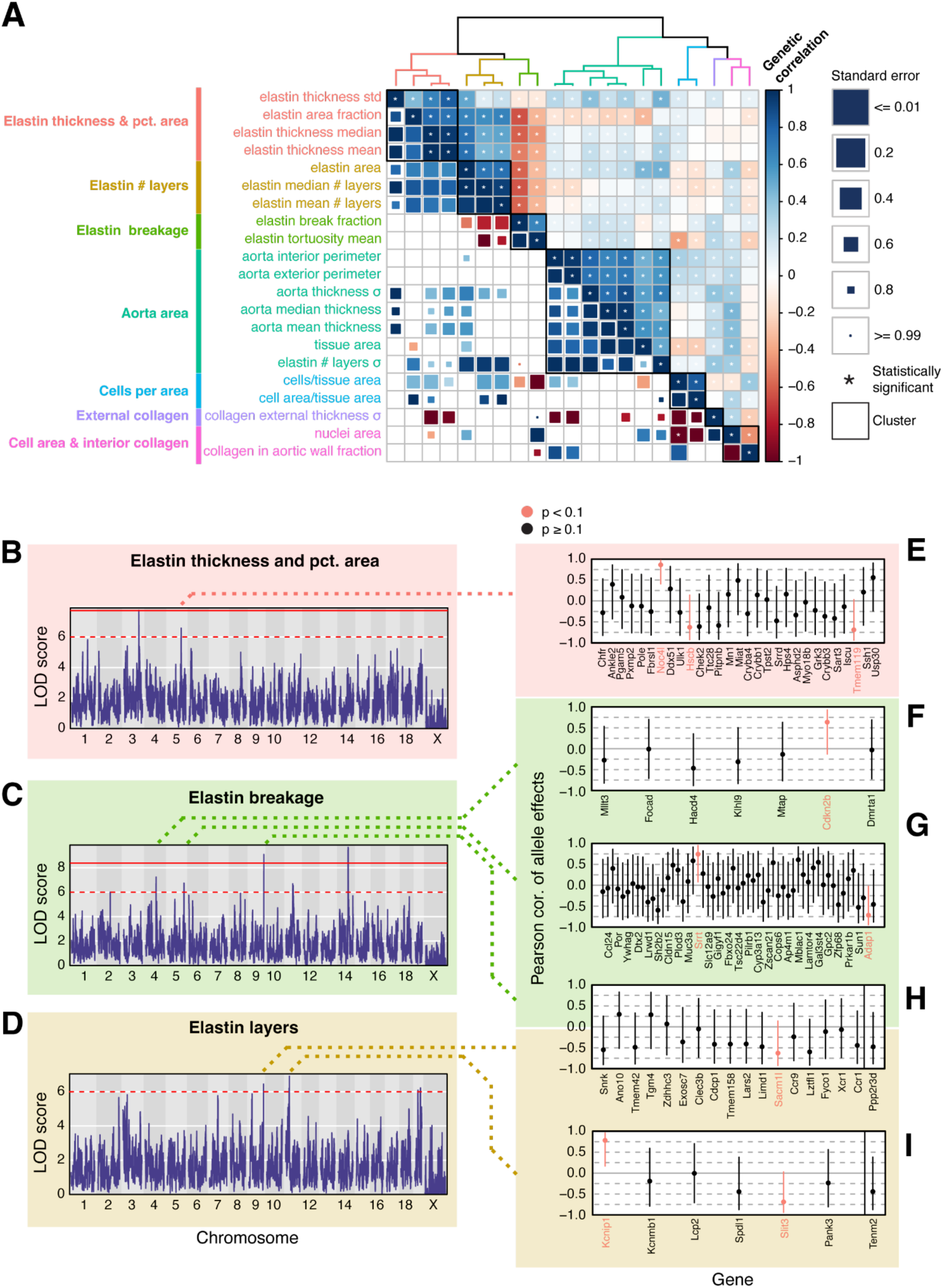
Genetic correlations among extracellular matrix traits in the aorta. **A**, Hierarchically clustered heatmap showing r_g_ (*lower triangle*) and r_pg_ (*upper triangle*) among aorta phenotypes. The accompanying dendrogram shows euclidean distances among traits based on their r_pg_ estimates. Clusters of aorta phenotypes that result in a maximal mean h^2^ ratio when combined are represented by colors in the dendrogram as well as black boxes in the heatmap. **B,** Manhattan plot of an additive whole-genome scan on aggregated aortic elastin thickness and pct. area phenotypes. The red line indicates a genome-wide significance threshold based on 1,000 permutations of the data. The dashed red line indicates a nominal significance threshold of LOD >= 6. **C,** Manhattan plot of an additive whole-genome scan on aggregated aortic elastin breakage phenotypes. **D,** Manhattan plot of an additive whole-genome scan on aggregated aortic elastin layers phenotypes. **E–I,** Correlations of haplotype effects between the peak markers at QTL influencing elastin phenotypes and *cis*-eQTL in an external DO cardiac tissue transcriptomic dataset. Candidate genes are highlighted with red text.

Two, six, and four QTL were detected associated with A_1_ (elastin thickness and percentage of aortic area consisting of elastin area), A_2_ (layers of elastin in the aorta), and A_3_ (elastin breakage), with a locus on chromosome 9 contributing to both A_2_ and A_3_ (**Fig. 5, Table S17**). We estimated effect sizes (best linear unbiased predictors, or BLUPs) and performed association mapping on the 2LOD support interval about each QTL (**Fig. S13-14**). Collectively, the QTL explain 28.59%, 51.87%, and 106.91% of variation in elastin area, breakage, and layers, respectively; however, due to the low sample sizes associated with these phenotypes (mean n = 112.57), the variance attributable to the QTL are likely inflated due to the Beavis effect. Most confidence intervals were large, spanning an average of 33 protein-coding genes (**Data S7**), although two QTL were mapped at single gene resolution: *Olfm4* influencing elastin breakage on chromosome 14 and *Sorcs1* influencing the number of layers of elastin on chromosome 19 (see **Discussion**).

To narrow down the list of candidate genes influencing elastin in the aorta, we relied on data from an external whole-heart RNA-seq experiment (Gerdes Gyuricza et al. 2022). For each QTL, we searched for expression QTL (eQTL) associated with genes within 2Mb of the peak marker in our meta analysis. We then computed the Pearson correlation coefficient of allele effects between the peak markers in the RNA-seq and meta-analysis datasets (**Fig. 5E-I; Fig S15**). For five of the QTL, one or more candidate genes were identified with expression patterns driven by the same alleles contributing to elastin phenotypes in the aorta. Notably, we identified *Cdkn2a* as the most likely candidate within the elastin breakage QTL on chromosome 4 (**Fig. 5F**; see **Discussion**).

### Meta-analysis of functional and histology-derived lung phenotypes reveals genetic contribution to pulmonary inflammation and/or tumorigenesis

In order to broadly characterize the genetic landscape of lung function in DO mice, we measured 79 functional lung phenotypes, including image-derived phenotypes from uCT scans as well as H&E and TC stains of lung tissue sections, in another distinct cohort of 240 mice, including 124 C57BL/6J and 116 DO mice (**Fig S16, Fig. 6**). The patterns of phenotypic correlations were highly similar when measured within the C57BL/6J versus DO populations (Pearson correlation across phenotype correlations = 0.737; p value < 2.2*10^-16^; **Fig S17A-B**), suggesting physiological generalizability between these inbred and outbred mice.

**Figure 6.**
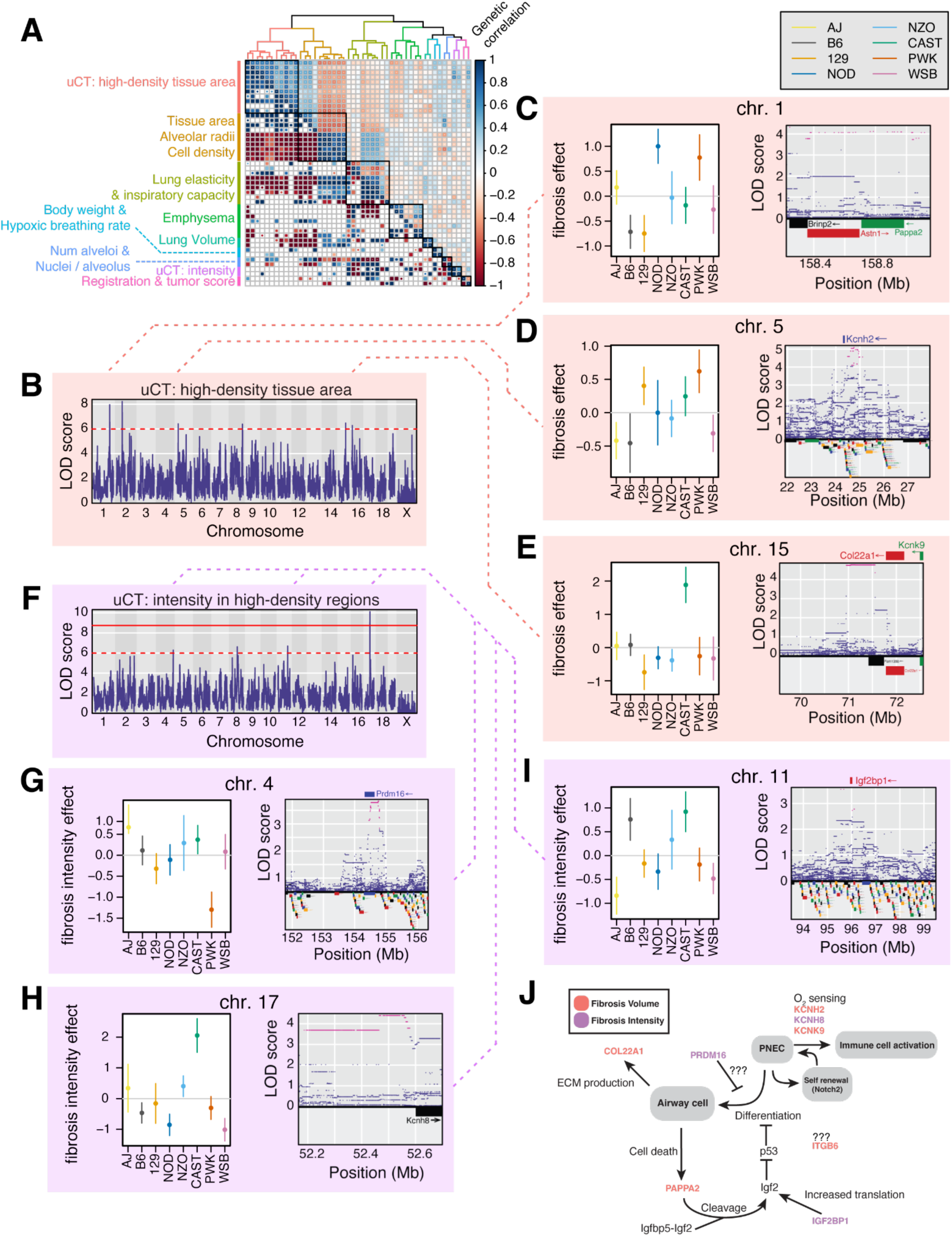
Genetic correlations among lung structural and functional phenotypes. **A**, Hierarchically clustered heatmap showing r_g_ (*lower triangle*) and r_pg_ (*upper triangle*) among lung phenotypes. The accompanying dendrogram shows euclidean distances among traits based on their *r_pg_* estimates. Clusters of lung phenotypes that result in a maximal mean *h^2^* ratio when combined are represented by colors in the dendrogram as well as black boxes in the heatmap. **B,** Manhattan plot of an additive whole-genome scan on aggregated fibrosis volume phenotypes. The red line indicates a genome-wide significance threshold based on 1,000 permutations of the data. The dashed red line indicates a nominal significance threshold of LOD >= 6. **C–E,** Allelic effects (BLUPs) and corresponding standard errors of the QTL influencing fibrosis volume in the lung (*left*), and variant association mapping of the 2 LOD support intervals around each QTL (*right*). Genes within the confidence intervals are shown in the bottom panel, and candidate genes are shown above variant association results. **F,** Manhattan plot of an additive whole-genome scan on aggregated fibrosis intensity phenotypes. **G–I,** Allelic effects (BLUPs) and corresponding standard errors of the QTL influencing fibrosis intensity in the lung (*left*), and variant association mapping of the 2 LOD support intervals around each QTL (*right*). **J,** Model depicting the cellular functions of pulmonary neuroendocrine cells (PNECs) and showcasing the roles of candidate genes may play in contributing to fibrotic uCT readouts in the lung. Candidate genes are colored by which whole-genome scan they were detected in.

Of the 79 lung phenotypes, *h*^2^ estimates met or exceeded 10% for 49 (62%; **Fig S17C**). Those 49 phenotypes were therefore included in downstream genetic analyses. As above, phenotypes were hierarchically clustered by their *r_pg_* values across the 1,898 phenotypes and grouped to maximize the h^2^ ratio of each meta-trait (**Fig. S18**), resulting in 9 clusters of traits (L_1_ - L_9_; **Fig. 6A**) enumerated in **Table S18**. Meta analyses were performed on each cluster, identifying a single QTL at permutation-based thresholds and an additional 31 loci at LOD >= 6, for a total of 32 QTL (3.56 QTL/meta analysis, **Fig. 6B, 6F; Fig. S19, Table S19**).

Two clusters of lung phenotypes – L1 and L8, corresponding to the % of high-density tissue and voxel intensity of high-density tissue, respectively – were uCT-derived lung tissue density measurements that may reflect several pathological conditions of the lung, including pulmonary fibrosis, inflammation, tumor presence, or edema (Vande Velde et al. 2016). To better understand the nature of the uCT-derived tissue density phenotypes, we examined the phenotypic relationships between uCT measurements and fibrosis measurements derived from trichrome (TC) staining of lung tissue sections, which are expressed in the percent area of the lung that stains for collagen (**Fig. S20A-G**). In C57BL/6J mice, the TC-derived collagen fraction increases from day 120 to day 600 (**Fig. S20A**), while the % area of high-density tissue and voxel intensity in high-density areas decreases between 120 and 150 days before increasing with age (**Fig. S20B-C**). Due to the resolution of the measurements, it is unclear whether the same trend occurs for collagen ratio in the lungs. In DO mice, the fraction of collagen and voxel intensity of high-density areas increased with age but not the fraction of high-density tissue (**Fig. S20D-F**). The area of high-density tissue was not significantly correlated with collagen fraction (r = −0.131, p = 0.299; **Fig. S20G**) but was significantly correlated with mean distance between nuclei (r = −0.424, p = 3.85*10^-6^; **Fig. S20H**) and presence of ectopic cell clusters (r = 0.414, p = 7.05*10^-6^; **Fig. S20I**), both of which are hypothesized to reflect inflammation and/or tumorigenesis in the lung tissue.

Meta analysis of uCT-derived tissue density phenotypes resulted in the detection of 6 QTL influencing the % high-density tissue and 4 QTL influencing the voxel intensity of high-density tissue (**Fig. 6B, 6F, Table S18, Data S8**). These QTL explain 94.2% and 62.7% of phenotypic variation in the traits, respectively; however, due to low sample sizes these estimates of QTL effects are likely inflated due to the Beavis effect (Xu 2003). Confidence intervals at the % high-density tissue and voxel intensity QTL spanned 19 and 49 genes on average, respectively (**Data S9**). However, variant association mapping resolved the locus on chromosome to the *cis*-regulatory region of a single gene, *Kcnh8*, which encodes a voltage-gated potassium channel expressed specifically in pulmonary neuroendocrine cells (PNECs) within the lung (**Fig. 6I**) (Kuo et al. 2022). Two additional potassium channels with PNEC-specific lung expression patterns, *Kcnh2* and *Kcnk9*, were identified within the 2LOD support intervals of QTL on chromosomes 5 and 15, respectively (**Fig. 6D-E**), with variant association mapping refining the set of candidate SNPs at chromosome 5 to those in the 5’ upstream region of *Kcnh2* (**Fig. 6D**). *Pappa2*, a protease that activates PNEC mitogen *IGF2* via cleavage of the *IGFBP5-IGF2* complex (Barrios et al. 2021), was one of three genes underlying the % high-density tissue QTL on chromosome 1 (**Fig. 6C**). Variant association mapping localized the effect at the chromosome 4 locus to candidate SNPs in the upstream region of *Pdrm16* (**Fig. 6G**) (Fei et al. 2019) and *Igf2bp1*, which increases levels of bioavailable *IGF2* through stabilizing interactions with its mRNA (Dai et al. 2011), on chromosome 11 (**Fig. 6H**).

## DISCUSSION

Our novel implementation of Haseman-Elston regression for the estimation of *r_g_* in DO mice provides a tool by which phenotypes can be jointly analyzed across studies, timepoints, environments, and intervention groups. We first used *r_g_* estimates to assess the extent to which age and dietary interventions impact the genetic architecture of complex traits including energy expenditure, food and water consumption, hematology, adiposity, frailty, immune cell, and body weight. Intra-trait *r_g_* suggested that gene-by-diet (*GxE*) interactions played a greater role in specifying phenotype than gene-by-age (*GxAge)* interactions across nearly all types of traits, with three exceptions – acoustic startle, frailty and grip – all of which relate to the deterioration of the body with age. Notably, the mean intra-trait *r_g_* among body weight phenotypes was negative implying a shared genetic architecture in which some variants have opposite effects on body weight depending on an animal’s diet. While intra-trait *r_g_* was higher among traits at early timepoints, these estimates decreased on average when comparing measurements from earlier timepoints (years 1 - 3) to those taken late in life (year 4). This finding suggests that the effects of variants may change more rapidly later in life, or it may reflect a survival bias present in the DRiDO study: most of the animals that survived to the fourth time point were under severe (40%) caloric restriction, which was previously shown to have dramatic effects on longevity and other health-related outcomes.

### Overcoming the limited power of DO mouse genetic studies through meta-analysis

Extending Haseman-Elston based *r_g_* estimates to over 7,000 DO mouse phenotypes facilitated a cross-study hierarchical clustering of traits based on genetic similarity. While individual *r_g_* estimates were noisy, hierarchical clustering based on patterns of genetic correlations (*r_pg_*) across the entire dataset produced 383 clusters of traits that, when aggregated for meta-analysis, increased statistical power for QTL discovery. To date, no larger meta-analysis has been conducted in DO mice, and our approach resulted in the detection of hundreds QTL influencing lifespan, immune composition, aortic tissue composition, and lung function, and the QTL detected in meta-analyses explained a large fraction of variance (over 40%) in their respective meta-traits. While more detailed genetic analyses were performed on several phenotypes of interest in this study (*see below*), many sets of correlated phenotypes remain to be analyzed, and we hope this dataset will be valuable to those interested in genetic analysis in DO mice.

The granularity of clustering used in this analysis was specified to maximize mean *h^2^* across all meta-analyses. However, we note that power increases in meta-analyses can arise from either i) combining multiple measurements of correlated traits from the same set of animals, or ii) combining correlated traits measured in different sets of animals. The former can increase the signal-to-noise ratio (*h^2^*) of genetic analysis if the error terms associated with combined traits from the same animals are orthogonal and cancel out when the traits are aggregated. The latter increases the sample size through the inclusion of additional animals, which will not increase *h^2^* but should increase the precision of the estimate. In practice, meta-analyses performed in this paper allow for both approaches towards trait clustering but as clustering was optimized for maximum *h^2^*, trait clusters in this analysis may be biased towards the combination of measurements from within cohorts. Future implementations may incorporate the standard error of the *h^2^*estimate as a criteria for clustering. Furthermore, a global solution to maximize *h^2^* may not produce clustering ideal for any one particular set of phenotypes; alternative clustering solutions for individual phenotypes may be explored in the future.

### Identification of common genetic drivers between mouse lifespan and correlated phenotypes

Estimates of *r_pg_* between lifespan measurements from three independent studies (Harrison, Shock, and DRiDO) revealed significant but modest correlations, suggesting potential differences in the genetic architecture of lifespan across study. Each lifespan study was conducted independently in different facilities/rooms using different chow, feeding schedules, temperatures, housing arrangements, and personnel, all of which could potentially redistribute mortality risk and influence the effects of genetic variants on lifespan. The inclusion of different experimental intervention groups may have also influenced the genetic factors responsible for longevity. For instance, the Harrison study examines dietary and pharmacological lifespan interventions in female mice, while the Shock study focuses on lifespan in males in females without intervention (Mullis et al. 2025). Lifespan phenotypes from these studies were not genetically correlated (**Fig. 3A**) and were correlated with different sets of meta-traits (**Fig. 3B**). Meanwhile, lifespan data from the DRiDO study, which includes large numbers of animals with and without interventions (Di Francesco et al. 2024), had patterns of *r_pg_* estimates with meta-traits that were significantly correlated with both the Harrison and Shock studies (**Fig. 3B**), implying that the genetics of lifespan in the DRiDO study likely encapsulates genetic effects relevant to both of the other two studies.

Despite low *r_pg_* among independent lifespan measurements, some QTL influencing lifespan are detected in multiple studies (Mullis et al. 2025) and are reproduced in a meta-analysis of longevity in DOs. A lifespan locus on chromosome 18, for example, was previously reported to affect variation in red blood cell volume (Di Francesco et al. 2024); here, the locus also shows correlated effects on the ratio of CD11b^-^/CD11c^-^ to CD11b^+^/CD11c^+^ natural killer (NK) cells that may be driven by the ubiquitin ligase *RNF125*. A second lifespan locus on chromosome 12 has correlated effects on several clusters of genetically correlated immune cell abundance and activity traits and implicates *ERH*, *SRSF5*, *RBM25*, or *ZFP36L1* as genes potentially influencing lifespan in DO mice. Collectively, our results suggest that variation in genes related to the immune system are contributing the most to variation in lifespan across DO mouse studies.

Frailty indexes (FIs), which each measure multiple clinically defined signs of deterioration in mice (Whitehead et al. 2014; Ruby et al. 2023), generally produce scores that correlate with age and are used as a measurement of overall health in animals. However, the extent to which these indexes reflect the underlying genetics of lifespan are unknown. While many individual frailty measurements were not significantly genetically correlated with lifespan, aggregate index scores (MFI, DFI, and clusters of phenotypically correlated frailty metrics) did often have significant, albeit weak, *r_pg_* with lifespan. One interpretation is that these individual measurements may capture some fraction of the genetic variation contributing to lifespan in addition to other components unrelated to longevity, while the signal from the variants associated with aging or longevity are retained in the aggregate scores. Clustering analysis of frailty traits revealed two clusters, F8 and F9, that were most genetically correlated with lifespan. These clusters represented traits such as nesting behavior, coat quality, temperature, and movement-related traits from the open field behavioral assay, indicating that a subset of frailty traits could potentially serve as surrogate markers for predicting longevity, albeit with limitations.

### Genetic contributors to the structural health of the mouse aorta

Diseases of the aorta are a significant contributor to human mortality, driven in part by the loss of the elastic properties of the tissue due to fragmentation, thinning, and improper cross-linking of elastin fibres. Mutations in the elastin gene *Eln* and other aortic extracellular matrix (ECM) proteins (e.g. fibrillins and collagens) are known to negatively affect the structure and function of the aorta, leading to adverse cardiac events. However, outside of proteins that comprise the aortic ECM, little is known about the genetic factors that influence the microstructure of aortic elastin filaments. Meta-analysis of elastin phenotypes revealed 12 novel QTL influencing elastin structure in the aorta; notably, none of these loci overlapped *Eln*, *Fbln-2*, *Fbln-5*, *Mmp-2*, or *Mmp-9*, genes implicated in aortic dissection and other cardiac events (Cheuk and Cheng 2011; Koullias et al. 2004; Nakashima 2010; Tsamis, Krawiec, and Vorp 2013). However, several notable candidate genes were identified, including *Cdkn2a*, a cyclin-dependent kinase that had previously been mapped in a genome-wide association study of human lifespan (Wright et al. 2019) and was previously associated with cardiovascular disease (Van Der Harst and Verweij 2018). Another notable candidate was *Olfm4*, which plays an important role in innate immunity and inflammation. Knockdown of this gene reduced inflammatory response in lipopolysaccharide (LPS)-stimulated rat cardiac cells, resulting in enhanced proliferation and decreased apoptosis. *Olfm4*, an olfactomedin domain-containing glycoprotein regulated by NF-κB, is another noteworthy candidate that is involved in innate immunity and inflammation (Liu and Rodgers 2016) and plays a protective role in the digestive tract (Wang et al. 2018). Chen *et al* found that suppression of *Olfm4* has a protective effect in lipopolysaccharide (LPS)-stimulated cardiomyocytes (H. Chen, Liu, and Fang 2024), and the gene has emerged as a biomarker for the severity of certain infections (Liu and Rodgers 2022). Association of this gene with elastin breakage suggests a genetic link between inflammation and degradation of the ECM within the aorta.

### Genetic contributors to the functional health of the mouse lung

MicroCT-based imaging of the lung in DO mice revealed numerous candidate genes influencing variation in tissue density of the lungs of adult mice. These measurements are unlikely to reflect pulmonary fibrosis, but may capture inflammation, a key characteristic of interstitial lung disease (ILD). Interestingly, three of the candidate genes (*Kcnh2, Kcnh8,* and *Kcnk9*) are voltage-gated potassium channels expressed robustly in pulmonary neuroendocrine cells (PNECs) and have been proposed as a mechanism by which PNECs function as O_2_ sensors within the airways (Kuo et al. 2022). In addition to this role, PNECs have been shown to regulate inflammation in the lung through the secretion of CGRP, suggesting a potential relationship between O_2_ sensing, immune cell recruitment, and fibrosis in the lung. PNECs may also function as stem cells in response to airway damage (Reynolds et al. 2000; Ouadah et al. 2019), which may be induced via IGF2-IGFR signaling,and are putative precursors to small cell lung cancer (SCLC)(Song et al. 2012). Notably, two additional candidates, *Pappa2* and *Igf2bp1*, may influence PNEC stem cell activity by modulating levels of bioavailable IGF2; the former through cleavage of IGFBP5, which binds IGF2 and prevents signaling, and the later through stabilizing mRNA encoded by the *Igf2* locus. A summary of candidate genes, and how they relate to the functions of PNECs, is depicted in **Fig. 6J**. In addition to candidates directly implicated in PNEC biology, *Prdm16* has been shown to play a role in suppressing both pulmonary fibrosis via antiproliferative effects (Fei et al. 2019; Tan et al. 2021). Together, the candidates identified in our analysis suggest an important role of PNEC biology in ILD and/or tumorigenesis.

### Discovery opportunities deriving from future data-sharing across the DO mouse research community

DO mice are a powerful model system for studying the genetics of complex traits, with great potential to parallel and complement human genetic studies. The human and DO mouse genetic research communities have common commitments to the open sharing of data. In the case of DO mice, the lack of personal privacy concerns enables easy sharing of raw individual-level data. Although large, open-access databases of DO phenotypes and mapping results have already been established (Vincent et al. 2022), the DO community currently lacks the breadth of tools and resources available for facilitating cross-study genetic analysis compared to human cohorts. The implementation of Haseman-Elston regression to estimate genetic correlations among independently-collected traits provides a new mechanism for collaborative analysis across independently-conducted DO studies that is akin to the utility provided to human studies by the LDSC tool (Bulik-Sullivan et al. 2015). Here, we have used it to unite the corpus of DO physiological and behavior studies into a common framework, enabling insights that would not have been possible from those studies in isolation.

We have sought to be comprehensive in our integrated analysis of available DO mouse data. However, the true value of these analyses was realized through focus and review of the patterns of clustering and QTL from particular domains of biology. Here, we focused on gleaning insight from our personal areas of interest: lifespan, frailty, and elastic tissues/organs (aorta and lung).

We hope that the availability of these clusters will enable similar insights across domains where we have chosen not to focus our attention – including immune cell composition, hematological parameters, cardiovascular function, body weight, adiposity, feeding, circadian rhythm, pancreatic function, and bone strength. As new DO mouse studies are published, we hope to continue expanding this network into new domains using the Haseman-Elston method. In particular, recent expansions in available transcriptomic, metabolomic, proteomic and lipidomic measurements from DO mice (e.g. (Keller et al. 2018; Starcher et al. 2021; Skelly et al. 2020)) set the stage for the construction of comprehensive molecular correlation matrices.

## METHODS

### Dataset curation

Data used in these analyses were collected from a variety of sources, including internal Calico studies and publicly available studies of DO mice (**Table S2**). In order to standardize genetic analysis, only datasets including age, sex, and DO generation wave information were included in our analysis.

### Genotyping

Genotypes for Calico internal studies were obtained using the mouse universal genotyping array (GigaMUGA; 143,259 markers) (Morgan et al. 2016). Genotyping was performed from tail tips by Neogen Genomics (Lincoln, NE, USA). Founder genotypes were reconstructed using the R/qtl2 software (Broman et al. 2019)and samples with call rates at or above 90% were retained for analysis.

Mice from all studies were genotyped using the MUGA, MegaMUGA, or GigaMUGA genotyping arrays (**Table S2**) and genotypes were called using R/qtl2. To facilitate analysis across different arrays, genotype probabilities from each study were interpolated to a set of 69,005 pseudo-markers (**Data S3**) and converted to founder haplotype probabilities via hidden Markov model in R/qtl2.

### Adjustment and normalization of trait values

To facilitate analysis of phenotype data across studies, all phenotypes were processed using a common workflow. First, for each phenotype, outliers –defined as trait values differing from the mean by an excess of five standard deviations– were excluded. To account for common covariates in DO mouse studies, we fit the following linear fixed-effects model:

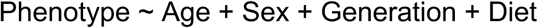

Where ‘Age’ corresponds to the age of the animals in days at the time of trait measurement, ‘Sex’ corresponds to the sex of the animals, ‘Generation’ corresponds to the DO mouse generation wave, and ‘Diet’ corresponds to any dietary interventions or drug treatments used in the studies (when applicable). We performed z-score normalization on the residuals of each model and the resulting phenotype scores were used for downstream analysis.

Phenotypes from the DRiDO study included mice from different dietary groups. These phenotypes were included in the analysis as both aggregate traits spanning diet groups (which included a categorical ‘Diet’ term in the linear model) and as individual, diet-specific phenotypes. Diet-specific phenotypes were fit using fixed-effects models without a diet term and were processed and z-normalized separately from aggregated phenotypes.

Phenotypes from the Svenson HFD study included mice that were fed a high fat diet for different numbers of days. These phenotypes were modeled using a continuous ‘Diet’ term that accounted for the number of days each mouse spent on the diet.

### Estimation of narrow-sense heritability of traits via Haseman-Elston (HE) regression

We estimated the narrow-sense heritability (h^2^) of traits in Diversity Outbred (DO) mice using Haseman-Elston (HE) regression, a statistical method in which the z-normalized cross product of trait values among pairs of related individuals (**Fig S1A**) are regressed on their corresponding kinship estimates, as in **Fig S1B**. The model used in this regression takes the form of:

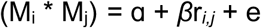

Where *i* and *j* are a pair of related individuals, (M_i_ * M_j_) is the product of the z-score normalized trait values of trait *M* for *i* and *j*, and r*_i,j_* is the standardized kinship estimate of *i* and *j* by identity by state (IBS). The slope, β_Y_, of the resulting best-fit line represents the covariance in trait values of Y as a function of relatedness, and can be converted to a phenotypic correlation of Y with itself as a function of additive genetic effects (in other words, h^2^) via the following equation:

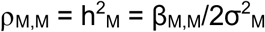

Where h^2^_M_ is the narrow-sense heritability of trait Y and σ^2^_M_ is the variance in the measurement of Y (Visscher et al. 2006). Because *M* is normalized, σ^2^_M_ is equal to 1 and the equation is simplified to:

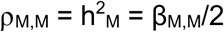

The relationship between genotype (G), h^2^, phenotype (P), and correlation of trait values due to additive genetic variance (⍴_M,M_) for two individuals is depicted in a structural equation model in **Fig 1C**.

The standard error about the HE regression coefficient was computed using the following formula:

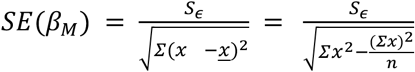

Where *SE*(*β*_*M*_) is the standard error of the HE regression coefficient for trait M in the downsampled population, x refers to pairwise kinship estimates on which the cross product of M is regressed, and S_Ɛ_ is the standard error of the estimate, defined as:

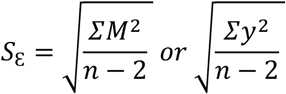

Where *M* or *y* are the cross-product of trait values for trait *M* that are regressed on pairwise kinship estimates and *n* is the number of pairwise observations. Then, as with the regression coefficient, the standard error of the HE regression coefficient was then divided by twice the variance of the z-score normalized trait *M* used in the regression:

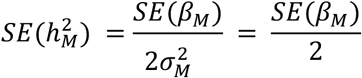

### Assessing the performance of HE regression

To assess the performance of our implementation of HE regression, we performed simulations in which HE regression was used to estimate the narrow-sense heritability of phenotypes with known genetic architecture and h^2^ values using the ‘PhenotypeSimulator’ package in R [**ref here**]. Genotypes for simulated individuals were constructed from 1,000 unlinked diploid markers at seeded minor allele frequencies of 0.05, 0.1, 0.3, or 0.4. Kinship was computed from standardized genotype values and divided by 2 to reflect the probability of sampling the same allele twice from a highly heterozygous individual, as in DO mice. Simulated fixed genetic effects were additive and entirely independent of one another; in each simulated population, 10/1000 genetic markers were randomly selected as causal and effect sizes of these markers were drawn from a normal distribution. Individual phenotypes were computed without noise from the simulated genotypes and independent background noise was added to the set of phenotypes according to the h^2^ of the trait being simulated. At each sample size and h^2^, 1000 different populations and sets of phenotypes were simulated, and HE regression estimates were computed (**Fig. S1D**).

We also compared the performance of our implementation of HE regression to an orthogonal Bayesian model that was previously used to estimate h^2^ for 36 physiological traits in DO mice (Zhang et al. 2022). Heritability estimates were computed on the residual values for these 36 traits after correcting for age, sex, generation wave, and diet using a linear fixed-effects model of the form:

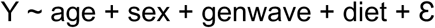

where sex, generation wave, and diet were encoded as factors and age in days was encoded as a numeric variable. The resulting HE regression estimates were compared directly to the heritability estimates reported in the original study (**Fig. 1A**).

### Extension of Haseman-Elston regression to genetic correlations

HE regression can also be applied to the estimation of genetic correlations for pairs of traits. This involves regressing the z-score normalized cross product of trait values for two traits, *M* and *N*, among pairs of individuals on their pairwise kinship estimates (**Fig. S1A-B**):

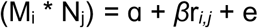

The slope of the resulting best-fit line, β_M,N_, represents the covariance in the cross-product of traits *M* and *N* as a function of relatedness, and can be converted to a phenotypic covariance of traits *M* and *N* as a function of additive genetic effects using a slightly modified form of the prior equation:

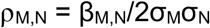

Which again simplifies due to the use of z-score normalized trait values in the regression:

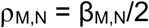

While ⍴_M,M_ is equivalent to narrow-sense heritability for trait *M*, because trait *M* is perfectly genetically correlated with itself by definition, this relationship does not hold true for a pair of traits that are not genetically equivalent (**Fig S1C**). Instead, ⍴_M,N_ is a function of the heritabilities of *M* and *N* as well as the genetic correlation of the two traits:

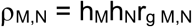

Where h_M_ and h_N_ are the square root of traits *M* and *N*, respectively, and r_g M,N_ is the genetic correlation of *M* and *N*. Thus, the genetic correlation for a pair of traits can be computed from three pieces of information: h^2^ estimates for each individual trait and HE regression coefficient for the pair of traits.

The standard error about HE regression coefficients for traits *M* and *N*, as well as for the corresponding *h^2^* estimates were computed as detailed above. Standard error estimates were carried through the structural equation model to compute SE for the r_g_ estimate via the following equation:

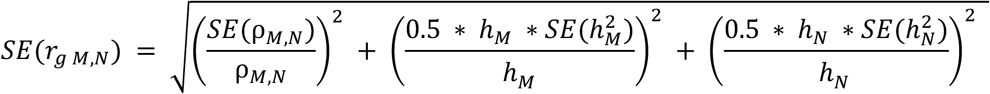

***Empirical derivation of a standard error (SE) correction factor for HE regression*** Derivation of a standard error correction factor for HE regression was performed using simulated phenotypes and populations. Simulations for each phenotype at each population size were repeated 1,000 times.

Phenotypes at varying *h^2^* levels (0.01, 0.05, 0.1, 0.2, 0.4, 0.8, 0.99) were simulated for populations of 10,000 individuals via the ‘PhenotypeSimulator’ package in R. Genotypes for each population were simulated at 1,000 SNPs. 10 SNPs were selected to be causal with independent, additive effects on their respective phenotype, with minor allele frequencies sampled from the following list of values: 0.05, 0.1, 0.3, 0.4) and were standardised as described in Yang et al (Yang, J., Lee, S.H., Goddard, M.E., Visschzer, P.M. (2011). Kinship matrices were computed via the getKinship() function and values were divided by 2 to match kinship matrices computed via the ‘QTL2’ package in R, which computes the probability of sampling a particular allele among pairs of individuals.

For each simulated phenotype, the corresponding population was split randomly into two subpopulations consisting of 5,000 individuals. These subpopulations were then downsampled to varying population sizes ranging from 220 to 5,000 individuals. For each downsampled population, *h^2^* was computed via Haseman-Elston regression and the standard error of the *h^2^* estimate was computed as detailed above.

To assess whether these standard error estimates were appropriate, we utilized the approach taken by Ruby *et al*. in which *h^2^*estimates and standard errors from each set of 1,000 simulated populations were used to generate expected and observed differences among the corresponding downsampled subpopulations (Ruby et al. 2018, see supplemental text section 5.1-2, supplemental figure S2). These expected and observed differences were then used to generate a vector of ratios using the formula:

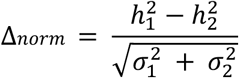

Where *Δnorm* is the vector of expected:observed differences, *h^2^_1_* and *h^2^_2_* are the heritability estimates of the two downsampled subpopulations, and *σ^2^_1_* and *σ^2^_2_* are the variances about the heritability estimates of the two subpopulations (Ruby et al. 2018) (see Ruby et al. 2018, supplemental text equation 56). Repeated across a sufficiently large number of samplings, the standard deviation of *Δnorm* values, σ_Δ_*_norm_*, should be approximately 1, with values <1 or >1 signifying a greater or lesser precision of the SE estimate than the null expectation. σ_Δ_*_norm_* was used to assess the precision of HE standard errors across population size and *h^2^*.

After finding that standard error estimates decreased in precision (σ_Δ_*_norm_* > 1) as a function of both the *h^2^* of the phenotype and the size of the population (**Fig. S21**), we utilized a linear regression based approach to find a correction factor, *F*, that would result in the expected precision of the standard error terms (σ_Δ_*_norm_* ∼ 1) across phenotypes of varying *h^2^* measured in populations of varying size. Across a series of heritability values (0.1, 0.2, 0.4, 0.8, 0.99), σ_Δ_*_norm_* increased linearly with the square root of the sample size (**Fig. S22**), and so the following linear model was used to fit σ_Δ_*_norm_* as a function of √n at each *h^2^*:

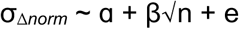

Where ɑ, β, and e are the intercept, slope, and error of the regression, respectively. Secondary linear regressions were then run that fit the resulting ɑ and β terms as a function of *h^2^* (**Fig. S23**):

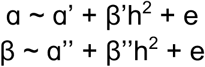

The square root of the sample size, ɑ’, ɑ’’, β’, and β’’ were then used to create a SE correction factor, *F*, that should result in SE estimates with the expected levels of precision at a particular *h^2^* and sample size:

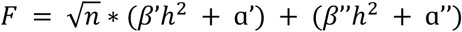

The empirically derived formula for F used in the estimation of SE of *h^2^* and *r_g_* estimates in this manuscript is:

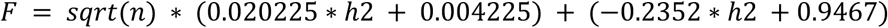

And corrected SE estimates (*SE_corrected_*) are derived via:

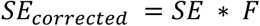

This correction factor was then applied to SE estimates for 4,529 experimental traits in DO mice that varied in both *h^2^* and sample size. Each trait was split randomly into two samples and *Δnorm* was computed. σ_Δ_*_norm_* for the set of traits was 1.160 (compared to 1.203 using uncorrected SE values), suggesting that SE estimates for traits in our dataset have an approximately expected level of precision in our data (**Fig S24**). Considering only individual traits with relatively large sample sizes (n ≥ 800), σ_Δ_*_norm_* was 1.199 (compared to 1.320 using uncorrected SE values), suggesting that our correction factor reduces the loss of precision in SE estimates as sample size increases.

### Intra-trait genetic correlations

Intra-trait genetic correlations were estimated for phenotypes measured in **D**ietary **R**estriction **i**n **DO** mice (DRiDO) study (Di Francesco et al. 2024). These phenotypes were measured annually (or bi-annually) for up to four years per animal as well as across five dietary groups (ad libitum, or “AL”; 20% and 40% caloric restriction, or ‘20CR’ and ‘40CR’; one and two day intermittent fasting, or ‘1D’ and ‘2D’).

To estimate the average intra-trait r_g_ among yearly time points or diet groups, r_g_ was estimated for each trait among each pair of time points or diets. For each trait with bi-annual measurements (denoted as ‘YearX_A’ and ‘YearX_B’), the first measurement was used. r_g_ estimates greater than 1 or −1 by more than the standard error of the r_g_ estimate (r_g_ > 1+se or r_g_ < −1 - se) were excluded from the analysis. After intra-trait r_g_ was estimated for all traits across all pairs of time points or diets, the mean and standard error of intra-trait r_g_ for all traits was reported for each pair of time points or diets via heatmap in **Fig. 1C-D**. Individual yearly traits were used in the estimation of intra-trait r_g_ across diets.

To assess patterns of intra-trait r_g_ across trait categories, all pairwise r_g_ estimates were calculated across all pairs of time points or diet groups. r_g_ estimates greater than 1 or −1 by more than the standard error of the r_g_ estimate (r_g_ > 1+se or r_g_ < −1 - se) were excluded from the analysis. For each trait, the mean and standard error of its r_g_ estimates among time points or diets was calculated and mean values are reported via violin plots in **Fig. 1E-F** and summarized in **Fig. 1G**.

### Hierarchical clustering of traits based on Pearson genetic correlations (r_pg_)

For each pair of traits, Pearson genetic correlations (r_pg_) were estimated using z-score normalized vectors of genetic correlations (r_g_) from the entire dataset of 7,233 high-confidence phenotypes, which included phenotypes from the DRiDO and lifespan studies that were derived from individual dietary groups or drug treatments. High confidence phenotypes were defined as those with *h^2^* estimates ≥ 0.1. Because Haseman-Elston regression can also produce h^2^ estimates greater than 1 due to error associated with the slope of the regression (ꞵ), we also excluded any traits with *h^2^* estimates that exceeded 1 by greater than their standard error estimate (*h^2^* > 1+se(*r_g_*) were excluded).

Because positive and negative genetic correlations are both indicative of a shared genetic basis, the absolute value of *r_pg_*, or |*r_pg_*|, was used for clustering analysis. To facilitate hierarchical clustering of traits, |*r_pg_*| values were converted to euclidean distance using the formula:

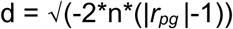

Where d is the euclidean distance between the pair of traits, n is the number of observations – in this case, r_g_ measurements– used to estimate *r_pg_*, and |*r_pg_*| is the absolute value of the Pearson genetic correlation estimate for the pair of traits.

Hierarchical clustering was performed on a matrix of squared distances among 1,898 high-confidence phenotypes (diet- and drug-specific phenotypes were excluded) using the “ward.D2” hierarchical clustering algorithm from the function hcut() from the R package “factoextra” (Kassambara and Mundt 2020). The level of clustering, *k*, that maximized average *h*^2^ when performing meta-analysis of trait clusters, was used (see “*Construction of meta-traits from clustered phenotypes to maximize h^2^” below)*.

### Determination of a level of hierarchical clustering to maximize h^2^

Because *r_g_* and *r_pg_* indicate the degree to which pairs of traits share underlying genetic effects, traits with higher *r_g_* or *r_pg_* values should be more likely to arise from one or more of the same causal variants. Combining data from genetically correlated traits should increase the statistical power to detect shared QTL, which may increase the *h^2^*estimate of the aggregated phenotype data in a mega-analysis. To test this, we combined 1,000 random pairs of traits (see ***Construction of meta-traits*** below) and estimated the *h^2^* of the resulting meta-traits (*h^2^_meta_*). For each pair, we took a ‘heritability ratio’:

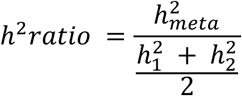

Where *h^2^_meta_* is the heritability of a meta-trait resulting from the combination of two sets of trait data, *h^2^_1_* is the heritability of the first combined trait, and *h^2^_2_* is the heritability of the second. There was a highly significant correlation between the degree of shared genetic architecture (*|r_pg_|*) and the *h^2^* ratio for the pairs of traits (r = 0.328, p = 1.38* 10^-26^; **Fig. S4**), implying that a ‘heritability ratio’ could be used to determine an appropriate level of clustering in our data.

Heritability ratios were applied to the phenome-wide dataset (1,898 traits) to determine a level of clustering, *k*, that would maximize the average *h^2^* when performing mega-analyses. At each possible level of hierarchical clustering (i.e. from 2 clusters of traits to 1,897 clusters of traits), *h^2^* ratios were computed as follows:

At each level of clustering, *k*, we first cut the dendrogram to specify *k* clusters of traits. Within each cluster, a meta-trait was constructed by computing the mean of the z-score normalized phenotype values for each animal in the cluster. Next, *h^2^_meta_* was estimated for each of the *k* meta-traits using HE regression. Each meta-trait comprises two sub-dendrograms, which represent the two sets of traits grouped together at a particular level of clustering; for each meta-trat, the *h^2^* of its constituent sub-dendrograms, *h^2^_sub1_* and *h^2^_sub2_* were also estimated. The *h^2^* ratio for each meta-trait was defined as *h^2^_meta_* divided by the mean of *h^2^_sub1_* and *h^2^_sub2_*, expressed as:

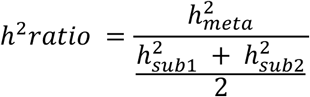

The mean *h^2^* ratio was computed across all *k* meta-traits at a particular level of clustering. We selected a level of clustering that maximized the mean of the *h^2^* ratio among all groups of traits, which should maximize our statistical power to detect genetic effects influencing meta-traits.

### Construction of meta-traits

Meta-traits were constructed from sets of clustered phenotypes at a particular level of clustering (see above) by taking the mean of the z-score normalized phenotype values within each trait cluster. Because traits were hierarchically clustered based on the absolute value of their Pearson genetic correlation (|r_pg_|), traits within particular clusters may be negatively genetically correlated with one another. To facilitate the construction of meta-traits from negatively correlated phenotypes, a “focal phenotype” was assigned from each cluster, defined as the trait with the highest h^2^ estimate. Each other trait within a cluster was then compared to this focal phenotype; traits with negative r_pg_ estimates were sign-flipped and traits with positive r_pg_ estimates were not. After this processing, the average of the clustered phenotypes was taken as the meta-trait value for each animal. Animals lacking any phenotype measurements for a given trait cluster were assigned “NA”.

### Additive whole-genome scans of meta-traits

Genetic analysis was conducted using the “rqtl2” package in R (R Core Team 2024; Broman et al. 2019). Whole-genome scans were performed for each meta-trait using the scan1() function utilizing a mixed effects model in which a focal trait is regressed on 8-state allele probabilities for each individual. Because age, sex, DO generation wave, and intervention (diet or drug treatment, referred to as ‘Diet’ above) were already accounted for during the preprocessing of phenotype data for Haseman-Elston regression and subsequent meta-trait construction, no additive covariates were used in whole-genome scans.

For each scan, a significance threshold was established using 1,000 permutations of the data, with each permutation consisting of a whole-genome scan in which meta-trait values were randomized. The maximum LOD score observed in each permutation was recorded, resulting in a distribution of 1,000 maximum LOD scores. The 95^th^ percentile of this distribution was used as the significance threshold (ɑ = 0.05) for the corresponding meta-trait. QTL with LOD scores greater than this threshold were considered significant at our permutation-based threshold. In addition to this significance level, a significance level of LOD ≥ 6 was also used to identify loci contributing to variance in lifespan. While less conservative, this threshold is more stringent than a previously reported method (Wright et al. 2022). We report 2 LOD support intervals (‘2LOD SI’), corresponding to a 2 LOD drop around each peak position, about each peak marker identified in whole-genome scans.

### QTL effect size estimation and variance explained

Best linear unbiased predictors (BLUPs) and standard errors were calculated for all QTL using the scan1blups() function in “rqtl2” on meta-traits without the use of additive covariates, as above. The fraction of phenotypic variance explained by each QTL was calculated using the LOD score at the sentinel genotyping marker via the formula (Broman and Sen 2009):

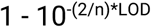

Where n is the number of observations (i.e. animals) and LOD is the LOD score of the peak marker at each QTL. In specified instances involving low sample sizes, LOD scores from the sentinel SNP identified in variant association mapping, rather than the sentinel genotyping marker, was used to estimate the fraction of variance explained by a QTL.

### Variant association / fine mapping of QTL

Variant association mapping was conducted within a 2LOD interval around the peak position of each QTL. Fine mapping was performed via the scan1snps() function in “rqtl2” without additive covariates, as detailed in the section *Additive whole-genome scans of meta-traits* above. Variant and gene SQLite datasets used in this analysis are available at the “rqtl2” user guide website: https://kbroman.org/qtl2/assets/vignettes/user_guide.html.

### Correlation of haplotype effects across trait clusters

We used 2LOD support intervals around QTL to determine which QTL from meta-traits overlapped previously detected, reproducible lifespan QTL on chromosomes 12 and 18 from meta-trait C_all_252 (DRiDO and meta-lifespan), and chromosome 16 from meta-trait C_all_216 (Shock study lifespan).

For each lifespan QTL, the top 1% of significant SNPs from variant association mapping were identified. The nearest genotyping markers to each of these SNPs were defined as the “focal markers” for each lifespan QTL. The set of unique focal markers at each lifespan QTL were then used to compute Pearson correlations between effect sizes at lifespan QTL and overlapping QTL. Sets of highly correlated haplotype effects (p < 1*10^-10^) were prioritized for downstream analysis.

### scRNA-seq expression in human protein atlas dataset

QTL that both overlap lifespan QTL and have haplotype effects correlated with the effects of the lifespan loci were considered potential lifespan loci. The meta-traits in which these QTL were detected were used to determine cell types of interest that could be useful for refining lists of candidate genes within the chromosome 12 and 18 confidence intervals. All genes within the 2LOD support intervals of the lifespan QTL on chromosomes 12 and 18 were recorded, and the expression levels of these genes within relevant cell types were examined using single-cell RNA seq (sc-RNAseq) data from the Human Protein Atlas (Uhlén et al. 2015; Karlsson et al. 2021).

sc-RNAseq data were downloaded from the Human Protein Atlas website at: https://www.proteinatlas.org/humanproteome/single+cell/single+cell+type/data#cell_type_data

### Correlation of haplotype effects at cardiac eQTL

An external transcriptomic dataset derived from the cardiac tissue of DO mice was used to identify *cis*-eQTL overlapping with QTL detected in meta-analysis of aorta phenotypes (Gerdes Gyuricza et al. 2022). This dataset was accessed via the Jackson Laboratory QTL Viewer website at https://churchilllab.jax.org/qtlviewer/JAC/DOHeart. For each aorta QTL, genes with *cis*-eQTL and starting positions within 2Mb of the sentinel marker were recorded. Pearson correlation coefficients were computed between the haplotype effects (BLUPs) at the peak positions of the aorta QTL and any nearby *cis*-eQTL. BLUPs were not available at each genomic marker within the external dataset, so correlations were based only on the haplotype effects at peak markers rather than across all significantly associated markers within the 2LOD support intervals of the QTL. Correlation coefficients with p < 0.1 were considered significant in our analysis.

This procedure was performed using proteomic data from the same external dataset, but no significant *cis*-pQTL were detected within 2Mb of the aorta QTL from this study.

### Open field and frailty assays

We implemented an open field system that would be compatible with the varied coat colors of DO mice. Details about this system, including instructions for assembly, use, and source code for the analysis of video data, are provided in this GitHub repository: https://github.com/graham-calico/OpenFieldAssay. For the phenotypes reported here, videos were collected and analyzed for 10 minutes after the mouse was placed in the box. Phenotypes were generated by the openFieldAnalysis.py script using the following non-default command-line options: “-b best -- adjust_fps”.

All MFI data analyzed here are from (Ruby et al. 2023), as are many of the analyzed DFI data. Additional DFI measurements reported here were collected as described by (Ruby et al. 2023).

### Aortic dissection

There were a total of 138 Diversity Outbred mouse aortas collected across 9 aged cohorts: 11 mo (n=26), 14 mo (n=24), 17 mo (n=15), 20 mo (n=11), 23 mo (n=8), 25 mo (n=5), 29 mo (n=20), and 32 mo (n=29). The whole aorta was removed, stripped of adherent tissue, then cut into four specific segments labeled with different colored dye for identification. Four segments: the ascending aorta, was cut from the base of the aorta connected to the heart and separated before the brachiocephalic trunk; the aortic arc, start at the brachiocephalic trunk and cut at the left subclavian artery; the descending aorta, start at the end of the left subclavian artery and cut at 6mm then split into two (proximal & distal) 3mm aortic segments. Each tissue was embedded into paraffin blocks and sectioned for Hematoxylin & Eosin (H&E) stain, Verhoeff van Gieson (VvG) stain, and Mason Trichrome (Trichrome) stain.

### Lung histology

Female C57BL/6J and DO mice were obtained from the Jackson Laboratory and allowed to acclimate for 2-4 weeks. Lungs were perfused with sterile PBS and inflated with 1.5% low melting point agarose (Invitrogen) at a pressure of 25cm H_2_O. Lungs were excised and fixed in 10% neutral buffered formalin for 24 hours at room temperature. Left lobes were processed for histology (serial H&E and Trichrome collagen stains at 6-8 positions in the lobe with a 200 um distance between positions). All procedures used in our studies were reviewed and approved by the Calico Animal Care and Use Committee.

### Micro-CT imaging of lung tissue

To assess lung structure, mice were anaesthetized with isoflurane (1-3%) in medical air. Isoflurane levels were adjusted to maintain a breathing rate of 50-60 breath per minute. Focused lung scans with respiratory gating were acquired with a MILabs micro-CT system (U-CT, MILabs, Utrecht, Netherlands) using the following settings: mouse ultra focus protocol, X-ray voltage 60 kV, current 160 µA, exposure 50 ms, binning of 2X2, 0.5 degrees step angle, 6 projections per angle. Micro-CT images were reconstructed using CTRecon software (Gremse-IT GmbH, Germany) at 50 µm isotropic resolution with retrospective respiratory gating. C57BL/6J and the DO mice were imaged once. Radiation delivered per scan was less than 1 Gy and imaging time is approximately 6 min.

Lungs were automatically segmented using a machine learning (U-Net) model trained on manual segmentations of lungs (Ronneberger, Fischer, and Brox 2015). After segmentation, lung masks were registered (rigid followed by affine) to a reference atlas using Elastix (Version 5.2) (Klein et al. 2010; Shamonin 2013). Masks for different lobes (Left, right cranial, right middle, right caudal and right accessory) were propagated to the original image space and used for lobe specific analysis. Image intensity thresholds were defined for fibrosis or inflammation (> −250 HU) and emphysema (< −750 HU). Different lung parameters like lung volume, high signal intensity volume “fibrosis volume”, emphysema volume, percentage of lung affected by fibrosis or emphysema were computed for each lobe.

### Oxygen transfer measurements

Carbon monoxide diffusion measurements is a commonly used technique to measure lung function (transfer rate from alveoli to the blood stream). A surrogate method has been developed to estimate oxygen transfer rates based on changes in blood oxygen saturation (spO2) when animals are breathing gas mixtures containing 21% or 7% oxygen. Mice were anesthetized with 1.5% isoflurane, transferred to a heating pad to maintain body temperature at 37°C and maintained on 0.75% isoflurane. A rectal temperature probe was used to monitor body temperature and pulse oximeter clip (MouseOxPlus, Starr Life Sciences, PA, USA) positioned on the thigh on which hair removal had been performed on the previous day using a depilatory cream. A breathing pad was attached to calculate the breathing rate. Mice were subject to hypoxia at 7% oxygen + 93% nitrogen for 2 minutes followed by normoxia at 21% oxygen + 79% nitrogen for 2 minutes. This exposure pattern was repeated three times. Temperature, breathing rate, heart rate and oxygen saturation were being recorded continuously during the experiment. Linear fits were performed for blood oxygen saturation during the switch from hypoxia to normoxia. We define the slope of this fit as oxygen transfer rate.

### Lung mechanics measurements

Mechanical lung parameters were measured with the flexiVent (FX, SCIREQ, Montreal, QC, Canada) system. Mice were anesthetized with ketamine (70 mg/kg) and xylazine (8 mg/kg). A 20G cannula was inserted intratracheally and connected to the flexiVent while the mice were placed in a supine position. Mice were mechanically ventilated with a tidal volume of 10 mL/kg at a breathing frequency of 150 breaths/min and with a positive end-expiratory pressure of 3 cmH_2_O. After checking for leaks and once the ventilation was underway, mice were paralyzed by injecting of rocuronium (2 mg/kg) intraperitoneally, to avoid spontaneous breathing during the procedure. Lung mechanics measurements were made using the flexiVent system’s automated algorithms and were repeated three times.

A Deep Inflation maneuver inflated the lungs to maximum capacity at a pressure of 30 cmH_2_O, maintained over a time period of 3 s, to measure the inspiratory capacity (IC). To calculate total respiratory system resistance (R_rs_), compliance (C_rs_), and elastance (E_rs_), a single-frequency (SnapShot-90) perturbation was used in alignment with mice respiratory rate and tidal volume. Impedance based measurements like newtonian resistance (R_n_), tissue damping (G), and tissue elastance (H) was obtained by running a broadband (Quick Prime-3) forced oscillation perturbation which uses a range of frequencies above and below the respiratory rate. Inspiratory capacity (A), curvature of the deflating PV loop (K), and quasi-static compliance (C_st_) were measured using stepwise pressure-controlled pressure–volume (PVs-P) loops. The forced expired volume over 0.1 s (FEV0.1) was obtained by performing the The negative pressure-driven forced expiratory (NPFE) maneuver by inflating the mouse lungs to a pressure of 30 cm H_2_O for 1 s, and holding that pressure for a period of 2 seconds followed by connecting the animal’s airways to the negative pressure reservoir (− 55cm H_2_O) for 2 seconds. Data was excluded if the model fit was poor (coefficient of determination less than 0.9) for every model.

### Histological image preprocessing and ROI extraction

We developed automated computational pipelines for quantitative analysis of histological images, tailored to specific staining techniques and tissue types. These pipelines facilitate efficient handling of large whole-slide images, extraction of regions of interest (ROIs), and quantification of morphological and structural features pertinent to our study. The detailed methodology for image preprocessing and ROI extraction is extensively described in our separate publication (Lefebvre and Mullis 2025); here, we briefly summarize the key steps and specific analyses performed. Whole-slide images were downsampled to reduce computational load while preserving sufficient detail for accurate region detection. Optical density (OD) images were computed to enhance contrast between stained tissue and background. Background masking and adaptive thresholding were applied to segment tissue regions effectively. Morphological operations refined the tissue masks, and connected-component analysis identified distinct ROIs defined by bounding boxes. The downsampling scaling factor was recorded to ensure precise mapping between the downsampled images and the original resolution.

### Analysis of aortic elastic laminae in VVG-stained images

We quantified elastin structures within the aortic wall from Verhoeff–Van Gieson (VVG)-stained images, assessing parameters such as elastin layer thickness, continuity, waviness (tortuosity), and the number of elastin layers. The pipeline for VVG analysis is extensively detailed in our separate publication (Lefebvre and Mullis 2025). Here, we briefly summarize the pipeline. High-resolution images were processed to extract ROIs based on predefined bounding boxes. Color deconvolution isolated elastin staining in OD space, separating VVG stain components and removing artifacts. Elastin masks were created by thresholding the OD image, followed by morphological operations to refine the mask. Skeletonization reduced fibers to their centerlines for structural analysis. A network representation was constructed, classifying nodes as tip points, branch points, or regular points, and metrics like network complexity were calculated. Elastin thickness was measured using a distance transform sampled at skeleton pixels, providing a pixel-wise representation across the tissue. Waviness was quantified by calculating tortuosity along fiber paths, comparing curved distances to straight-line distances. The number of elastin layers was determined by analyzing the elastin mask and counting distinct components in horizontal and vertical directions.

### Quantification of aortic cellular metrics in HE-stained images

In Hematoxylin and Eosin (HE)-stained images, we quantified cellular metrics by detecting and segmenting nuclei to assess cellularity. Whole-slide images were read using the tifffile library, and ROIs were extracted based on predefined bounding boxes. Color deconvolution separated the hematoxylin and eosin stain contributions. A background mask excluded non-tissue areas through intensity thresholding, color ratio calculations, Gaussian filtering, and morphological operations. Tissue regions were identified by thresholding OD images. Nuclei detection and segmentation involved thresholding the smoothed OD channels, combining masks, and using distance transforms and watershed segmentation. Small objects were removed to eliminate noise, and connected components were labeled to identify individual nuclei. Cellular metrics such as the number of cells per tissue area and cell area per tissue area were calculated, providing insights into cell density and tissue occupancy by cells.

### Analysis of aortic wall metrics in trichrome-stained images

We quantified collagen density within the aortic wall, wall thickness, external collagen lining thickness, and perimeters of the aorta in trichrome-stained images. Whole-slide images were read, and ROIs were extracted. Color deconvolution separated the staining components specific to trichrome staining. Collagen masks were generated by thresholding the appropriate OD channel after Gaussian filtering. An aortic wall mask was created by summing OD channels, applying Gaussian filtering, thresholding, and refining with morphological operations. The collagen mask and aortic wall mask were combined to identify collagen within the aortic wall, and the fraction of collagen was calculated by the overlap area. Thickness measurements were obtained using distance transforms and skeletonization of the masks, providing detailed measurements across the tissue. Perimeters were calculated using connected-component analysis and morphological operations. Statistical analyses of thickness maps included metrics such as mean, standard deviation, coefficient of variation, and skewness.

### Quantification of alveolar structures and nuclear metrics in HE-stained lung images

In HE-stained lung images, we quantified alveolar structures and nuclear metrics to assess lung tissue architecture and cellularity. Images were downsampled for computational efficiency. A tissue mask was generated from grayscale images using adaptive thresholding and morphological operations. Alveolar structures were detected using multi-scale Laplacian or Gaussian filters, and alveolar centers were identified from local maxima in the distance-transformed image. Alveolar radii were measured by averaging distances from centers to the nearest tissue borders. Nuclei were detected via color deconvolution to extract the hematoxylin channel, followed by Gaussian filtering and thresholding. Local maxima identified individual nuclei, and nearest neighbor distances provided information on nuclear distribution and density. Cluster analyses were conducted to assess nuclear clustering based on proximity and density.

### Quantification of collagen content in trichrome-stained lung images

We quantified collagen content in trichrome-stained lung images by generating tissue masks and isolating collagen-specific stains via color deconvolution. Collagen masks were created by thresholding the appropriate OD channel after Gaussian filtering. Vessel regions were excluded using a vessel mask generated through image filtering and thresholding. Quantitative metrics were derived by calculating the areas occupied by tissue and collagen within the lung tissue mask. Ratios representing the proportion of lung tissue occupied by collagen were calculated by dividing the collagen area by the sum of tissue and collagen areas.

### Histological image data analysis and validation

Quantitative metrics from each pipeline were compiled into structured data formats and saved as CSV files for statistical analysis. A thread-safe writing mechanism ensured data integrity during concurrent processing. Optional validation images were generated to facilitate visual inspection of processing results, including overlays of masks and thickness maps on original images. All image processing operations utilized established libraries such as NumPy, SciPy, and scikit-image. Our custom-built histo-tools library was used for stain deconvolution, providing robust and efficient implementations of the required algorithms. This approach ensured consistent and efficient processing across multiple subsamples and slides, enabling large-scale quantitative analysis of histological images.

### In vivo assays and husbandry

All research was performed as part of Calico Life Sciences LLC (South San Francisco, CA) AAALAC-accredited animal care and use program. All research and animal use in this study was approved by the Calico Institutional Animal Care and Use Committee.

## Acknowledgments

We would like to thank Gary Churchill, Andrea Di Francesco, Anurag Sethi, and Jinguo Huang for their thoughtful feedback on the project and manuscript.

## Funding

This study was funded by Calico Life Sciences LLC, South San Francisco, CA.

### Data Availability

Code for performing HE genetic regression is available at GitHub: https://github.com/calico/HE-regression

Hardware designs, code, and ML models for open field analysis are available at Github: https://github.com/graham-calico/OpenFieldAssay

Supplemental tables and data files are available at Dryad. DOI: https://doi.org/10.5061/dryad.sj3tx96gd

Dryad review link: http://datadryad.org/share/-ubFH2a8wDiIbFH3Q8r75I7DElWAERn7PfExYgpB95c

### Author Contributions

JGR conceptualized the study. MNM and JGR developed the methodology associated with the genetic analyses. MNM performed genetic and statistical analyses and data curation. AEL analyzed histological images. AL, JZ-S and CZ collected aortic tissues. AG and JR collected and processed lung functional/histological data; KS, MS, and JR collected and processed lung microCT data. FS and JGR implemented the open field assay: AL collected and JGR processed those data. KW and AR provided technical support and training. MNM created the figures; MNM and JGR wrote the manuscript.

## SUPPLEMENTARY FIGURES

**Figure S1.**
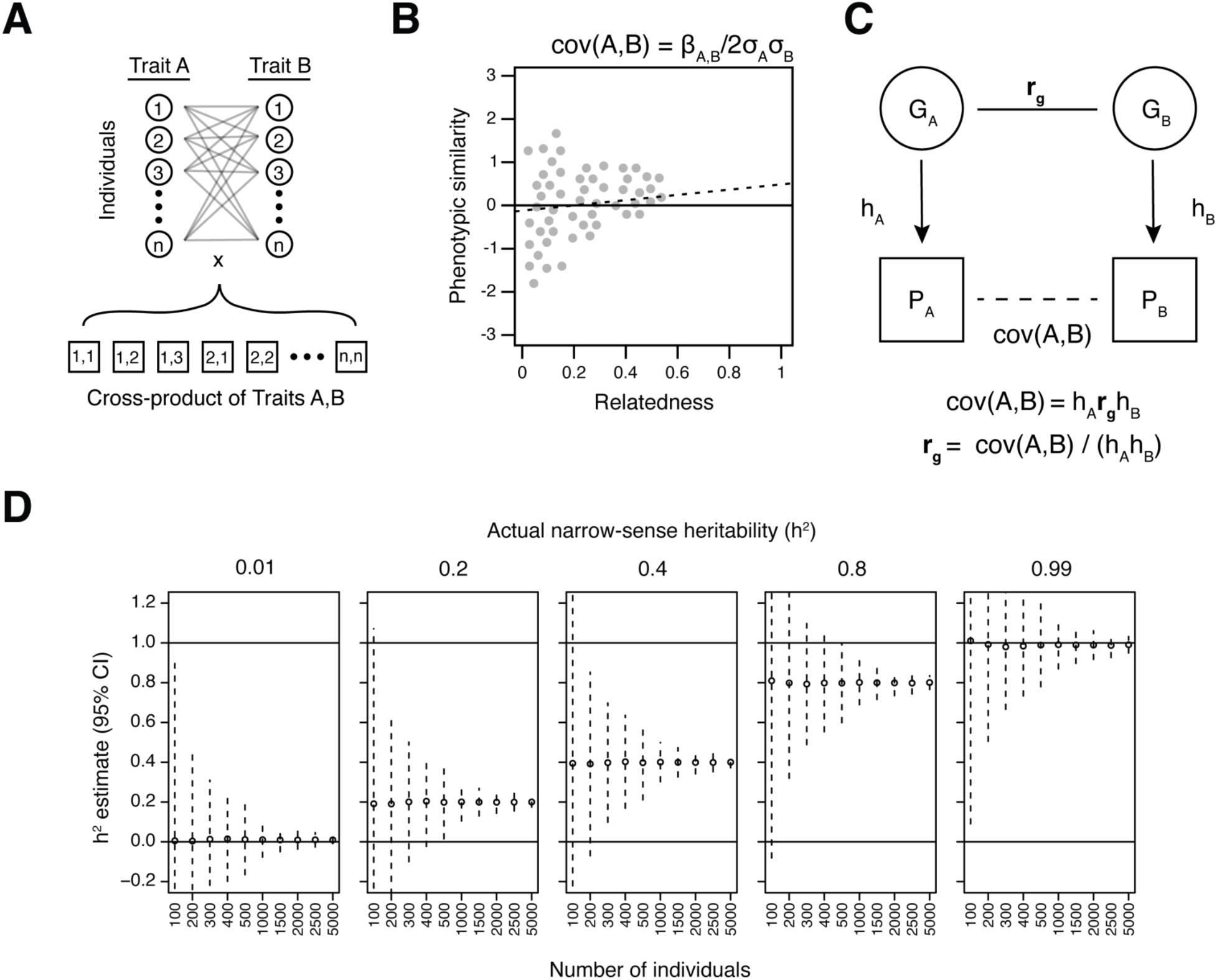
Haseman-Elston regression for estimation of genetic correlations (*r_g_*). **A**, Individual phenotype data is collected for two traits (‘A’ and ‘B’). Distributions of traits are z-score normalized and a cross product is taken by multiplying each measurement of trait A by each measurement of trait B. The result is a set of pairwise phenotypic similarity scores from all combinations of individuals. **B,** Phenotypic similarity scores for each pair of individuals are regressed on pairwise kinship values. The coefficient of the regression can be used to determine the covariance of traits A and B. **C,** A structural equation model relates the covariance of traits A and B as a function of relatedness to the genetic correlation (*r_g_*) of the two traits. The h^2^ and covariance of traits A and B must be known in order to solve for *r_g_*. **D,** Haseman-Elston *h*^2^ estimates for simulated phenotypes. Mean Haseman-Elston *h^2^*estimates and 95% confidence intervals of 1,000 randomly simulated traits at different sample sizes. Actual *h^2^* of the simulated traits is shown at the top of each panel.

**Figure S2.**
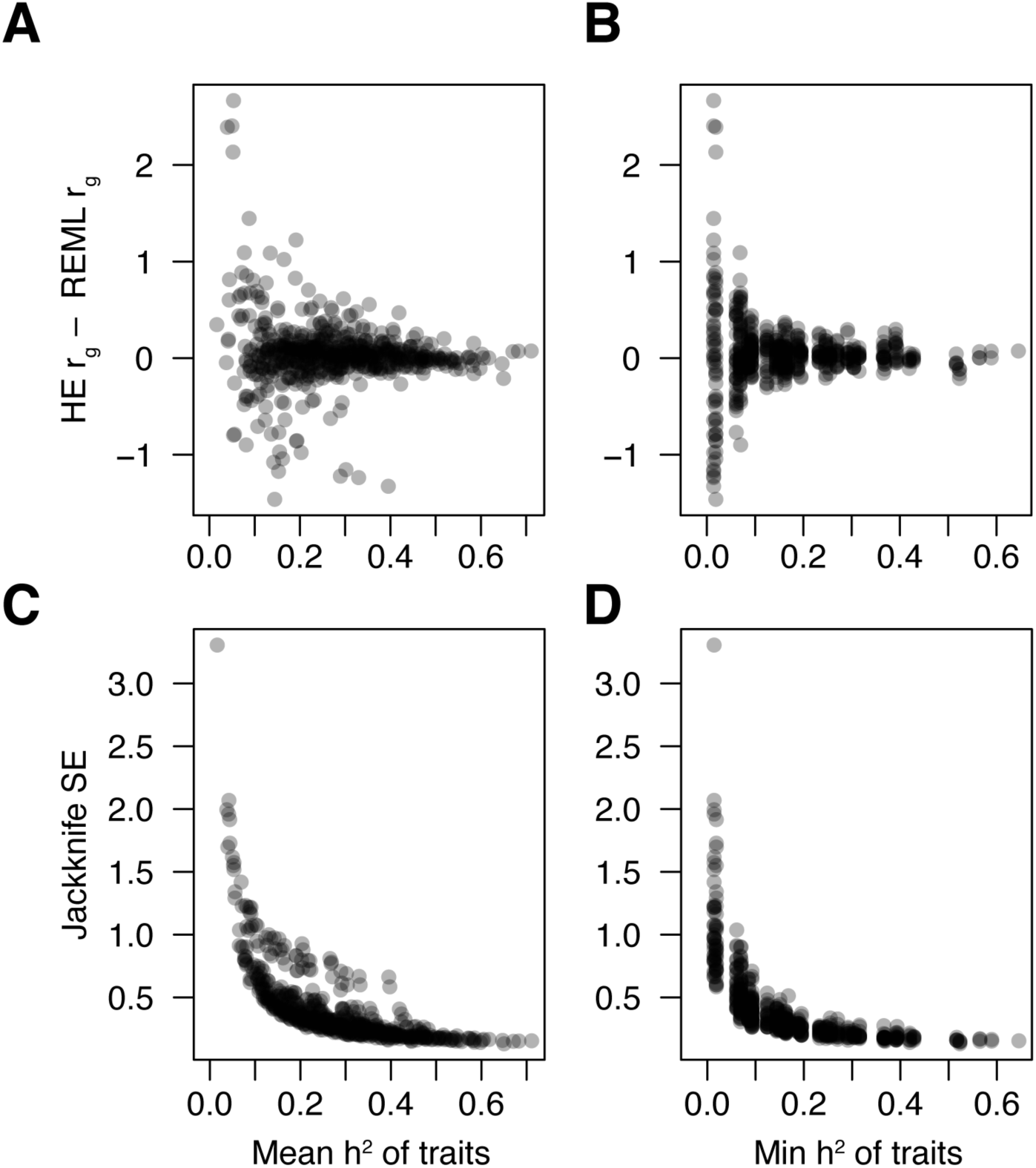
Differences between Haseman-Elston and REML *r_g_* estimates are driven by *h*^2^ of trait pairs. **A**, Differences in pairwise r_g_ estimates between Haseman-Elston regression and REML as a function of the mean *h*^2^ of the traits being compared. **B,** Differences in pairwise *r_g_* estimates between Haseman-Elston regression and REML as a function of the minimum *h*^2^ of the traits being compared. **C,** Standard error of Haseman-Elston based *r_g_* estimates as a function of the mean *h*^2^ of the traits being compared. **D,** Standard error of Haseman-Elston based *r_g_* estimates as a function of the minimum *h*^2^ of the traits being compared.

**Figure S3.**
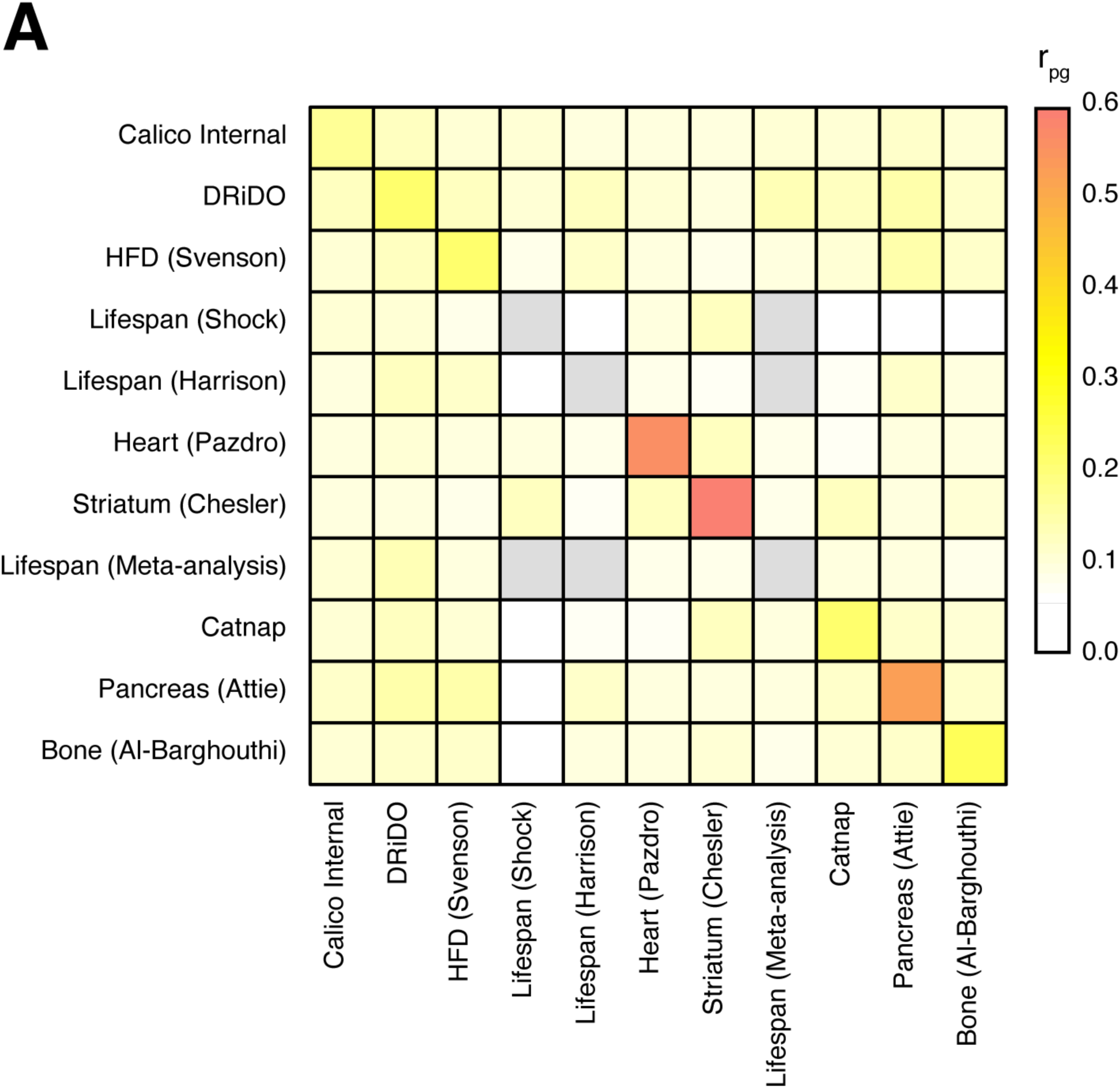
Mean inter- and intra-study *r_pg_* estimates. Mean *r_pg_* estimates among traits from each study. Values derived from a single trait pair are represented by gray boxes. ‘Calico internal’ traits comprise open field, frailty, lung, and aorta traits, including histology-based phenotypes.

**Figure S4.**
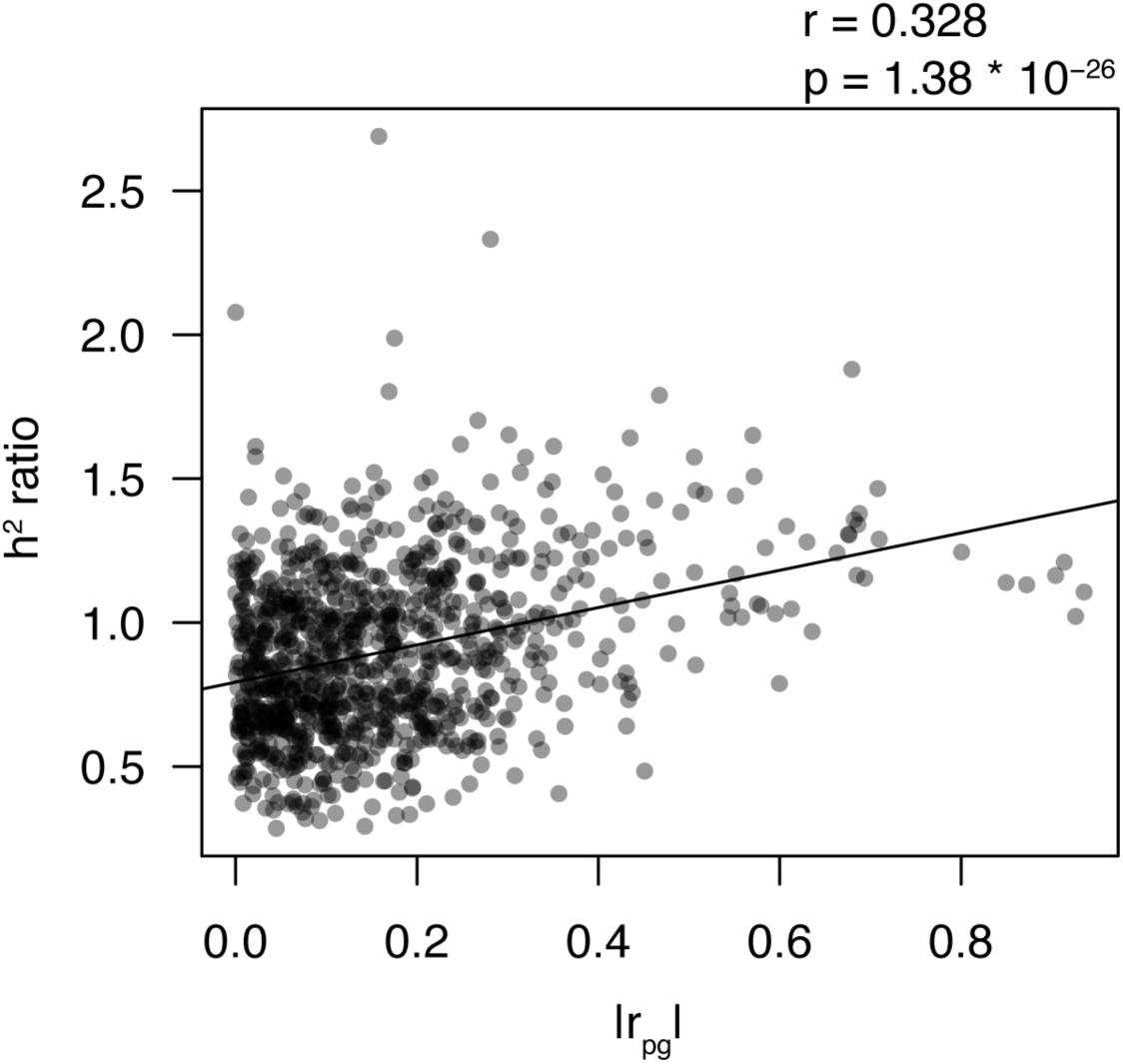
Heritability of aggregated trait pairs increases relative to individual heritability estimates as a function of r_pg_. Scatterplot of the h^2^ ratio for 1,000 random pairs of traits as a function of the absolute value of their r_pg_. The h^2^ ratio for traits A and B is defined as the h^2^_meta_ / mean(h^2^_A_,h^2^_B_), where h^2^_meta_ is the heritability of a meta-trait constituting traits A and B, and h^2^_A_ and h^2^_B_ are the individual heritabilities of traits A and B. The Pearson correlation coefficient and p-value between the h^2^ ratio and |r_pg_| are reported.

**Figure S5.**
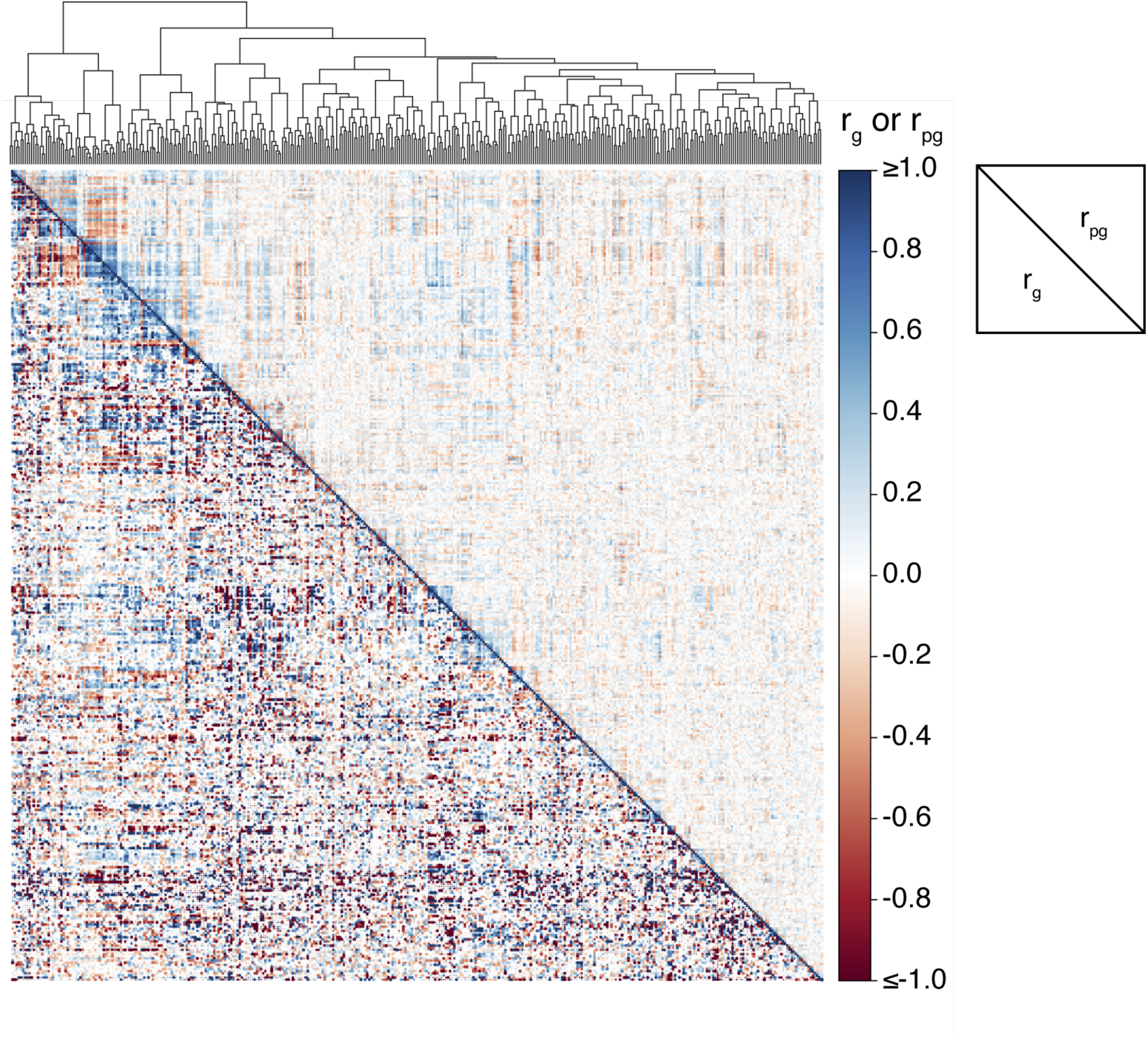
Genetic correlations among 383 meta-traits. Hierarchically clustered heatmap of r_g_ (*lower triangle*) and r_pg_ (*upper triangle*) estimates among 383 DO meta-traits. The accompanying dendrogram shows euclidean distances among meta-traits based on their r_pg_ estimates.

**Figure S6.**
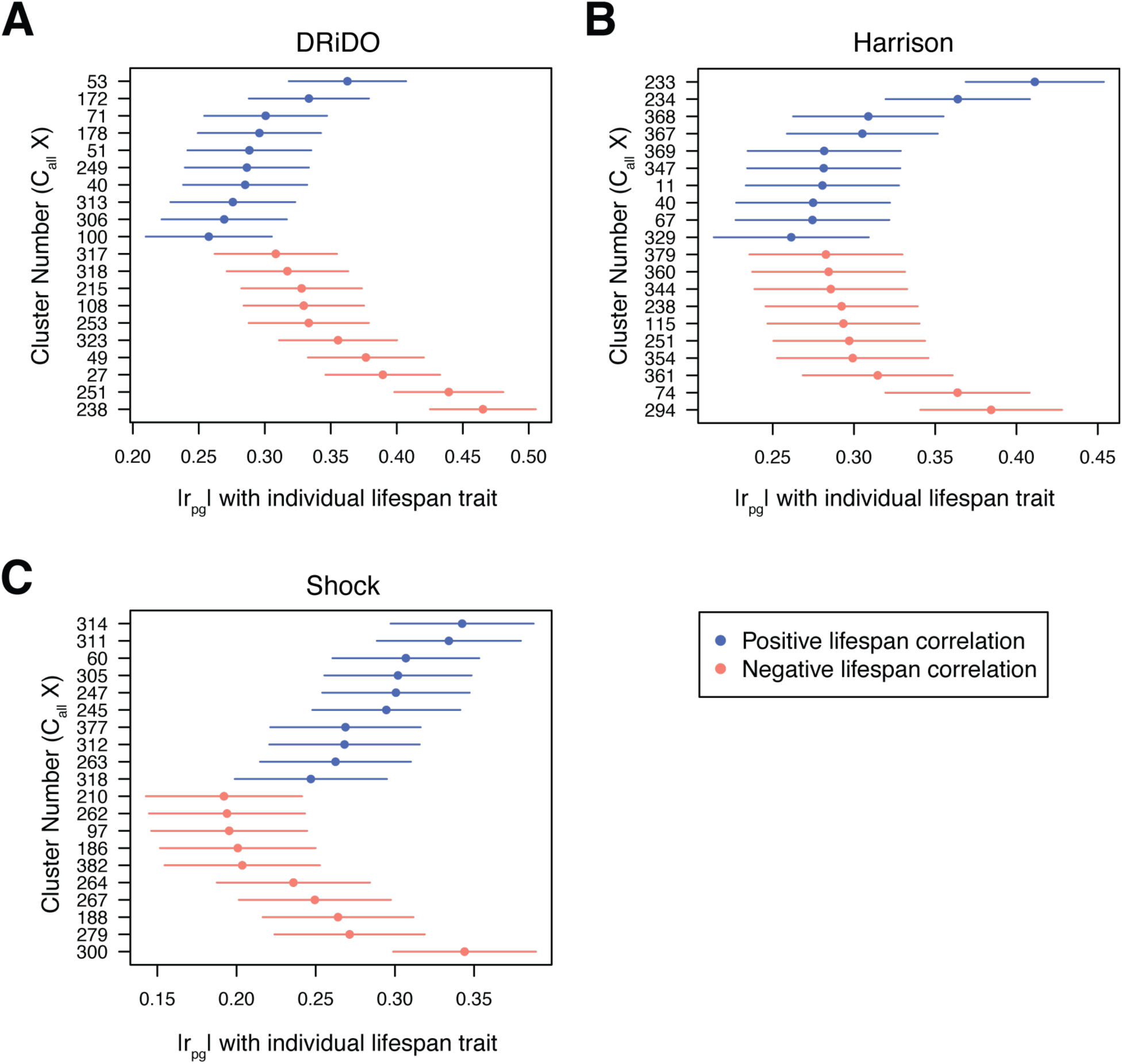
r_pg_ estimates between individual lifespan studies and meta-traits. The strongest positive and negative r_pg_ estimates between meta-traits and lifespan data measured in **A,** the DRiDO study, **B,** the Harrison study, and **C,** the Shock study. Error bars correspond to the standard error of each r_pg_ estimate. Meta-trait number (cluster number) is shown on the left hand side of the plot. The traits comprising each cluster can be found in **Table S7**.

**Figure S7.**
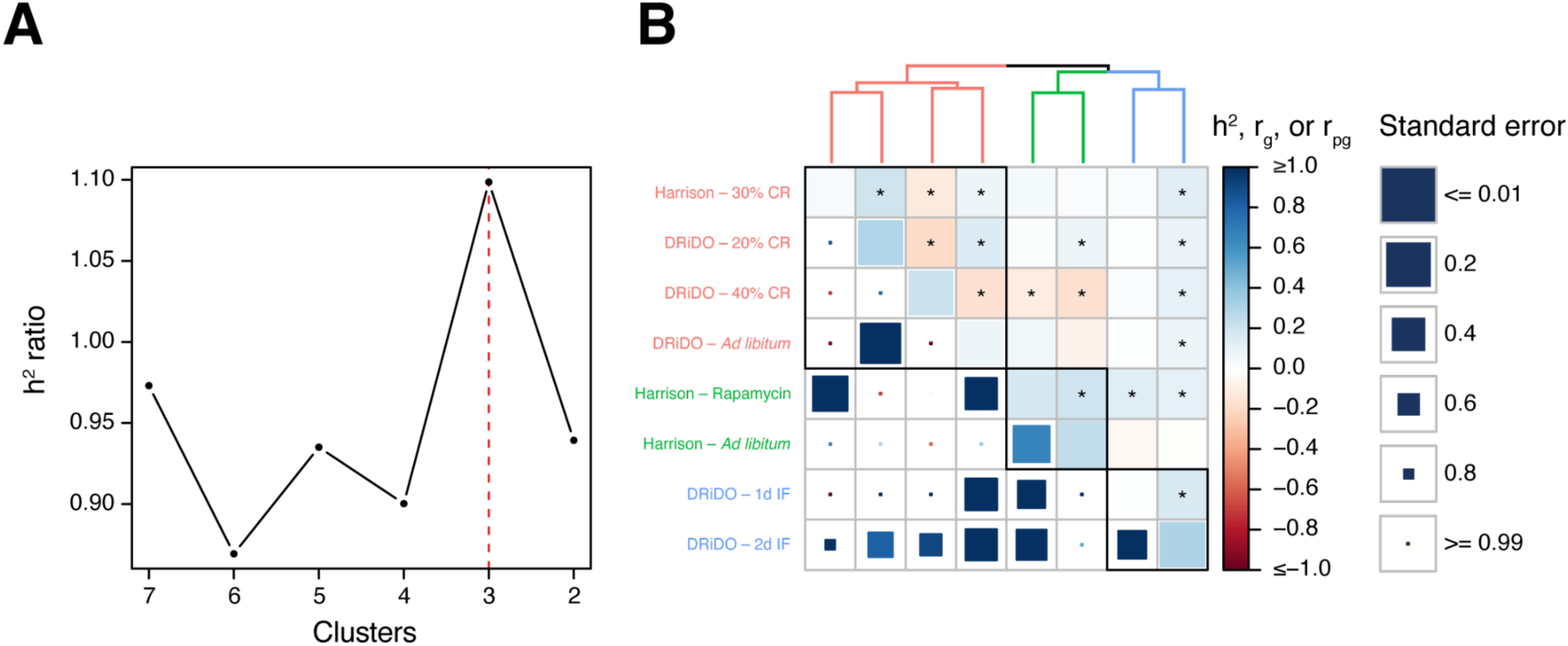
Genetic correlations among diet-specific lifespan measurements. **A**, Plot of mean h^2^ ratios among clusters of diet- or drug-specific lifespan measurements at different levels of hierarchical clustering (k). **B,** Hierarchically clustered heatmap of r_g_ (*lower triangle*) and r_pg_ (*upper triangle*) among lifespan measurements within individual dietary or drug intervention groups. Statistically significant estimates after correcting for phenome-wide multiple testing are denoted by an asterisk.

**Figure S8.**
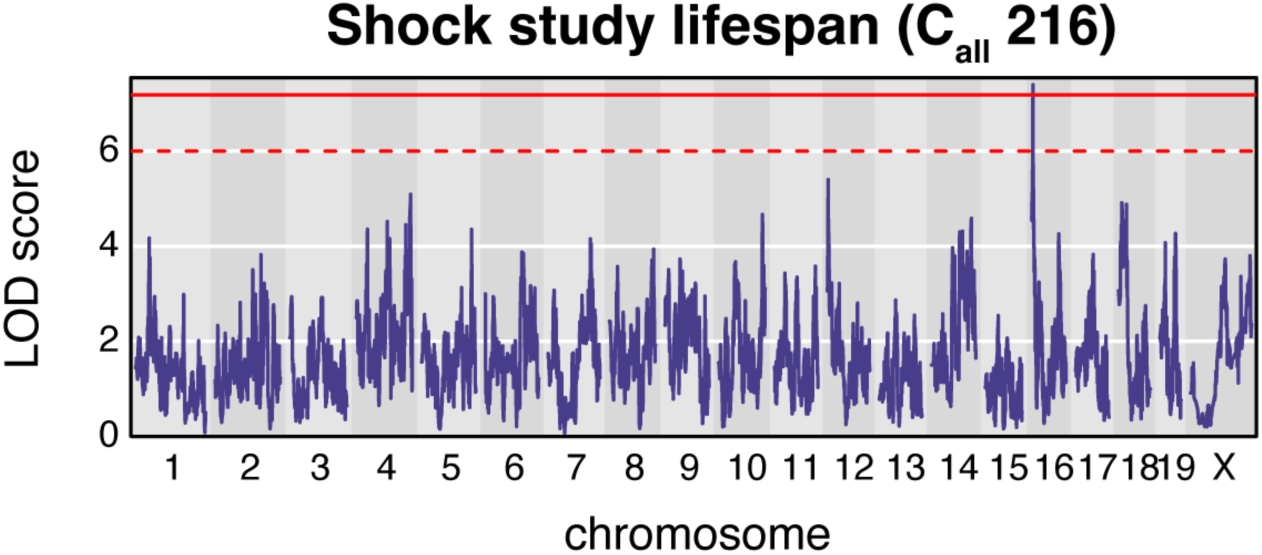
Additive whole-genome scan on lifespan measurements from the Shock study. Solid red lines indicate a trait-specific and permutation-based genome-wide significance threshold. Dashed red lines indicate a nominal significance threshold of LOD > 6.

**Figure S9.**
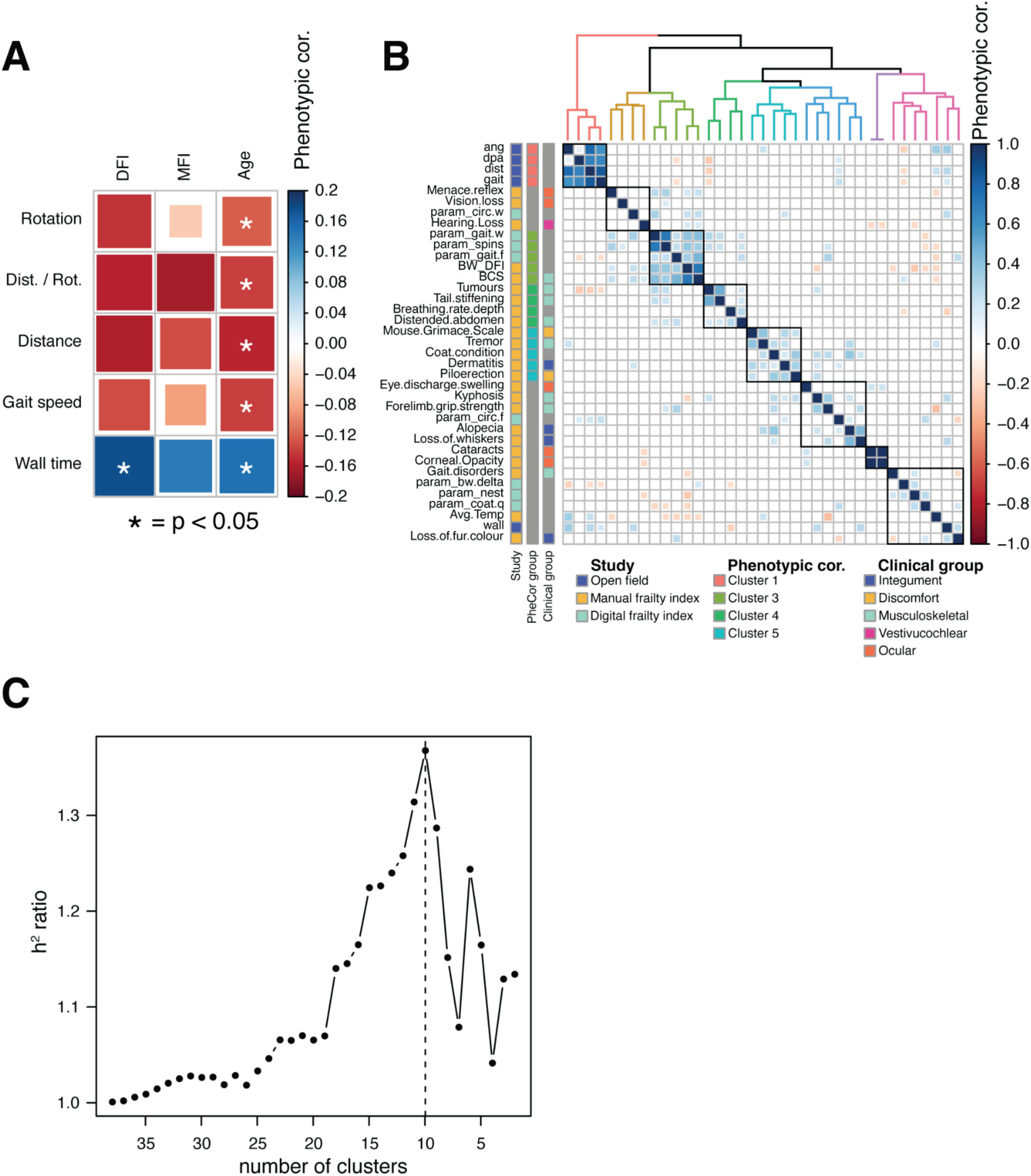
Phenotypic correlations among frailty traits. **A**, Phenotypic correlations between open field traits and mean digital frailty index scores (DFI), mean manual frailty index scores (MFI), and age. Statistically significant correlations are denoted by an asterisk. **B,** Phenotypic correlations among open field, DFI, and MFI traits. **C**,

**Figure S10.**
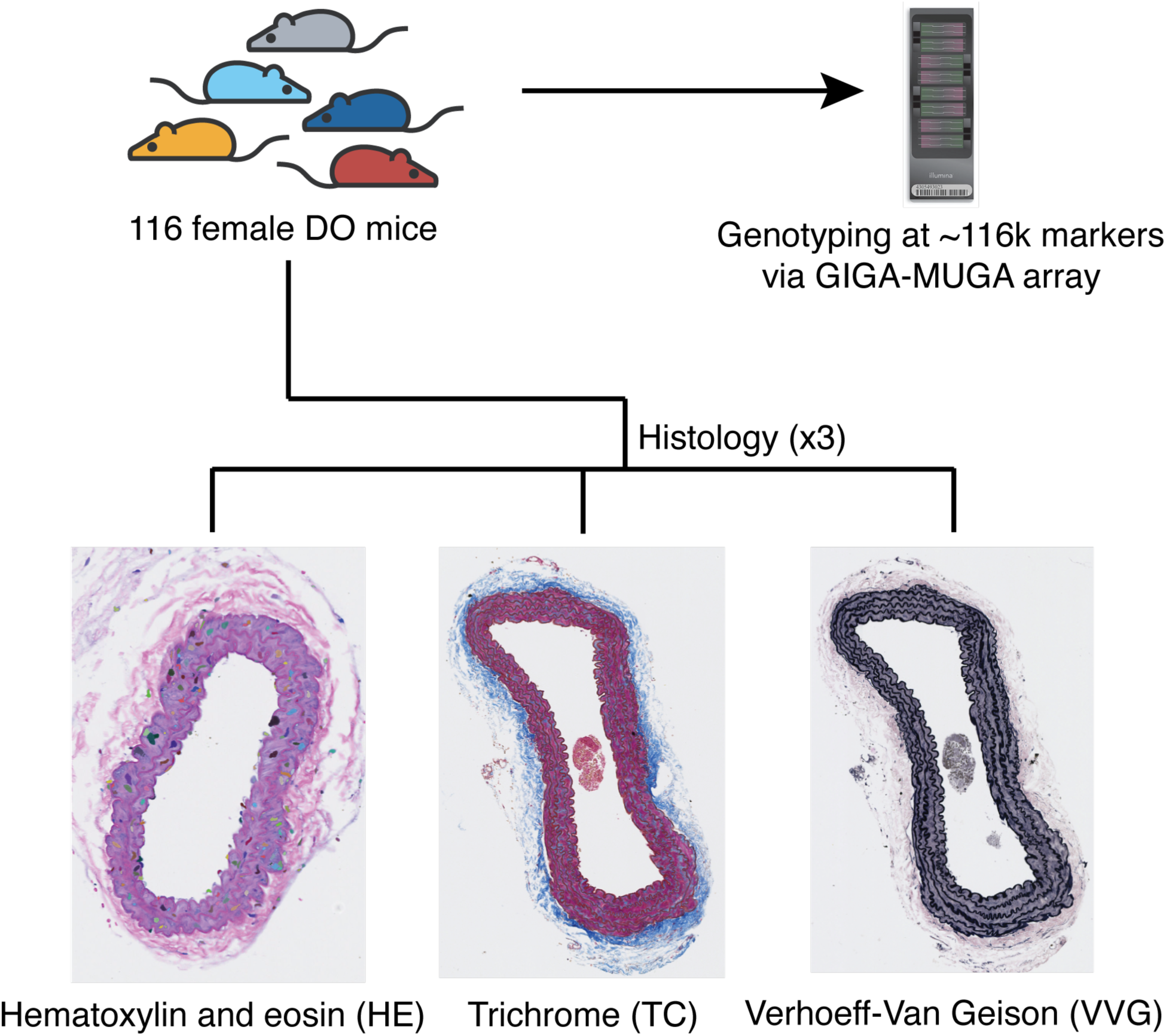
A histological analysis of aorta structure in DO mice. 116 DO mice were genotyped and their aortas were harvested. Tissues were sectioned, stained with HE, TC, or VVG, and imaged.

**Figure S11.**
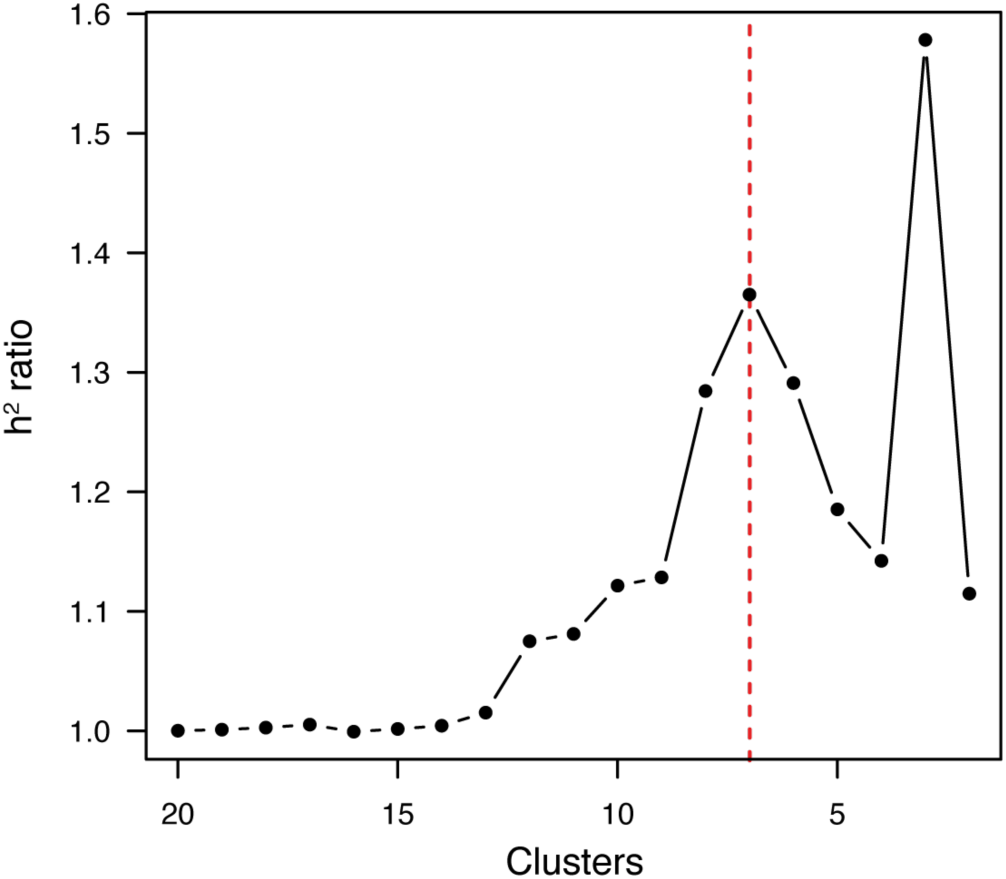
h^2^ ratio by clustering level for aorta phenotypes. Plot of mean h^2^ ratios among clusters of aorta meta-traits at different levels of hierarchical clustering (k).

**Figure S12.**
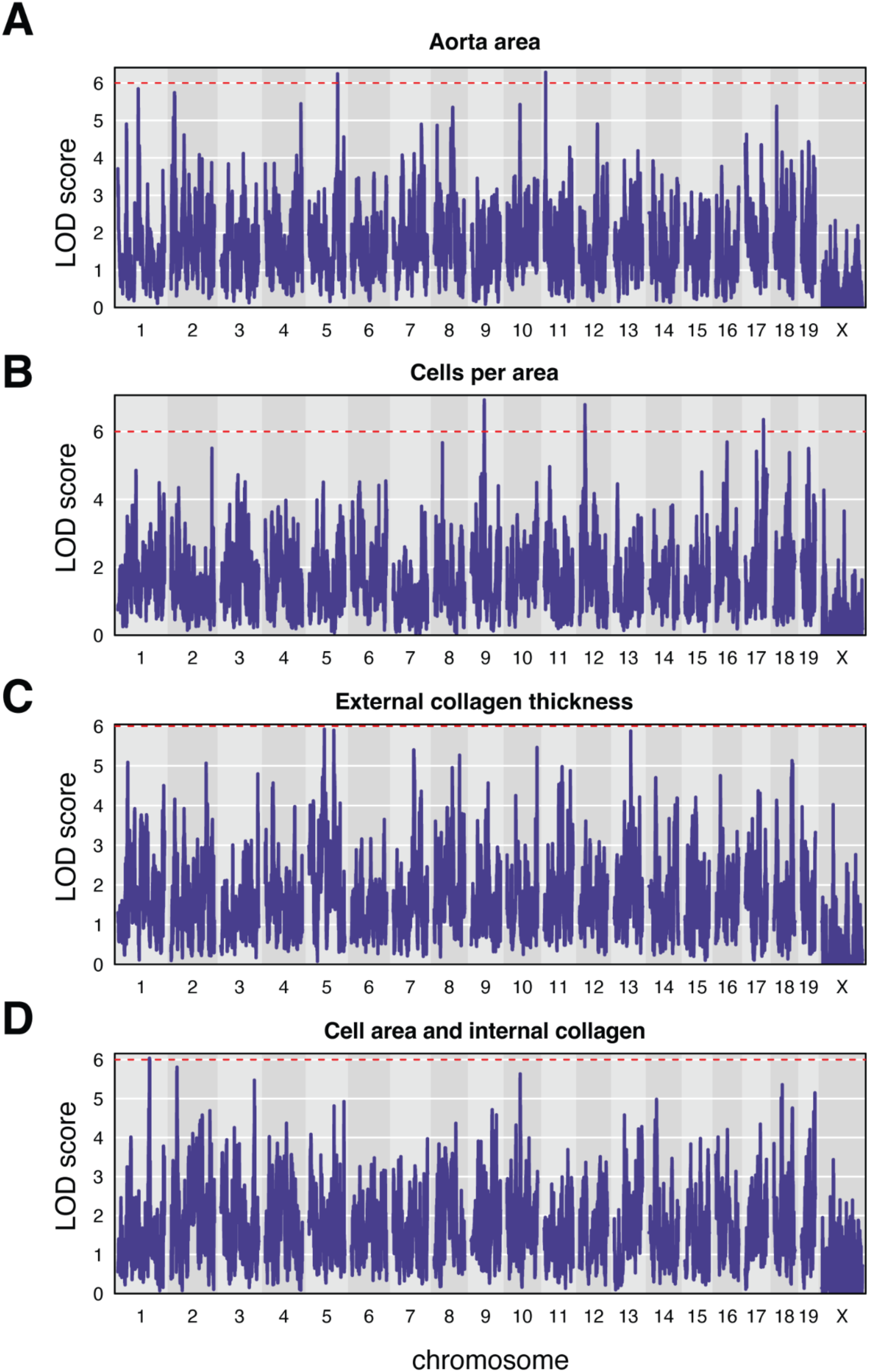
Genetic mapping of additional aorta meta-traits. Manhattan plots of additive whole-genome scans on aggregated aorta phenotypes: **A,** Aorta area. **B,** Cells per area. **C,** External collagen thickness. **D,** Cell area and internal collagen. Solid red lines indicate a trait-specific and permutation-based genome-wide significance threshold. Dashed red lines indicate a nominal significance threshold of LOD > 6.

**Figure S13.**
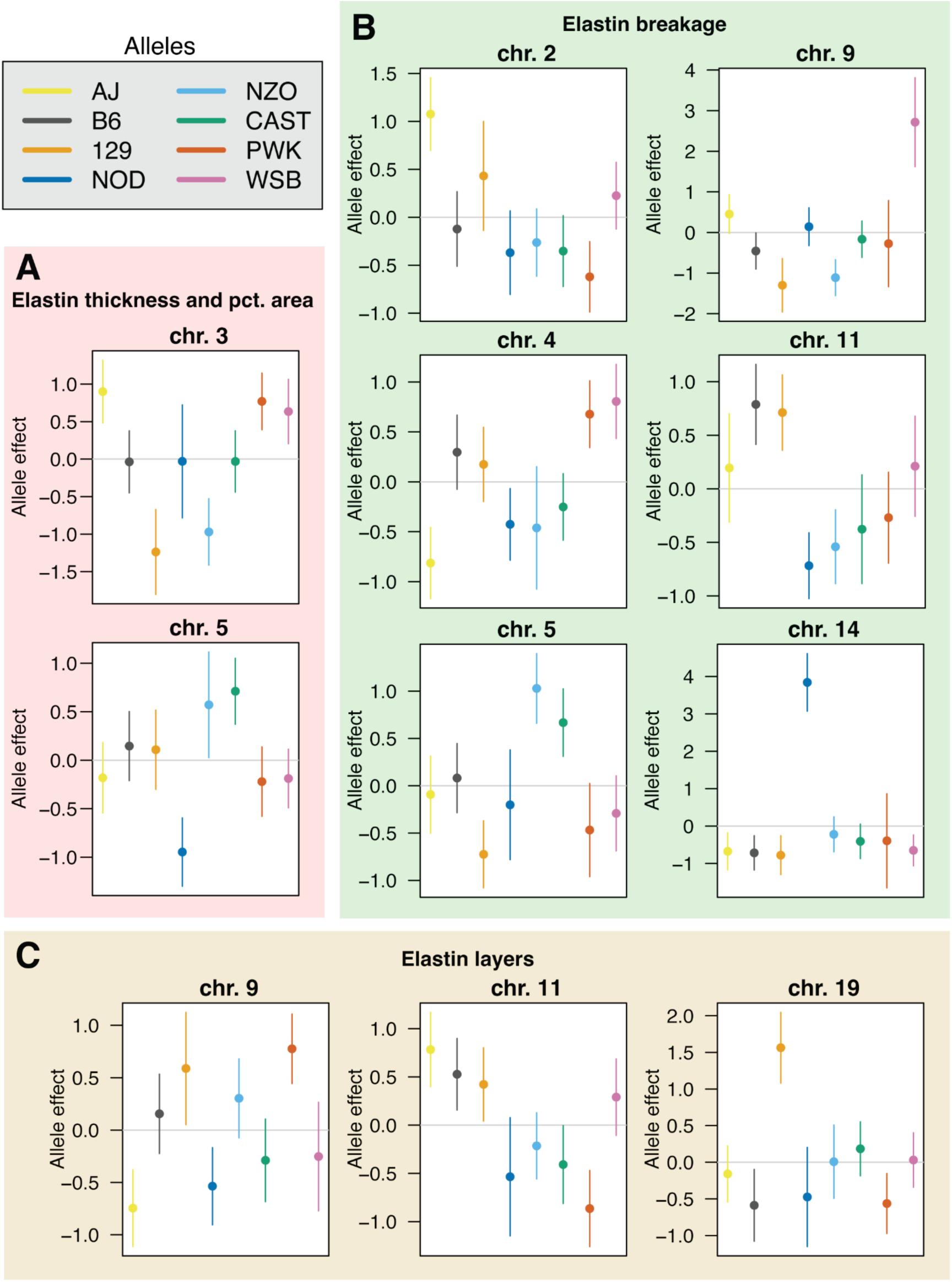
Haplotype effects of elastin QTL. Allelic effects (BLUPs) and corresponding standard errors of the loci detected in genome-wide scans of elastin meta-traits. **A,** Elastin thickness and percent area. **B,** Elastin breakage and tortuosity. **C,** Number of elastin layers.

**Figure S14.**
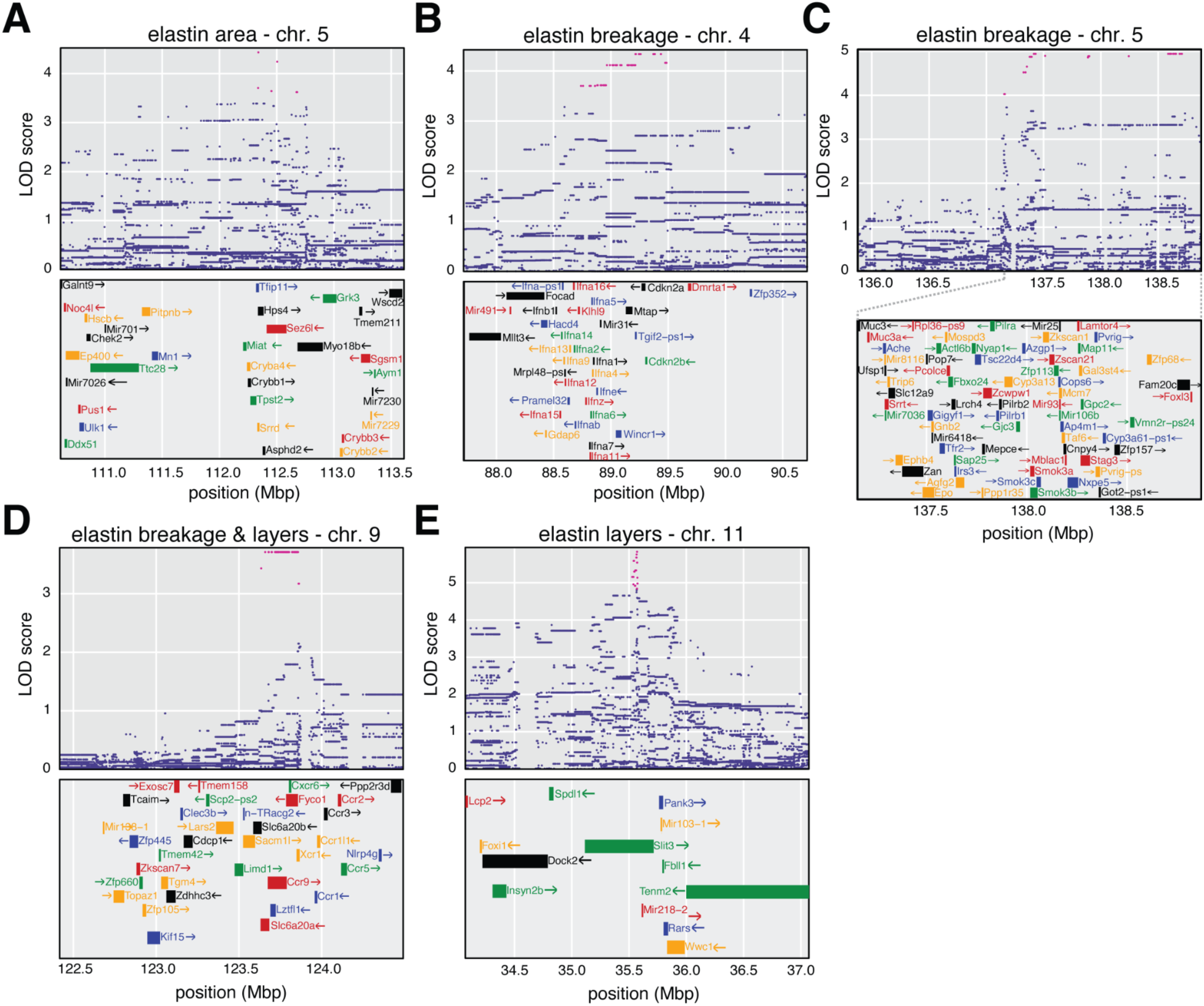
Finemapping of elastin QTL with correlated allelic effects in cardiac cis-eQTL. **A-E,** Variant association mapping plots for each elastin QTL at which the allelic effects were significantly correlated with at least one cis-eQTL in an external transcriptomic dataset from cardiac tissue in aging mice. (*Top*) The LOD scores for each variant within the 2 LOD support intervals around each locus in the meta-analysis. The most likely candidate SNPs are highlighted in pink. (*Bottom*) Genes within the 2 LOD support intervals are depicted, with the exception of panel **C**, which depicts the genes underlying the most likely SNPs.

**Figure S15.**
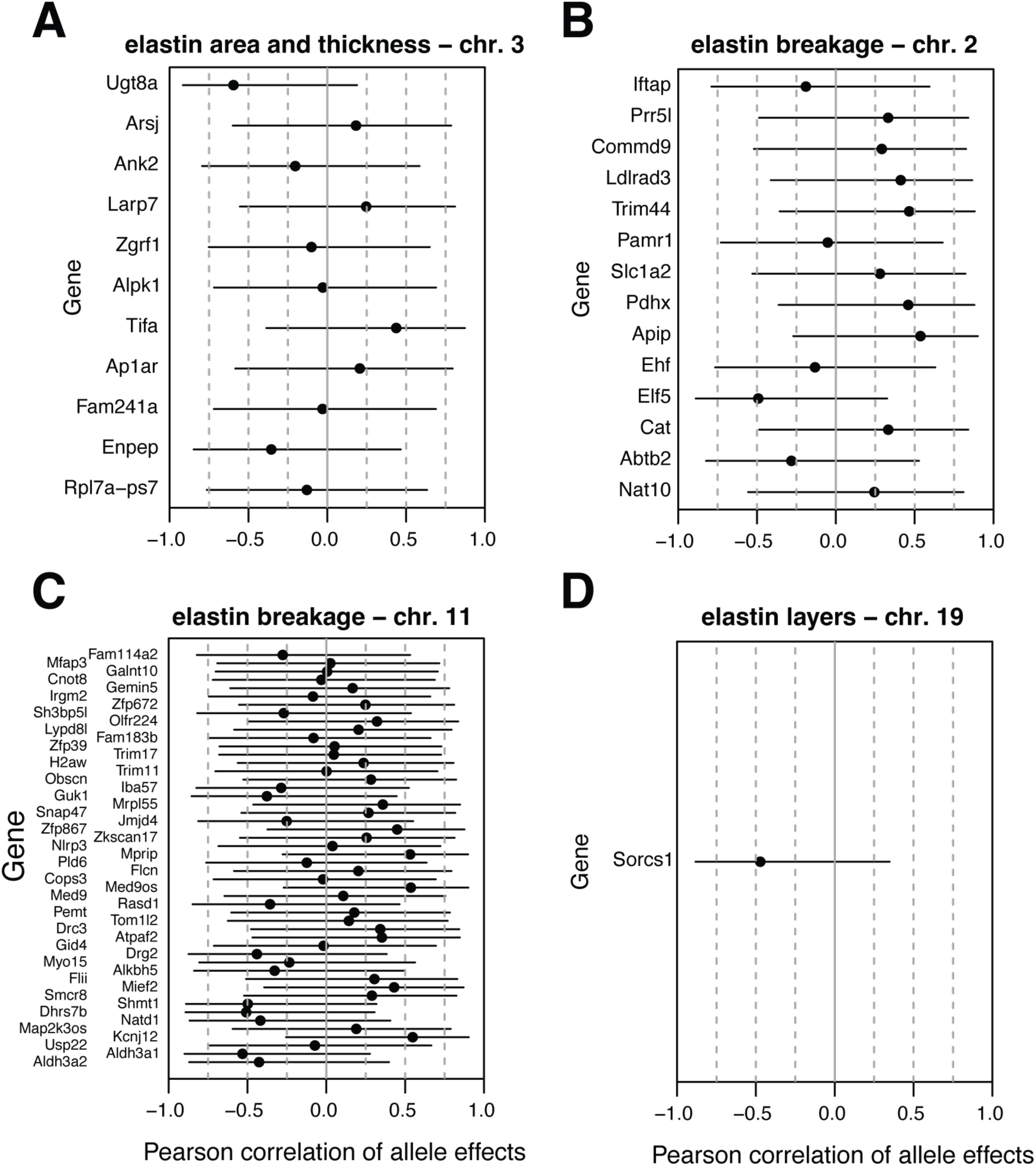
Additional allele effect correlations for additional elastin QTL. **A-D,** Correlations of haplotype effects between the peak markers at QTL influencing elastin phenotypes and *cis*-eQTL in an external DO cardiac tissue transcriptomic dataset. Candidate genes are highlighted with red text.

**Figure S16.**
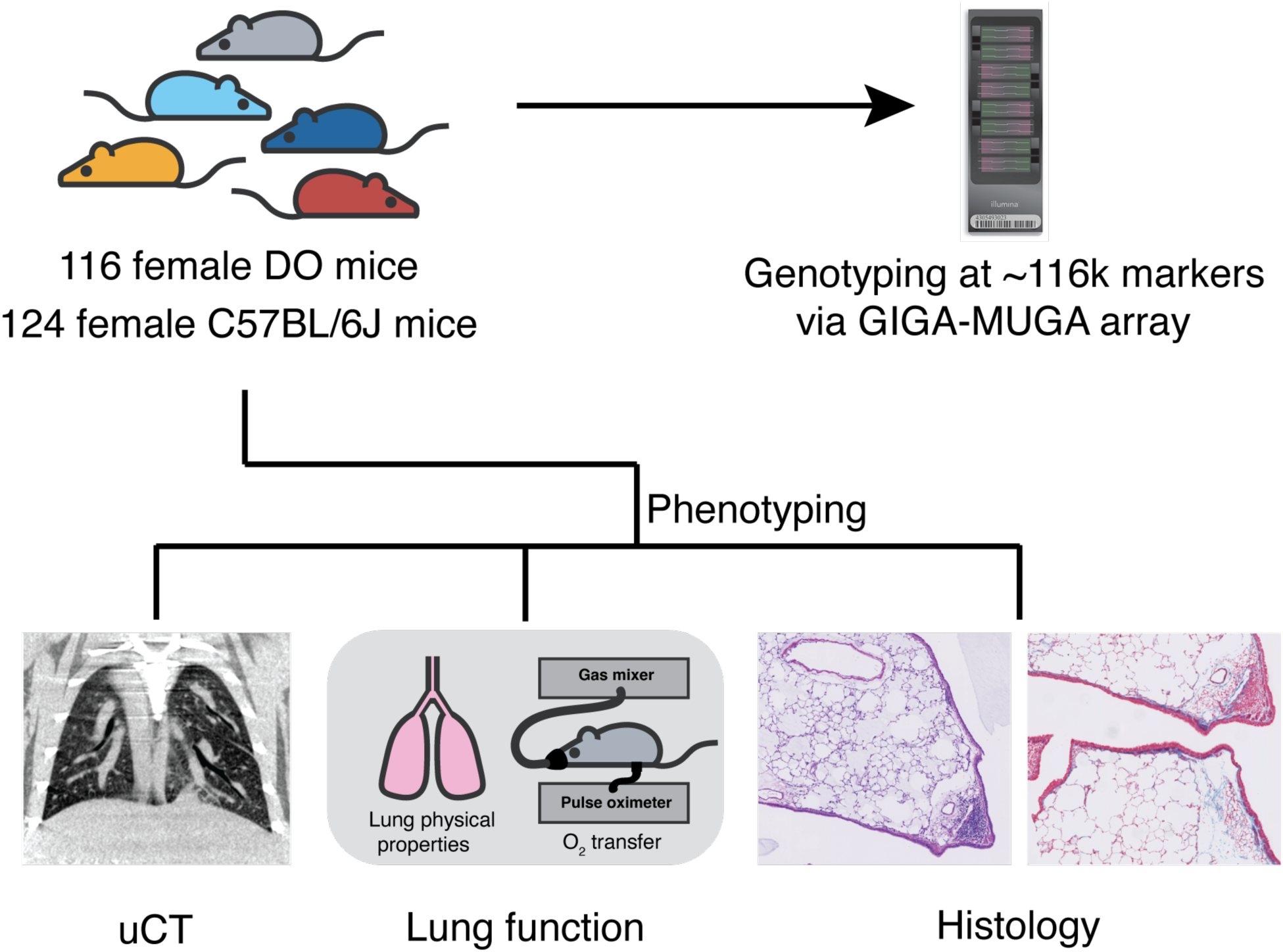
DO mice as a model for lung structure and function. Study design for the assessment of lung properties in mice. 116 DO mice (and 124 C57BL/6J) animals were genotyped and participated in a phenotyping pipeline that included micro-CT scans of the lungs and functional measurements. Upon exit from the study, lung tissue was harvested and fixed, and tissue sections were stained (TC or HE) and imaged.

**Figure S17.**
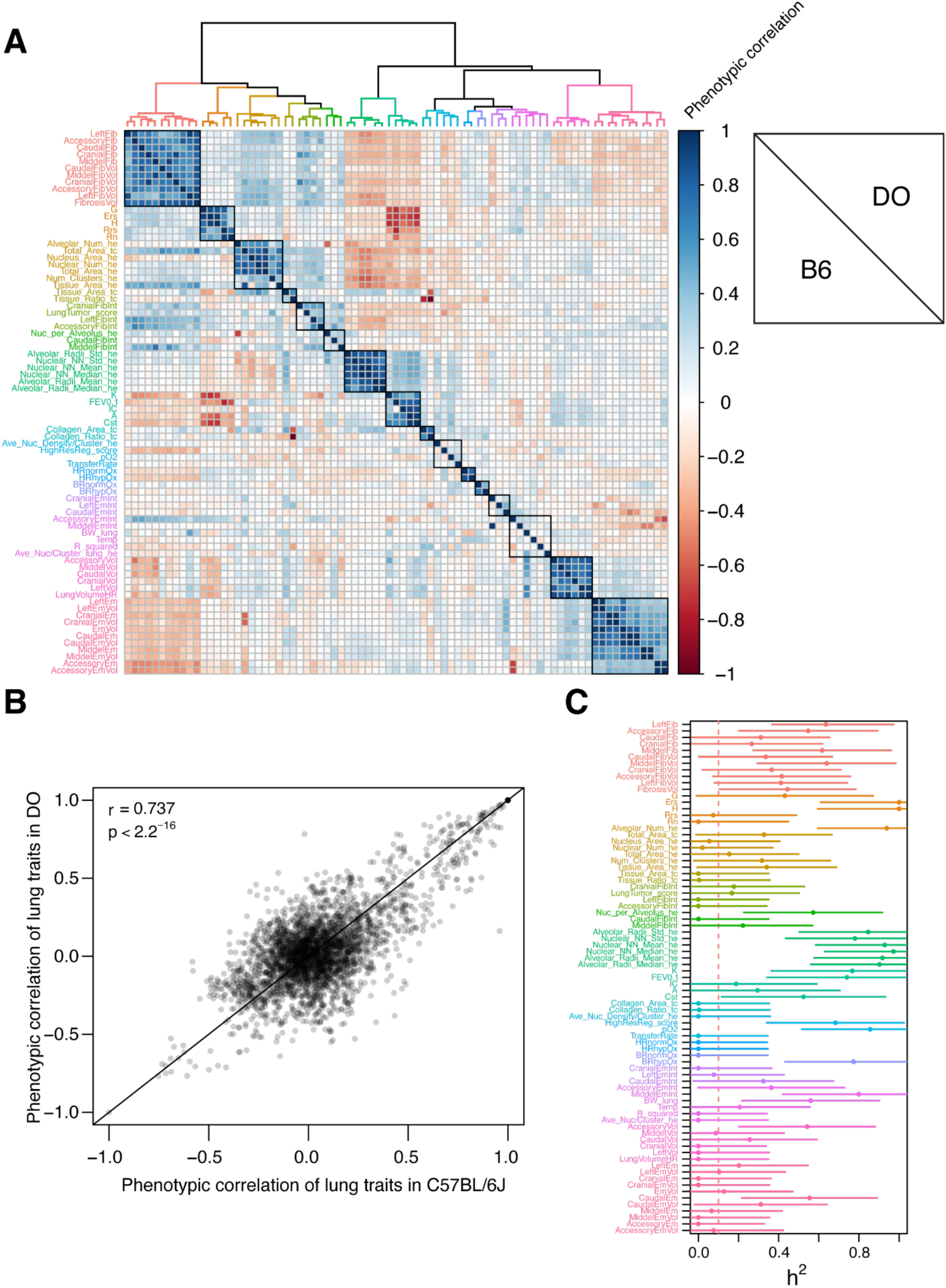
Phenotypic correlations and *h^2^* of lung phenotypes in C57BL/6J and DO mice. **A**, Phenotypic correlations among hierarchically clustered lung phenotypes in DO (*upper triangle*) and C57BL/6J (*lower triangle*) mice. Clusters were determined via silhouette score; phenotype names and the accompanying dendrogram are colored by cluster. **B**, Pairwise phenotypic correlations among pairs of lung traits plotted as a function of their correlation in C57BL/6J mice. **C,** Heritability estimates and standard errors of each lung phenotype prior to filtering. The dashed red line indicates the threshold of *h^2^*< 0.1 at which traits were excluded from downstream analysis.

**Figure S18.**
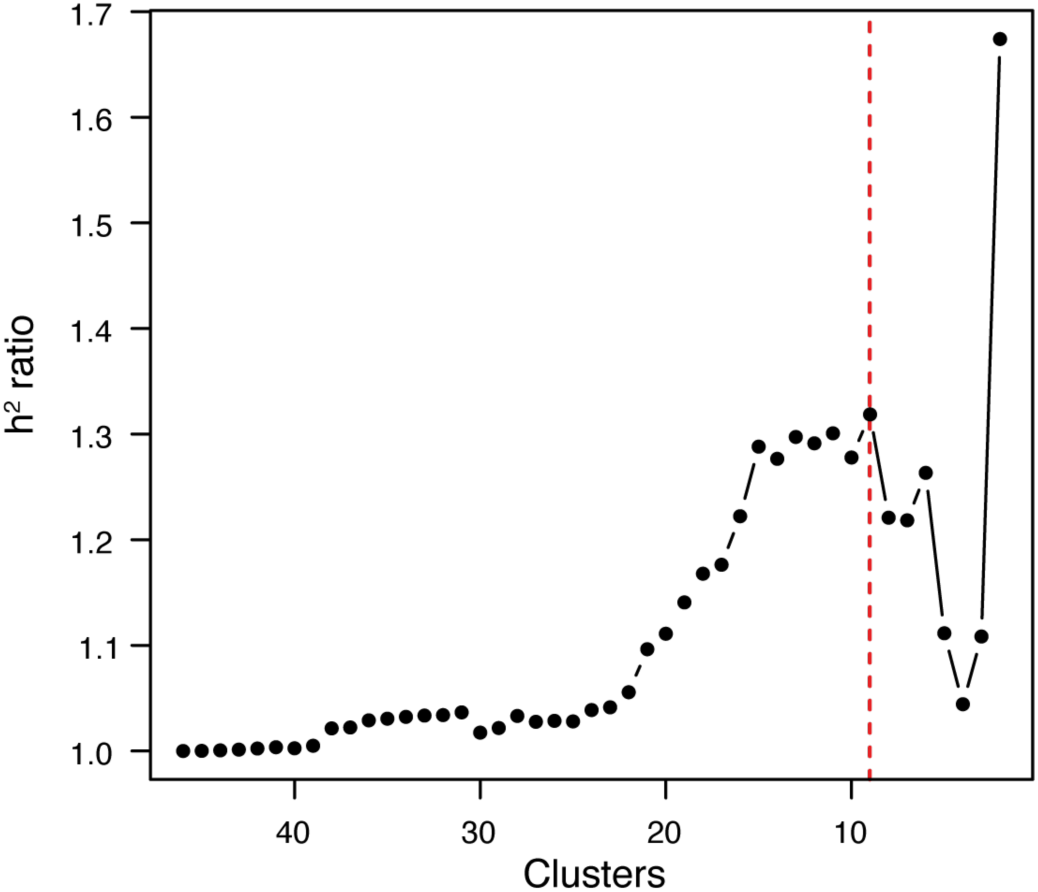
h^2^ ratio by clustering level for lung phenotypes. Plot of mean *h^2^*ratios among clusters of lung meta-traits at different levels of hierarchical clustering (k).

**Figure S19.**
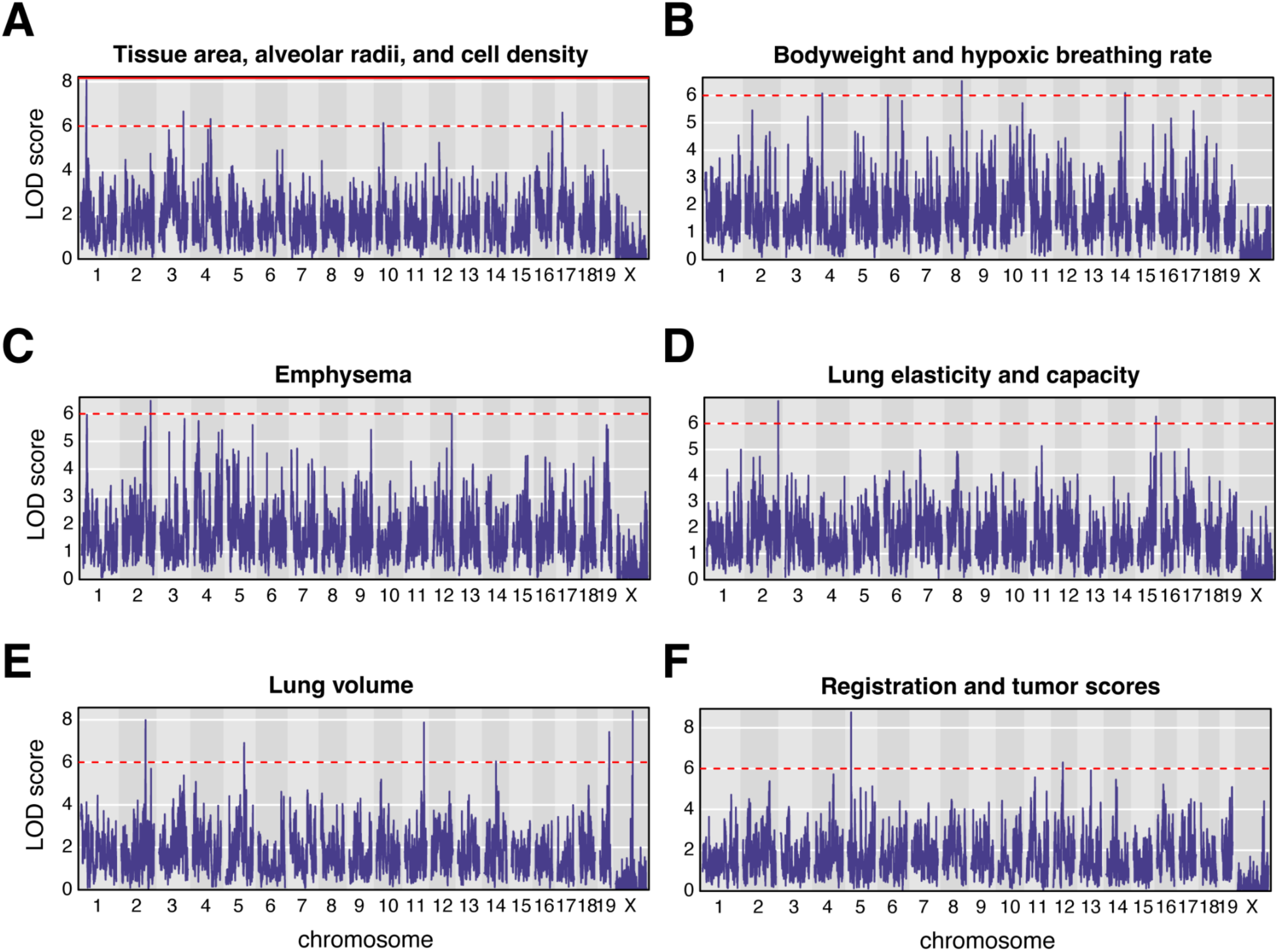
Genetic mapping of additional lung meta-traits. Manhattan plots of additive genome-wide scans for additional meta-traits composed of clustered lung phenotypes: **A**, Tissue area, alveolar radii, and cell density. **B,** Body weight and breathing rate under hypoxic conditions. **C,** Emphysema as measured via micro-CT imaging. **D,** Lung elasticity and capacity. **E,** Lung volume. **F,** Mouse registration and lung tumor scores. Solid red lines indicate a trait-specific and permutation-based genome-wide significance threshold. Dashed red lines indicate an additional threshold of LOD > 6.

**Figure S20.**
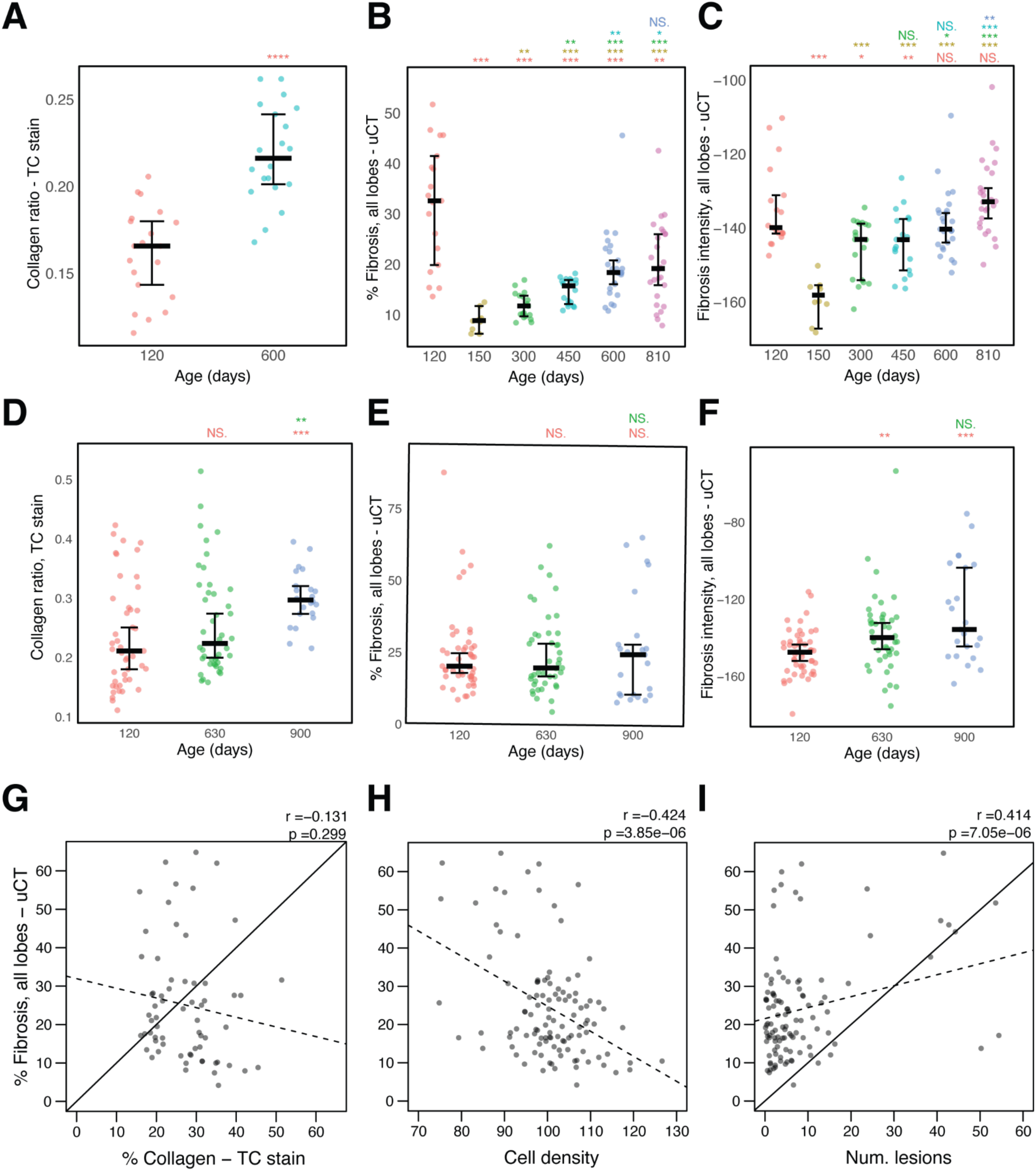
µCT-derived tissue density phenotypes are not correlated with collagen levels and may reflect pulmonary inflammation or tumorigenesis. **A**, Fraction of lung tissue area stained for collagen via trichrome (TC) staining as a function of age in C57BL/6J mice. **B,** Percent area of the lung categorized as “high density” via microCT scan in C57BL/6J mice. **C,** Pixel intensity of “high density” tissue in microCT scans in C57BL/6J mice. **D,** Fraction of lung tissue area stained for collagen via TC staining as a function of age in DO mice. **E,** Percent area of the lung categorized as “high density” via microCT scan in DO mice. **F,** Pixel intensity of “high density” tissue in microCT scans in DO mice. **G-H,** Percent area of the lung categorized as “high density” via microCT scan as a function of (**G**) percent tissue area stained for collagen via TC stain, (**H**) the cellular density in the tissue via H&E (HE) staining, and (**I**) the number of abnormal lesions in the tissue in DO mice. The Pearson correlation coefficients and corresponding p values are reported above each plot. The solid lines are lines of identity and the dashed lines are best-fine lines from linear models.

**Figure S21.**
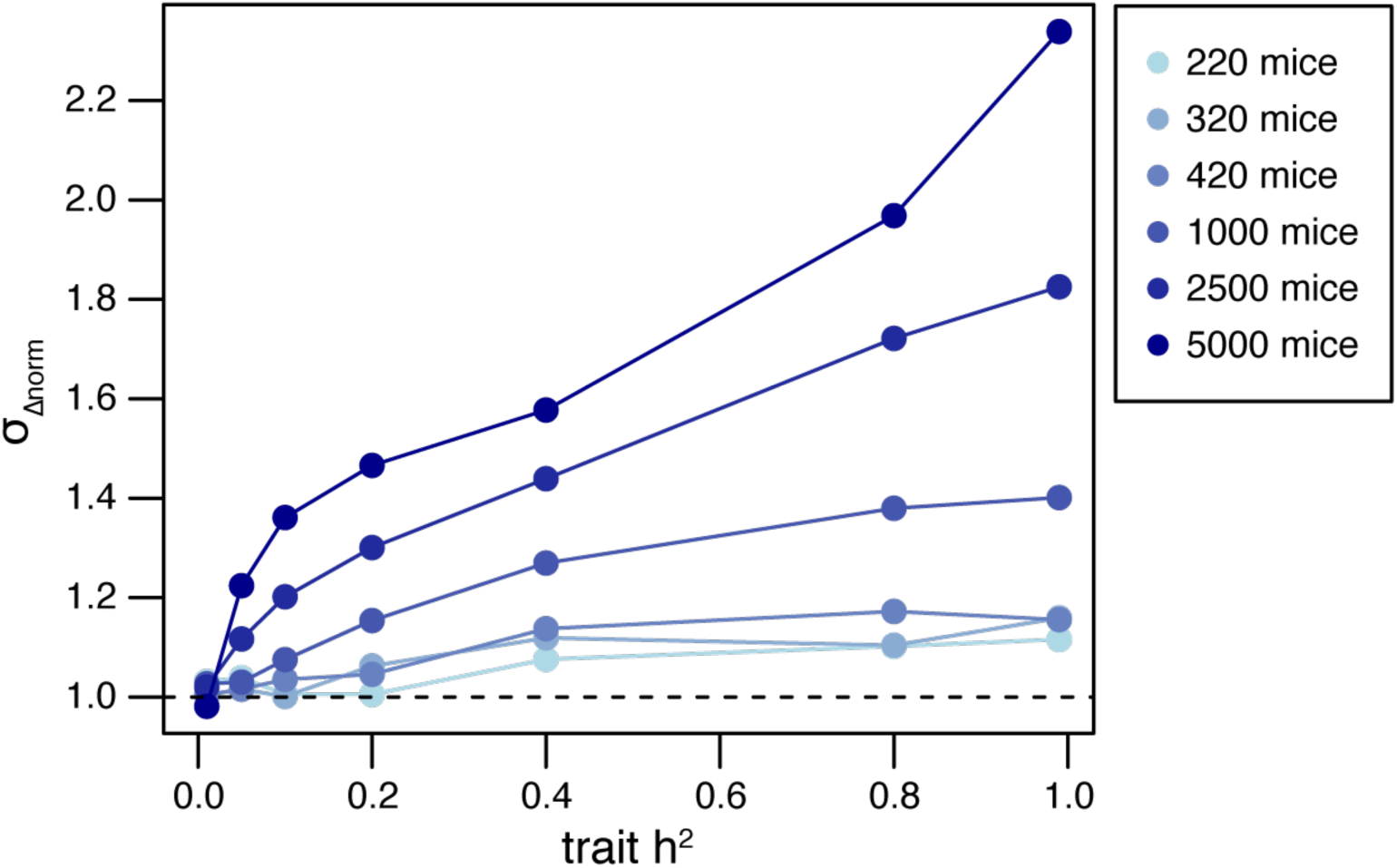
Precision of Haseman-Elston *h*^2^ standard error estimates decreases as a function of *h*^2^ and sample size. Standard deviation of σ_Δ_*_norm_* values for simulated phenotypes across a range of *h^2^* and population sizes. Values less than one indicate that the standard error of *h^2^* for the simulated phenotypes is greater than expected. Values greater than one indicate that the precision of the standard error of *h^2^* estimates is less than expected.

**Figure S22.**
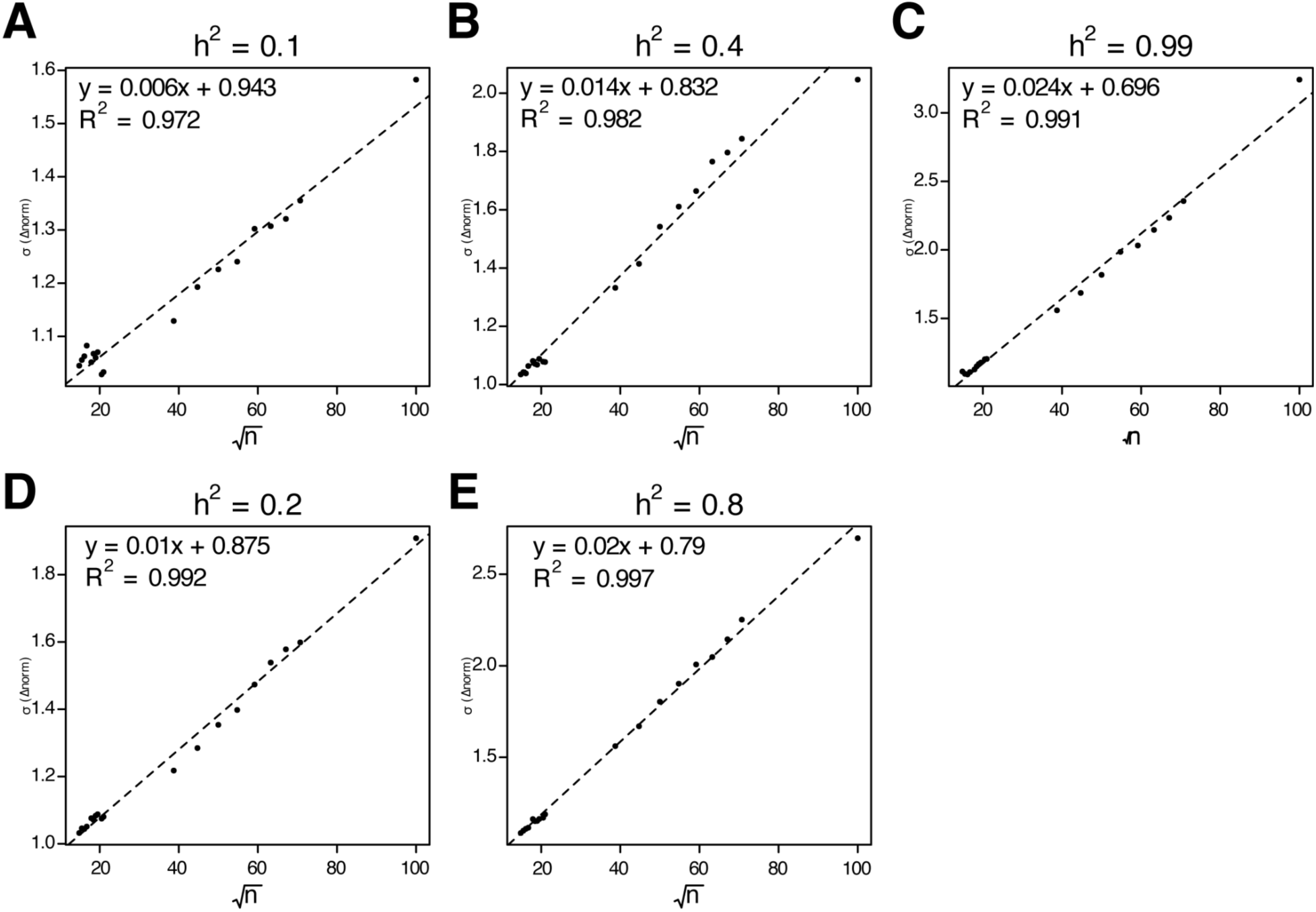
Linear relationship between σ_Δ_*_norm_* and √n of simulated traits. Linear regressions of σ_Δ_*_norm_* values for simulated phenotypes across different population sizes (n) at **A,** *h^2^* = 0.1, **B,** *h^2^* = 0.2, **C,** *h^2^* = 0.4, **D,** *h^2^* = 0.8, **E,** *h^2^* = 0.99. The coefficients and *R*^2^ of each linear regression are reported in the upper left of each panel.

**Figure S23.**
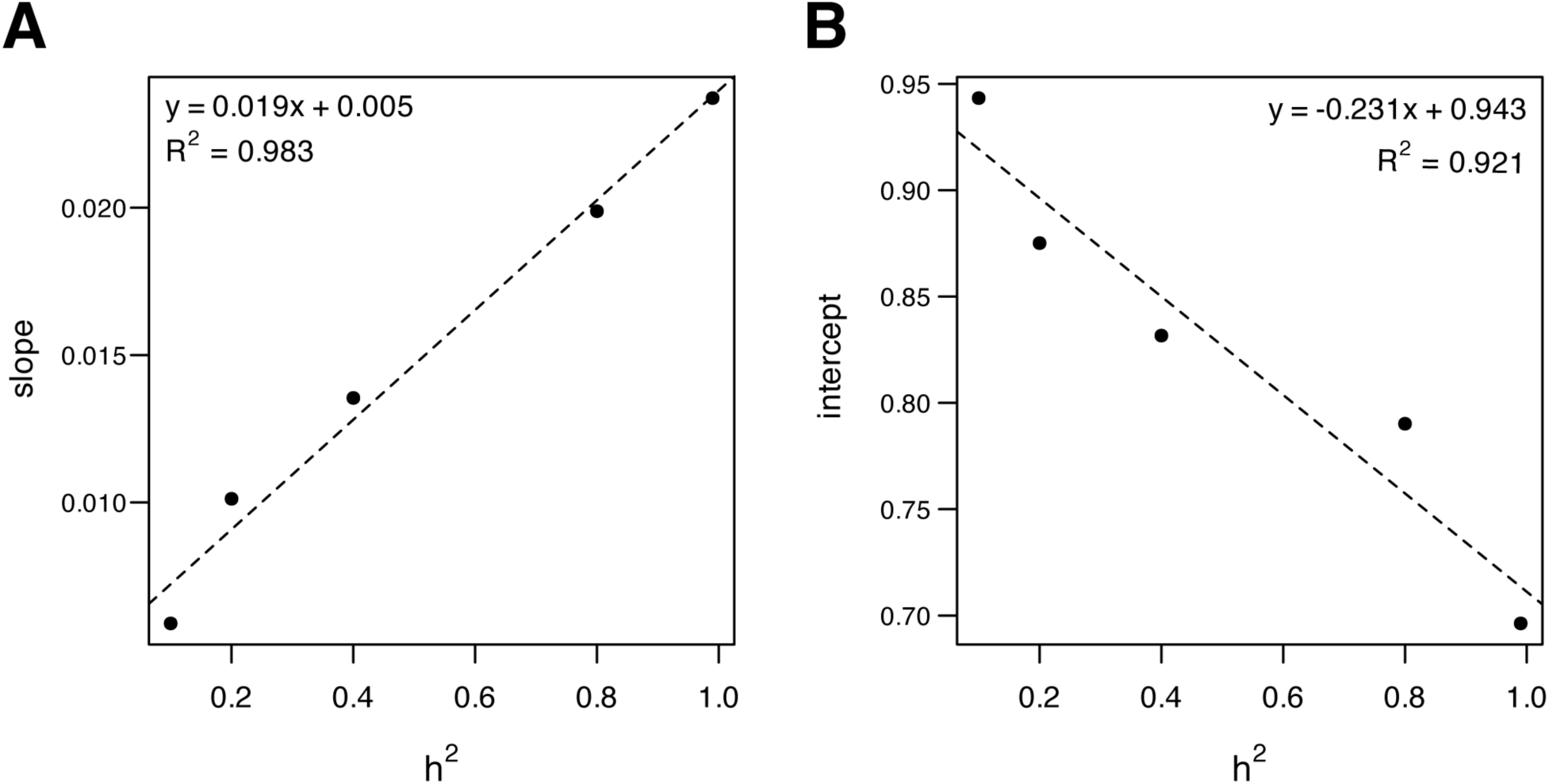
The relationship between coefficients from linear modeling of σ_Δ_*_norm_* and h^2^. The **A,** slope and **B,** intercept terms from linear regressions of σ_Δ_*_norm_* on √n plotted as a function of h^2^, demonstrating how standard error estimates deviate from expected precision as a function of the *h^2^* of the examined phenotype. The coefficients and *R*^2^ of each linear regression are reported in the upper left of each panel.

**Figure S24.**
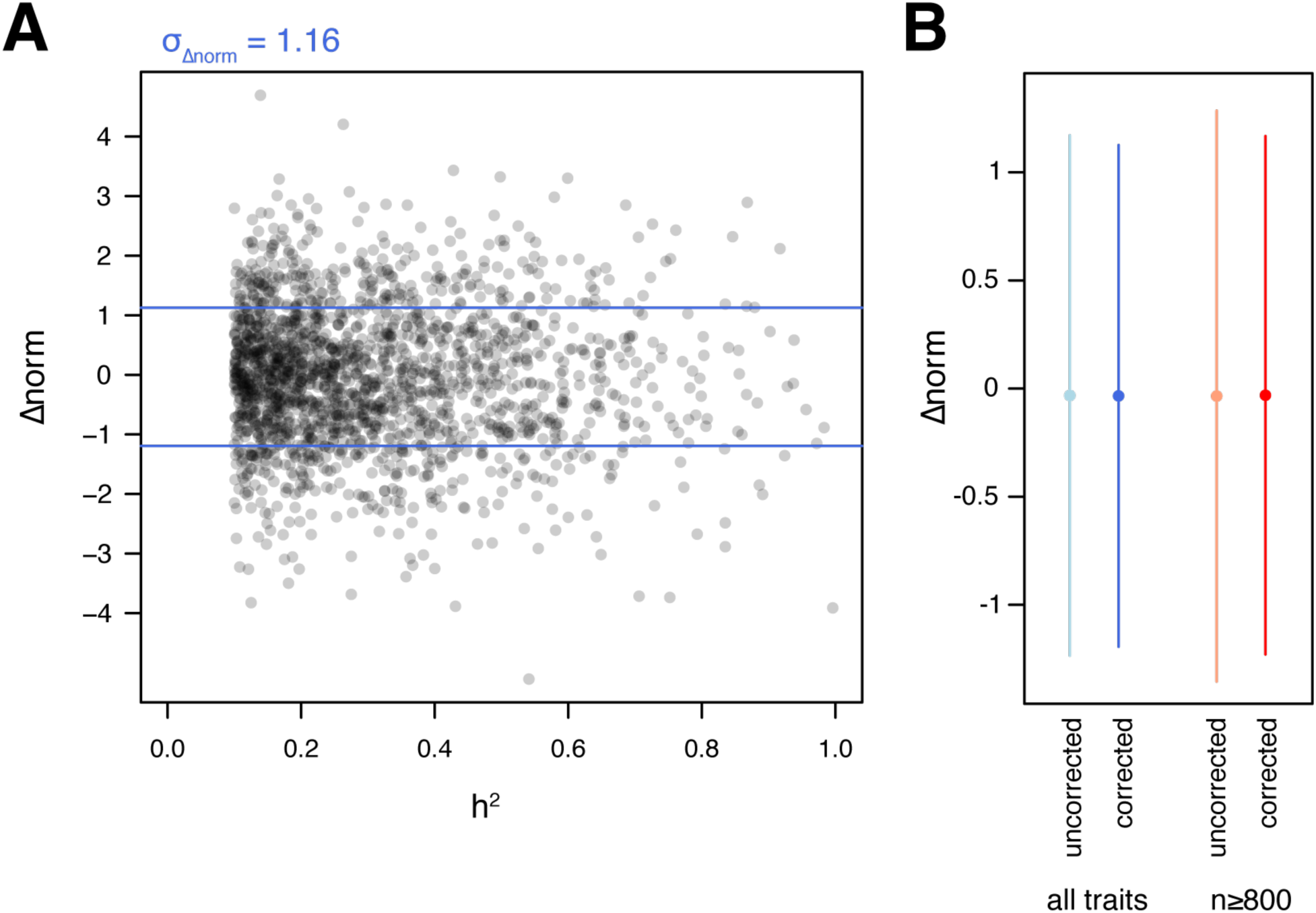
Precision of corrected standard errors from Haseman-Elston h^2^ estimates for DO mouse phenotypes. **A**, *Δnorm* values for 1,898 traits in DO mice using corrected standard error estimates are plotted as a function of trait h^2^. **B,** The mean and standard deviation of *Δnorm* values using corrected and uncorrected standard error estimates for all traits (*left*) and traits with larger sample sizes (*right*).

## SUPPLEMENTARY MATERIALS

**Table S1. Normalized DO mouse phenotypes.** Contains Mouse IDs, covariates used in phenotype processing, and z-score normalized phenotype values. ‘Mouse_ID’ is a unique identifier corresponding to each unique animal and includes study information. ‘Age’ is a covariate corresponding to the age of each animal at the time of phenotyping. For certain animals, different phenotypes were collected at different ages; for these animals, ‘Age’ is recorded as “variable”. ‘Sex’ corresponds to the sex of the animal (‘F’ for female and ‘M’ for male). ‘Diet’ corresponds to the dietary intervention group for each animal, but may also correspond to the number of days on a high fat diet (animals from the Svenson high-fat diet study, denoted as ‘Sven_HFD’) or pharmacological intervention. ‘Genwave’ corresponds to the DO generation of each animal. Additional columns correspond to various z-score normalized phenotypes after correction for the aforementioned covariates. Phenotype names contain a prefix corresponding to the study from which they were collected: ‘attie’ (pancreatic phenotypes from (Keller et al. 2018)), ‘bone’ (bone strength phenotypes from (Al-Barghouthi et al. 2021)), ‘cal_cr’ (DRiDO phenotypes), ‘cal_int’ (unpublished frailty, aorta, and lung phenotypes), ‘catnap’ (metabolic cage data from (Z. Chen et al. 2022)), ‘ches_striatum’ (phenotypes from (Bagley et al. 2022)), ‘lifespan’ (lifespan data from independent lifespan studies analyzed in (Mullis et al. 2025); ‘shock’ and ‘har’ correspond to the Shock and Harrison studies, respectively), ‘pazdro_heart’ (cardiac phenotypes from (Starcher et al. 2021)), and ‘Sven_HFD’ (various phenotypes from (Gatti et al. 2017)). Phenotypes may also contain a suffix corresponding to dietary intervention groups from the DRiDO study: ‘AL’ (*ad libitum*), ‘20’ (20% caloric restriction), ‘40’ (40% caloric restriction), ‘1D’ (one-day intermittent fasting), ‘2D’ (two-day intermittent fasting). Phenotypes from the DRiDO study collected annually or bi-annually will contain timepoint information corresponding to the year of the study in which they were collected (‘Y1’ - ‘Y4’). Biannual measurements are labeled with ‘A’ or ‘B’, corresponding to the first or second annual time point.

**Table S2. Data sources for DO mouse phenotypes.** Contains information on the studies from which phenotypes were obtained. This includes the number of animals used and phenotypes measured, the general focus of each study, the source of the phenotype and genotype data, the genotyping array used, and the prefix assigned to corresponding Mouse_ID and phenotype labels.

**Table S3. Heritability (*h^2^*) of individual traits.** Individual heritability estimates for each of the phenotypes in the dataset, before filtering based on *h^2^*. ‘Trait’ corresponds to one of the phenotypes in **Table S1**. ‘h2’ provides the heritability of the trait as estimated by Haseman-Elston regression. ‘SE’ corresponds to the standard error of the trait. ‘p’ corresponds to the p-value of the *h^2^* estimate. ‘n’ corresponds to the number of animals used in the estimate.

**Table S4. Pairwise genetic correlations (*r_g_*) of individual traits.** Pairwise genetic correlation estimates for each of the phenotypes in the dataset, after filtering based on *h^2^*. Each row contains the following information: the pair of traits (‘Trait1’ and ‘Trait2’, respectively), the *r_g_* estimate (‘rg’), the standard error of *r_g_* (‘SE’), the p value associated with the Haseman-Elston regression (‘rg_p’), the Pearson correlation coefficient of the phenotypes (‘r’), the p value associated with the Pearson correlation coefficient (‘r_p’), the *h^2^* estimates for each trait (‘h2_1’ and ‘h2_2’), and the number of animals in which the traits were measured (‘n_1’ and ‘n_2’).

**Table S5. Hierarchically clustered matrix of genetic correlations (*r_g_*).** Row and column names correspond to the DO phenome after filtering by *h^2^.* This matrix includes DRiDO phenotypes split by diet group.

**Table S6. Hierarchically clustered matrix of Pearson genetic correlations (*r_pg_*).** Row and column names correspond to the DO phenome after filtering by *h^2^.* This matrix includes DRiDO phenotypes split by diet group.

**Table S7. Meta-trait composition.** Lists 1,898 phenotypes, filtered by *h^2^* and excluding diet-specific DRiDO phenotypes. The ‘Cluster’ column specifies the meta-trait each trait comprises.

**Table S8. Meta-traits constructed from clustered individual phenotypes.** Meta-phenotype values for each animal in the dataset. Animal IDs are listed in the ‘Mouse_ID’ column, with each subsequent column corresponding to a meta-trait. IF none of the traits comprising a meta-trait were measured in a particular animal, the trait value for that animal is ‘NA’.

**Table S9. Heritability of meta-traits.** Heritability estimates for each of the meta-phenotypes in the dataset, excluding meta-traits composed of lifespan data from the Ellison study (prefixed with “lifespan_ell”, which were not significantly heritable and excluded from analysis). ‘Trait’ corresponds to a meta-trait, which is composed of individual phenotypes listed in **Table S8**. ‘h2’ provides the heritability of the trait as estimated by Haseman-Elston regression. ‘SE’ corresponds to the standard error of the trait. ‘p’ corresponds to the p-value of the *h^2^* estimate. ‘n’ corresponds to the number of animals used in the estimate.

**Table S10. Genetic correlations among meta-traits.** Pairwise genetic correlation estimates for each of the meta-phenotypes in the dataset, excluding meta-traits composed of lifespan data from the Ellison study (prefixed with “lifespan_ell”, which were not significantly heritable and excluded from analysis). Each row contains the following information: the pair of traits (‘Trait1’ and ‘Trait2’, respectively), the *r_g_* estimate (‘rg’), the standard error of *r_g_* (‘SE’), the p value associated with the Haseman-Elston regression (‘rg_p’), the Pearson correlation coefficient of the phenotypes (‘r’), the p value associated with the Pearson correlation coefficient (‘r_p’), the *h^2^* estimates for each trait (‘h2_1’ and ‘h2_2’), and the number of animals in which the traits were measured (‘n_1’ and ‘n_2’).

**Table S11. Hierarchically clustered matrix of genetic correlations among meta-traits.** Row and column names correspond to the meta-trait after removing two meta-traits assembled from low *h^2^* lifespan estimates.

**Table S12. Hierarchically clustered matrix of Pearson genetic correlations among meta-traits.** Row and column names correspond to the meta-trait after removing two meta-traits assembled from low *h^2^* lifespan estimates.

**Table S13. QTL detected in meta-traits at permutation-based significance levels.** For each QTL, the following information is recorded: the meta-trait associated with the locus (‘Trait’), the chromosome (‘Chr’) and position information (‘Pos’, in megabases), the LOD score (‘LOD’), and the bounds of the 2LOD support interval (‘CI_start’ and ‘CI-end’).

**Table S14. QTL detected in meta-traits at LOD ≥ 6.** For each QTL, the following information is recorded: the meta-trait associated with the locus (‘Trait’), the chromosome (‘Chr’) and position information (‘Pos’, in megabases), the LOD score (‘LOD’), and the bounds of the 2LOD support interval (‘CI_start’ and ‘CI-end’).

**Table S15. Frailty meta-trait composition.** Lists each of the frailty phenotypes after filtering by *h^2^*. The ‘Cluster’ column specifies the meta-trait each trait comprises.

**Table S16. Aorta meta-trait composition.** Lists each of the aorta phenotypes after filtering by *h^2^*. The ‘Cluster’ column specifies the meta-trait each trait comprises. Trait suffixes correspond to the type of stain from which the measurement was derived: H&E (‘*he’),* trichrome (‘_tc’), or Verhoeff-Van Gieson (VVG; ‘_vvg’).

**Table S17. QTL detected in aorta mega-analysis at LOD ≥ 6.** For each QTL, the following information is recorded: the meta-trait associated with the locus (‘Trait’), the chromosome (‘Chr’) and position information (‘Pos’, in megabases), the LOD score (‘LOD’), and the bounds of the 2LOD support interval (‘CI_start’ and ‘CI-end’).

**Table S18. Lung meta-trait composition.** Lists each of the frailty phenotypes after filtering by *h^2^*. The ‘Cluster’ column specifies the meta-trait each trait comprises. As in **Table S16**, trait suffixes correspond to the type of stain from which a phenotypic measurement was derived, when present.

**Table S19. QTL detected in lung mega-analysis at LOD ≥ 6.** For each QTL, the following information is recorded: the meta-trait associated with the locus (‘Trait’), the chromosome (‘Chr’) and position information (‘Pos’, in megabases), the LOD score (‘LOD’), and the bounds of the 2LOD support interval (‘CI_start’ and ‘CI-end’).

**Data S1. Allele probabilities for DO mice.** Haplotype probability information for the animals included in this study at ∼69k genetic pseudomarkers. A list of three-dimensional matrices, with dimensions corresponding to animal, marker, and founder allele probability.

**Data S2. Kinship data for DO mice.** A list of 20 kinship matrices generated via the ‘qtl2’ packages in R. Each matrix corresponds to a chromosome, with the 20^th^ matrix corresponding to the X chromosome.

**Data S3. Pseudomarkers used to interpolate genotype data across study.**

**Data S4. Genome-wide scan data for meta-traits.**

**Data S5. Variant association mapping for each phenome-wide meta-trait QTL, and lists of genome annotations within the confidence intervals of the QTL.**

**Data S6. Genome-wide scan data for aorta meta-traits.**

**Data S7. Variant association mapping for each aorta meta-trait QTL, and lists of genome annotations within the confidence intervals of the QTL.**

**Data S8. Genome-wide scan data for lung meta-traits.**

**Data S9. Lists of genome annotations within lung meta-trait QTL confidence intervals.**

